# Enhanced extracellular matrix production provides protection to cell wall-deficient *Escherichia coli*

**DOI:** 10.1101/2024.11.06.622226

**Authors:** Marjolein E. Crooijmans, Joost J. Willemse, Johannes H. de Winde, Dennis Claessen

## Abstract

*Escherichia coli*-induced recurrent urinary tract infections (rUTIs) present a complicated challenge within the medical field. Most first-line antibiotic treatments primarily target cell-wall synthesis, which can lead to the formation of cell wall-deficient cells. To investigate how such cells can sustain, we obtained an *E. coli* strain capable of efficiently proliferating without its cell wall. One of the mutations lead to enhanced expression of *rcsA*, encoding an important regulator involved in responding to cell envelope stress. RNA sequencing demonstrated an upregulation of genes associated with the production of extracellular matrix components, and this increased extracellular matrix production was confirmed using various imaging techniques. Remarkably, a subsequent long-term evolution experiment on this strain revealed a further augmentation in extracellular matrix production, coinciding with an enhanced ability to withstand harsh conditions. These findings demonstrate how *E. coli* adapts to loss of its cell wall and that an increased synthesis of matrix constituents can compensate for the protective properties of the cell wall.

**Significance:** The cell envelope is crucial for the protection of *E. coli*. During bacterial infections such as urinary tract infections, antibiotics that disrupt cell wall synthesis are commonly prescribed. However, this can stimulate the formation of wall-deficient bacteria that are still able to proliferate despite the presence of these drugs. Our findings reveal that loss of the cell wall in *E. coli* increases the production of extracellular matrix, a mechanism found in other unicellular organism too. This adaptation allows the bacteria to maintain their structural integrity and survive, highlighting a potential challenge in the successful treatment of bacterial infections.

## Introduction

The diderm cell envelope of *E. coli* provides an important barrier against environmental stressors (Silhavy, Kahne et al. 2010). This structure is comprised of multiple protective layers. Surrounding the cytoplasm is the inner membrane, predominantly composed of glycerophospholipids. The outer membrane primarily contains an additional lipopolysaccharide layer on the outer leaflet (Fivenson, Rohs et al. 2023). Inserted between these membranes is a thin layer of peptidoglycan, consisting of crosslinked repeating disaccharide strands (Egan, Errington et al. 2020). The discovery of penicillin in 1928 by Alexander Fleming, provided a direct way to disrupt the biosynthesis of peptidoglycan. This class of antibiotics, known as beta-lactams, block peptidoglycan binding proteins from crosslinking the nascent peptidoglycan strands, resulting in cell lysis (in most cases). While a lot of research has been focused on the synthesis of nascent cell wall, the survivability of cells lacking the cell wall remains relatively underexplored.

Despite the great successes obtained with beta-lactams, we are now aware that exposure to these cell wall-targeting agents can lead to the formation of cell-wall deficient cells in many bacteria (Errington, Mickiewicz et al. 2016, Errington 2017, Claessen and Errington 2019, Lazenby, Li et al. 2022). This phenomenon has been observed in *Staphylococcus aureus* (Mercier, Kawai et al. 2014), *Bacillus subtilis (Leaver, Dominguez-Cuevas et al. 2009)*, *Pseudomonas aeruginosa* (Spalding, Shirgill et al. 2022) and *E. coli* (Joseleau-Petit, Liébart et al. 2007, Petrovic Fabijan, Martinez-Martin et al. 2022). These types of cells have been directly linked to increased pathogenicity in the past (Conner, Coleman et al. 1968, Clasener 1972).

Loss of the cell wall does not automatically lead to efficient proliferation of such cells (Ramijan, Ultee et al. 2018, Tabata, Sogo et al. 2019). Typically, efficient proliferation without a cell wall occurs only when such cells acquire specific mutations. These mutations differ among organisms and between different L-forms (Liu, Zhang et al. 2023), but often lead to an increase in synthesis of membrane components (Leaver, Dominguez-Cuevas et al. 2009) and a reduction in oxidative damage (Kawai, Mercier et al. 2015). A mutant screening study performed with *E. coli* found 24 essential genes for formation of L-form colonies on agar (Glover, Yang et al. 2009). These genes belonged to pathways involved in, among others, cell envelope stress, DNA repair and outer membrane synthesis. Surprisingly, at least half of these genes belong to the Rcs (regulator of capsule synthesis) phosphorelay response and colanic acid biosynthesis pathways. This highlights the essential role of a functional envelope stress response and subsequent extracellular matrix modifications in response to this type of stress (Cambre, Zimmermann et al. 2015).

The extracellular matrix of *E. coli* is a network of macromolecules on the outside of the cell that includes protein assemblies, polysaccharides, metabolites, lipids and nucleic acids, in which the biofilm cells are embedded. The most abundant components are curli and cellulose (Jeffries, Fuller et al. 2019). Curli proteins are required for building an intricate network of amyloid fibers that facilitate cell-cell adhesion and biofilm formation. Cellulose, long chains of β-(1,4)-coupled glucose residues, provides structural stability and promotes intercellular interactions (Zogaj, Nimtz et al. 2001). The bacterial matrix also consists of two other polysaccharides, poly-N-acetylglucosamine (PGA), and colanic acid. PGA is an essential adhesin in biofilm formation (Wang, Preston et al. 2004). Colanic acid consists of repeating units of D-galactose, L-fucose, D-glucose, and D-glucuronic acid, forming a negatively-charged exopolysaccharide loosely attached to the outer membrane. Together, these components form an extracellular harness that provides robust resistance of walled bacteria to antibiotics, phages, drought and other stressors (Limoli, Jones et al. 2015). Nevertheless, insights into how these matrix components contribute to the lifestyle of cell wall-deficient bacteria remain unclear.

In this study, we generated a novel *E. coli* L-form strain and characterized the genetic alterations required to induce efficient wall-deficient growth. One of the mutations led to increased synthesis of extracellular matrix components, resulting in cell wall-deficient cells being surrounded by a dense layer of cellulose and amyloid fibers. A long-term evolution experiment on this new L-form strain further enhanced extracellular matrix production, coinciding with an increased robustness to the cells. Altogether, these results show that *E. coli* may elicit a protective system when cells become cell wall-deficient whereby the extracellular matrix is able to substitute for the protective properties of the cell wall.

## Results

### Characterization of a novel *E. coli* L-form strain

To study wall-deficient proliferation in greater dept, we created an unstable L-form of *E. coli* K-12 MG1655 by exposing spheroplasts to mutagenesis, see Fig. 1A (Crooijmans et al., *manuscript under review*). The resulting strain, hereinafter called L-form^O^, efficiently proliferates without its cell wall in the presence of penicillin on high sucrose-containing medium (LPB) (Fig. 1B, Supplementary Video 1). Cell size measurements indicated that L-forms were considerably smaller than spheroplasts (Supplementary Fig. 1, Supplementary Table 1), and mostly contain DNA (Supplementary Fig. 2). In line with earlier findings, we confirmed that wild-type *E. coli* cells can also proliferate without their cell wall, but only when cells are growing under a thin layer of agar and are exposed to penicillin (Fig. 1B, Supplementary Video 2) (Wu, Lee et al. 2020). Without penicillin, the wild-type cells remain their rod-shaped morphology when growing under the agar pad (Supplementary Video 3). In the absence of penicillin, the strain reverts to its original rod-shaped morphology, as seen in Supplementary Video 4. This phenomenon is also seen when spheroplasts are grown in the absence of penicillin (Supplementary Video 5). On high sucrose containing LPMA plates supplemented with penicillin, both L-form^0^ and revertant^0^ colonies have a mucoid and glossy morphology, contrary to the parental wild type (Fig. 1C).

**Fig. 1.**
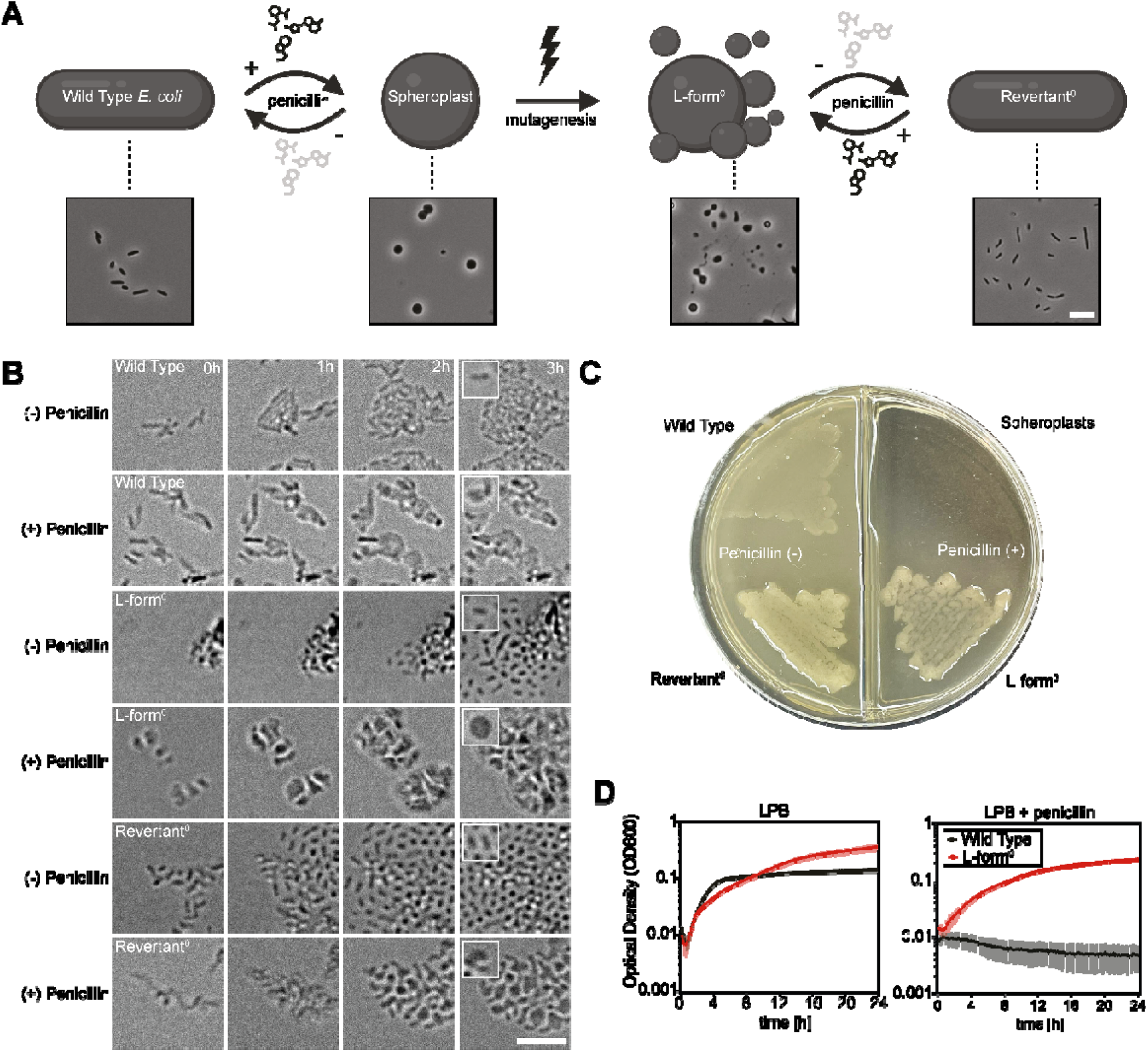
Penicillin- and UV-induced morphological changes in *E. coli.* (A) Schematic overview indicating the transformation process from wild type *E. coli* to spheroplasts, the L-form^0^ strain and its revertant^0^. Overnight grown cells were imaged using transmission light microscopy. The schematic overview was created using Biorender. (B) Timelapse images showing the growth of the wild-type strain cells (U66-GFP), the L-form^0^ strain and its revertant^0^ in the presence and absence of penicillin. Images taken with a Lionheart microscope. (C) Colony morphology of the wild-type, L-form^0^ and revertant^0^ strains grown overnight on LPMA with and without penicillin. Colonies of the L-form^0^ and revertant^0^ strain have a mucoid morphology, while those of the wild-type were non-mucoid. The spheroplasts are unable to grow on agar. (D) Growth curves of the wild type (U66-GFP, black) and L-form (red) strains in LPB medium with and without penicillin. The data points and error bars represent the mean and standard deviation across replicates (n=3). The scalebar represents 10 µm.

In the absence of penicillin, the L-form^0^ cells start to revert to the walled state within two to three hours (Supplementary Videos 2 and 5), corresponding to three and four new generations. The L-form^0^ colony forming units (CFU) count is estimated at the total of 6.7 x 10^6^ CFU ml^−1^ (OD_600_=0.1). Additionally, the L-form^0^ strain was able to grow to a higher optical density in LPB media compared to the wild type (Fig. 1D), indicating that the strain is optimized to grow in high osmotic environments.

### Identification and characterization of mutations associated with efficient *E. coli* L-form growth

To gain insight into the mutations underlying efficient proliferation without a cell wall, we sequenced the genome of the L-form strain and compared this to the parental strain (Crooijmans et al., *manuscript under review*). Comparative analysis identified five mutations, which were confirmed with PCR analysis (Supplementary Fig. 3). Three mutations were associated with genes that had not been associated with L-forms before: a C-to-A transversion in *ykgH,* a 17bp insertion in *slt* (Betzner, Ferreira et al. 1990, Korsak, Liebscher et al. 2005), and a G-to-A substitution in *yjbH* (Fig. 2A, Table 1). YkgH is predicted to be an inner membrane protein (Kim, Lee et al. 2013), *slt* is a well-studied murein transglycosylase, and *yjbH* encodes a putative lipoprotein that is expected to create beta-barrels in the outer membrane, potentially facilitating secretion of extracellular polymeric substances (EPS) (Ferrières, Aslam et al. 2007). In addition to these mutations, we also identified a frameshift mutation in the coding sequence of *ndh*, which encodes the type II NADH:quinone oxidoreductase and leading to a truncated version of the enzyme (Fig. 2A). The absence of this enzyme was previously found to promote proliferation of *Bacillus subtilis* L-forms by reducing the levels of reactive oxygen species (ROS), which are detrimental to L-form cells (Kawai, Mercier et al. 2015). Indeed, this mutation in Ndh also drastically reduced ROS levels in the newly obtained *E. coli* L-form and revertant strains, as shown by flow cytometry and microscopy imaging (Supplementary Fig. 4, Supplementary Table 2). When the cells were grown in LPB medium containing penicillin, we detected a 52% decrease in ROS in the L-form compared to the wild-type cells. Similar results were found when the revertant was grown in LB medium, although the reduction was only 22%.

**Fig. 2.**
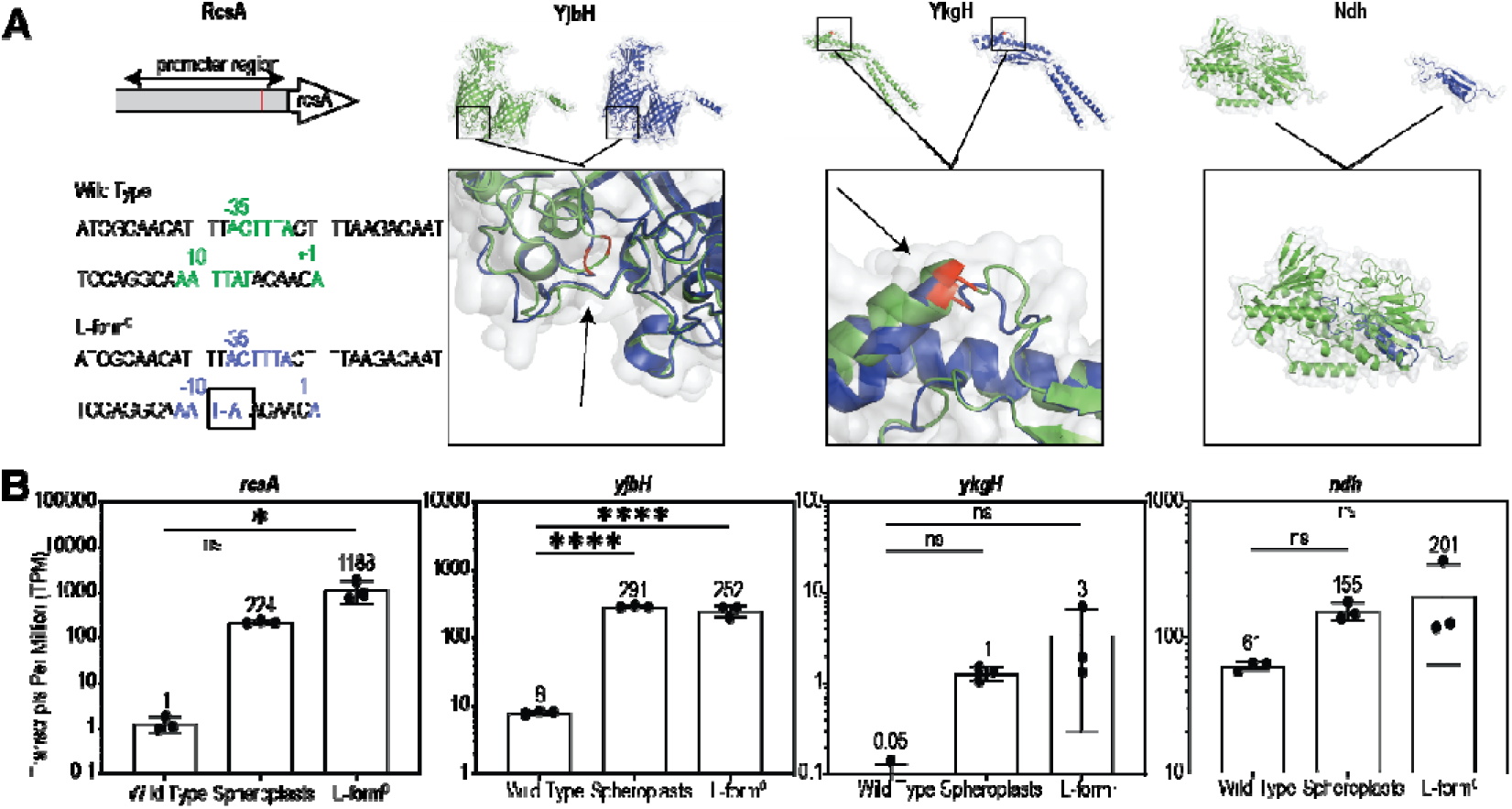
Genomic and transcriptomic analysis of *E. coli* L-form^0^ cells compared to those of the wild-type strain and spheroplasts. (A) Representation of the four genomic variations found in the L-form^0^ strain compared to the parental wild-type strain. Location of the nucleotide deletion in the promoter region of *rcsA* (left panel) and visualization of protein structure changes resulting from the deletions in *ndh* and two SNPs in *yjbH and ykgH,* as predicted by AlphaFold 2 software (right three panels). The wild-type form of the protein in shown in green, while the mutated form of the protein in the L-form^0^ strain is shown in blue. Arrows and red segments indicated the changes in the proteins. (B) RNA expression levels of *rcsA*, *yjbH*, *ykgH* and *ndh* in cells of the wild-type strain, spheroplasts and those of the L-form^0^ strain. The P values were calculated using ANOVA following Dunnett’s multiple comparisons test.

**Table 1.**
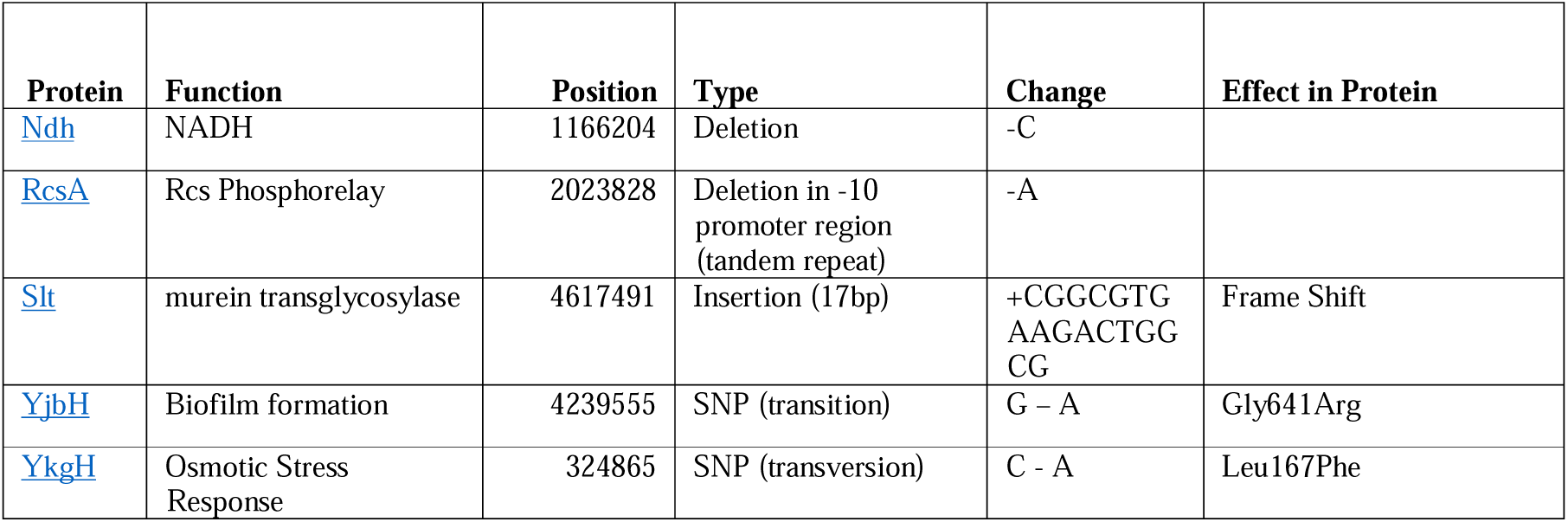
| List of mutations found in the L-form^0^ strain. Position reflects the genomic location. Function is function of affected protein.

The fifth mutation was a nucleotide deletion in the promoter region of *rcsA* (Sledjeski and Gottesman 1995), encoding a transcription regulator of the Rcs phosphorelay response pathway (Supplementary Fig. 5A). This outer membrane and peptidoglycan stress response system is known to be activated during exposure to penicillin treatment (Meng, Young et al. 2021). RcsA, also known as CpsR, is a self-inducer (Clarke 2010) that acts as a heterodimer combined with RcsB (Meng, Young et al. 2021).

To gain a better understanding of the consequences of this mutation, we performed RNA sequencing. When comparing the L-form strain to its parent, we found that the levels of *rcsA* were strongly upregulated (Fig. 2B). We observed a 906-fold increase in *rcsA* expression in the L-form compared to the parental strain, compared to a 5-fold increase in the spheroplasts (Supplementary Table 3). This upregulation of RcsA during cell wall stress is in line with several other studies (Laubacher and Ades 2008, Callewaert, Vanoirbeek et al. 2009, Ranjit and Young 2013, Mohiuddin, Massahi et al. 2022).

The dramatically increased expression of *rcsA* suggested an important beneficiary role for the L-form strain. Indeed, we noticed that a mutant lacking *rcsA* was unable to grow without its cell wall (Supplementary Fig. 5B). The inability to proliferate was also observed in the Δ*rcsB*, Δ*rcsC*, Δ*rcsD,* and Δ*rcsF* mutants from the KEIO collection (Baba, Ara et al. 2006) treated with penicillin, which all lysed within 10 hours (Supplementary Fig. 5B). Together, these results indeed indicate that RcsA constitutes an important stress response regulator in *E. coli*, which plays a critical role for enabling L-form growth.

### Augmented extracellular matrix production during L-form growth

RNA sequencing results offered additional insights into genes essential for growth in the absence of a cell wall (Fig. 3A). A list of all the uniquely expressed genes can be found in Supplementary Data 1. The most down-regulated genes in the L-form cells were involved in synthesizing hydrogenases (eg. HyaB, HyaC, HyaD, HyaF, see Fig. 3B, Supplementary Data 2-3), while the most upregulated genes encode proteins involved in extracellular polysaccharide synthesis. More specifically, the biosynthesis pathway for colanic acid was highly upregulated in the L-form^0^ strain, with the putative polysaccharide export protein Wza showing a more than 3,600-fold increase (Supplementary Data 4, Supplementary Fig. 6). The expression levels of known regulators of other extracellular matrix components were subsequently checked to see if other extracellular matrix components were also upregulated (Supplementary Fig. 6, Supplementary Table 3). Interestingly, in addition to the highly upregulated *rcsA* regulator for colanic acid synthesis, the expression of *csgD*, the curli and cellulose regulator was also significantly upregulated, being 31-fold higher expressed than in the parental wild-type. Several direct downstream targets of CsgD, such as major curlin subunit CsgA and minor curlin subunit CsgB were also upregulated (3-fold and 5-fold increase respectively, Supplementary Fig. 6, Supplementary Data 4). However, the expression of *nhaR*, the regulator of PGA synthesis was not upregulated in the L-form^0^ mutant (Supplementary Fig. 7).

**Fig. 3.**
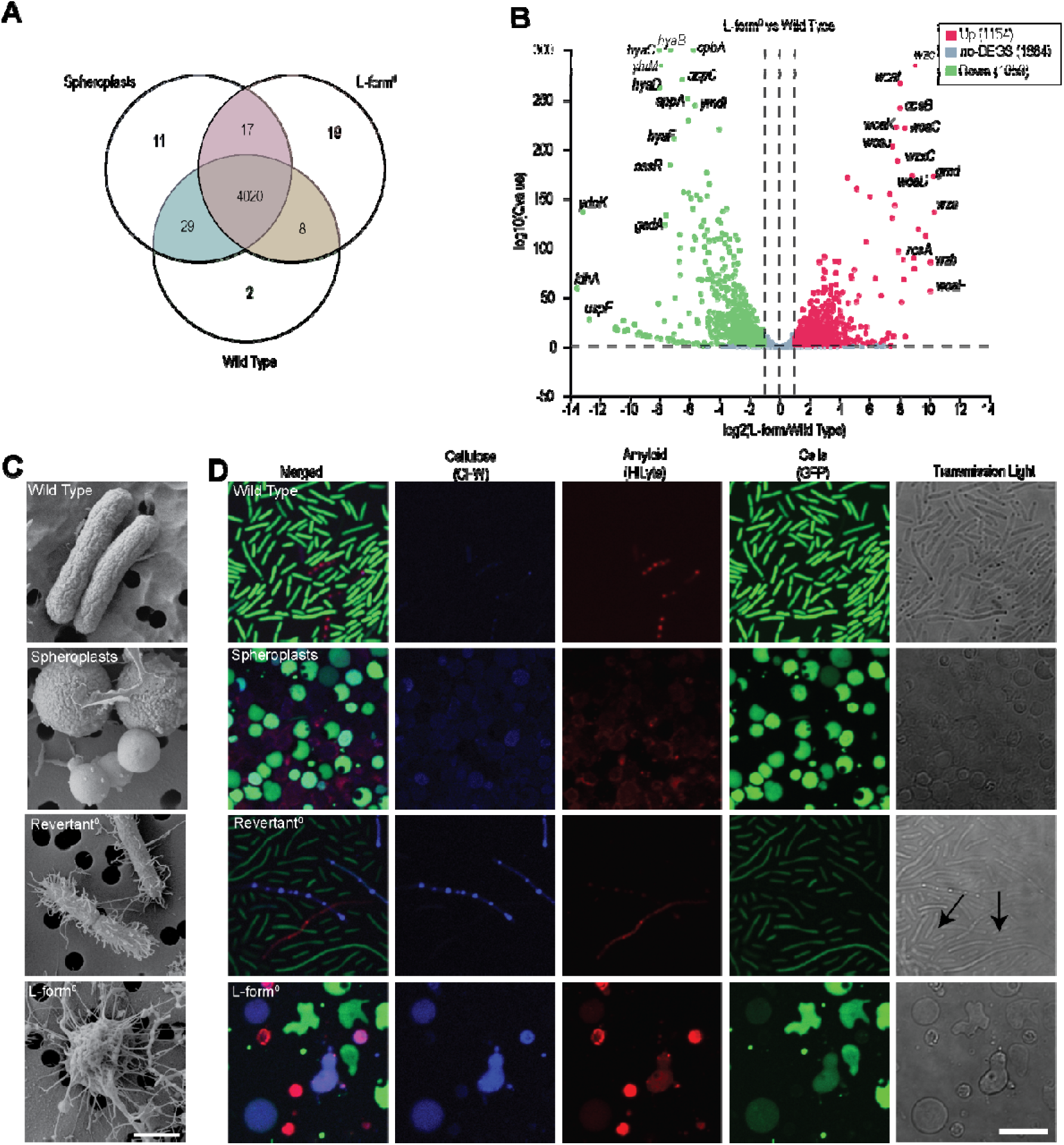
Enhanced extracellular matrix production is associated with L-form growth. **(A)** Venn diagram comparing gene expression among wild-type, L-form^0^, and spheroplast cells in LPB medium. The uniquely expressed genes can be found in Supplementary Data 1. (B) Volcano plot representing the genes that are significantly different expressed in L-form cells compared to those of the wild-type grown in LPB medium. The x-axis represents the log_2_ of the fold-change of the average reads per gene. Genes indicated in green are down-regulated, while those in red are up-regulated. The 150 most significant up- and downregulated genes can be found in Supplementary Data 2-3. (C) SEM images of the extracellular matrices in fixated wild-type, spheroplasts, revertant^0^ and L-form^0^ cells. The scalebar represents 1 µm. (D) Fluorescence microscopy images depicting cells labelled with cytoplasmatic GFP (expressed from the *serW* promoter) and stained with calcofluor white (CFW) to visualize cellulose and HiLyte to label amyloid structures. Spheroplasts and rod-shaped cells of the parental wild-type strain were used as a control. Arrowheads indicate spatially segregated bacteria. The scalebar represents 10 µm.

To determine whether the upregulated expression of regulatory genes involved in matrix production also enhanced extracellular matrix formation, we employed scanning electron microscopy (SEM). We observed a pronounced network of fibrillar structures associated with the L-form^0^ strain, in contrast to the relative smooth cell surface of spheroplasts and the parental wild-type (Fig. 3C). When the strain was reverted to walled growth, we still noticed an increased fibrillar network associated with the cell surface. Notably, the matrix was so extensive that it caused the cells to be spatially separated from each other (see Fig. 3D, arrowheads). To further characterize the extracellular matrix, we focused on two major components, cellulose and curli, and analyzed them using fluorescent dyes. While the revertant showed increased cellulose production compared to the wild type, the biggest increase was observed when comparing the L-form to spheroplasts (Fig. 3D). This was unexpected, as the parental strain (and the L-form strain) carries a *bcsQ* mutation that typically prevents cellulose production (Supplementary Fig. 8, (Serra, Richter et al. 2013). This prompted us to investigate a *rcsA*-overexpression strain, which also carries the *bcsQ* mutation (Fig. 4). Crucially, *rcsA* overexpression led to the upregulation of both cellulose and amyloids (see Fig. 4). These results suggest that elevated levels of RcsA can overcome the matrix production defects associated with the *bcsQ* mutation, enabling the production of cellulose and amyloids.

**Fig. 4.**
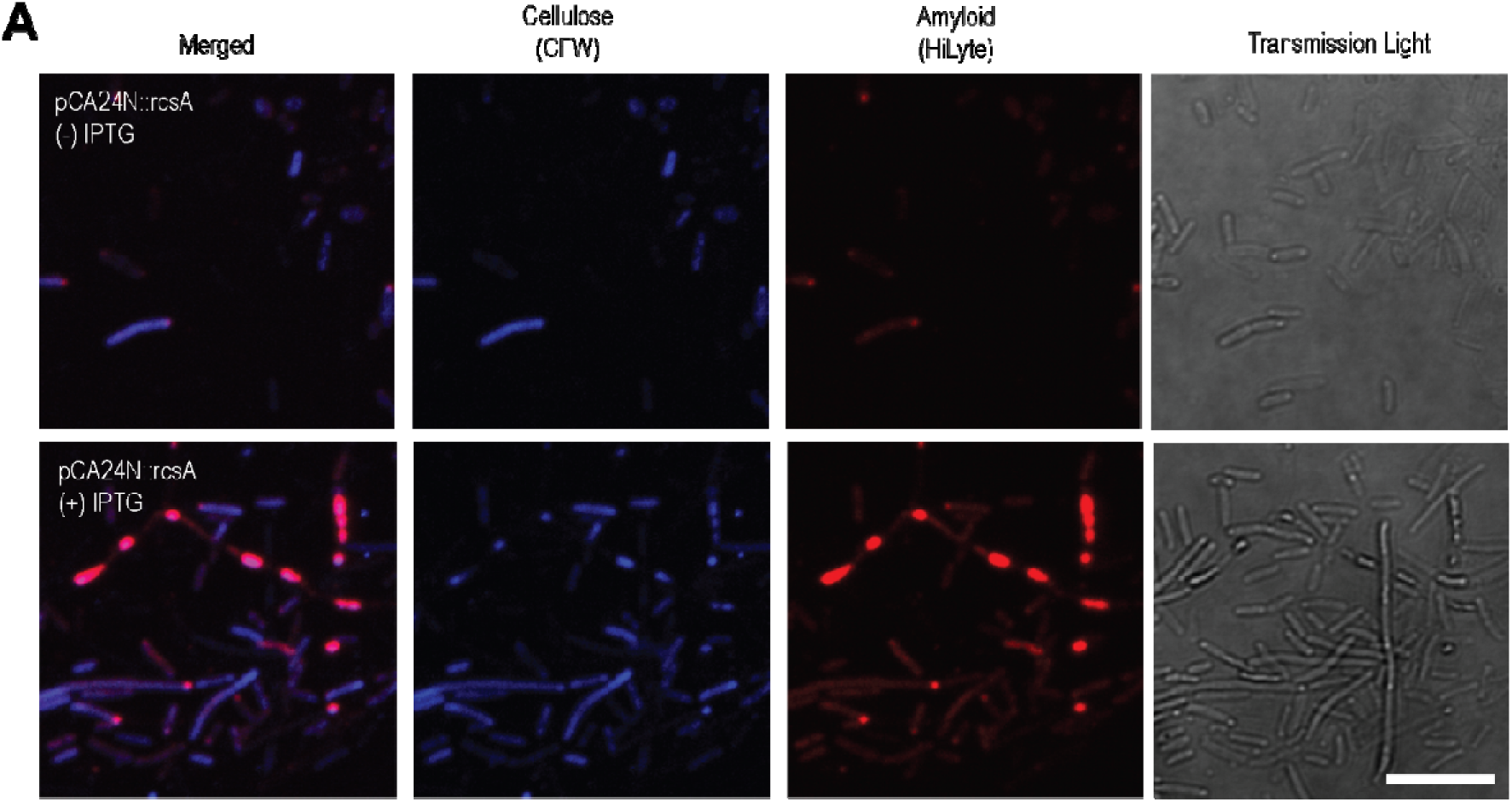
Overexpression of RcsA increases the production of cellulose and amyloids. (A) Fluorescence images of exponentially growing cells of the *rcsA*-inducible strain in LB medium in the absence (top panels) or presence (bottom panels) of IPTG. Cells were stained with CFW and HiLyte. The scalebar represents 10µm.

### Enhanced extracellular matrix production facilitates robust growth of cell wall-deficient L-forms

To study the effect of prolonged exposure to high-osmotic environments and wall-deficient growth, we evolved the L-form^0^ strain in LPB medium for 800 generations. The goal of this evolution experiment was to understand how continuous osmotic stress impacts bacterial adaptation and survival on the phenotypic and genotypic levels, especially in the wall-deficient state. The L-form^0^ strain maintained in a 37°C incubator and was transferred to a flask of fresh LPB medium supplemented with penicillin. To check for further mutations in *rcsA,* the promoter and coding sequence of this long-term evolution L-form (LTE), hereafter called “L-form^LTE^”, were amplified using PCR before sequencing and alignment to the sequence of the original L-form^0^ strain. Two additional SNPs were identified, one in the promoter region of *rcsA* and one in the gene itself (Table 2, Supplementary Table 4). One resulted in a silent mutation in the coding sequence, while the other SNP was in the promoter region of *rcsA* (Supplementary Fig. 9). The original deletion in the −10 region of the promoter remained, resulting in two changes in the promoter region of *rcsA* in the L-form^LTE^ strain compared to the wild type. In addition, we noticed that the prophage Rac was absent in the evolved strain (Supplementary Table 2). Interestingly, the loss of this prophage has been shown to promote enhanced biofilm formation (Liu, Li et al. 2015). Scanning electron microscopy (SEM) of the L-form^LTE^ strain revealed a denser and more organized network of the extracellular matrix, particularly evident in the revertant samples (Fig. 5A). Furthermore, fluorescent imaging demonstrated amyloid clustering in these revertants (Fig. 5B). Additional evidence of increased amyloid secretion was observed on LB plates supplemented with Congo Red (Fig. 5C), and the L-form^LTE^ strain exhibited stronger adhesion to polystyrene surfaces compared to the L-form^0^ strain (Fig. 5D, Supplementary Table 5). These findings are accompanied with a further decrease in ROS compared to the wild type and L-form^0^ (Supplementary Fig. 10, Supplementary Table 6). Importantly, these cells also grow faster and to a higher density compared to the original L-form^0^ strain (Fig. 5E), suggesting more robust growth under these conditions.

**Fig. 5.**
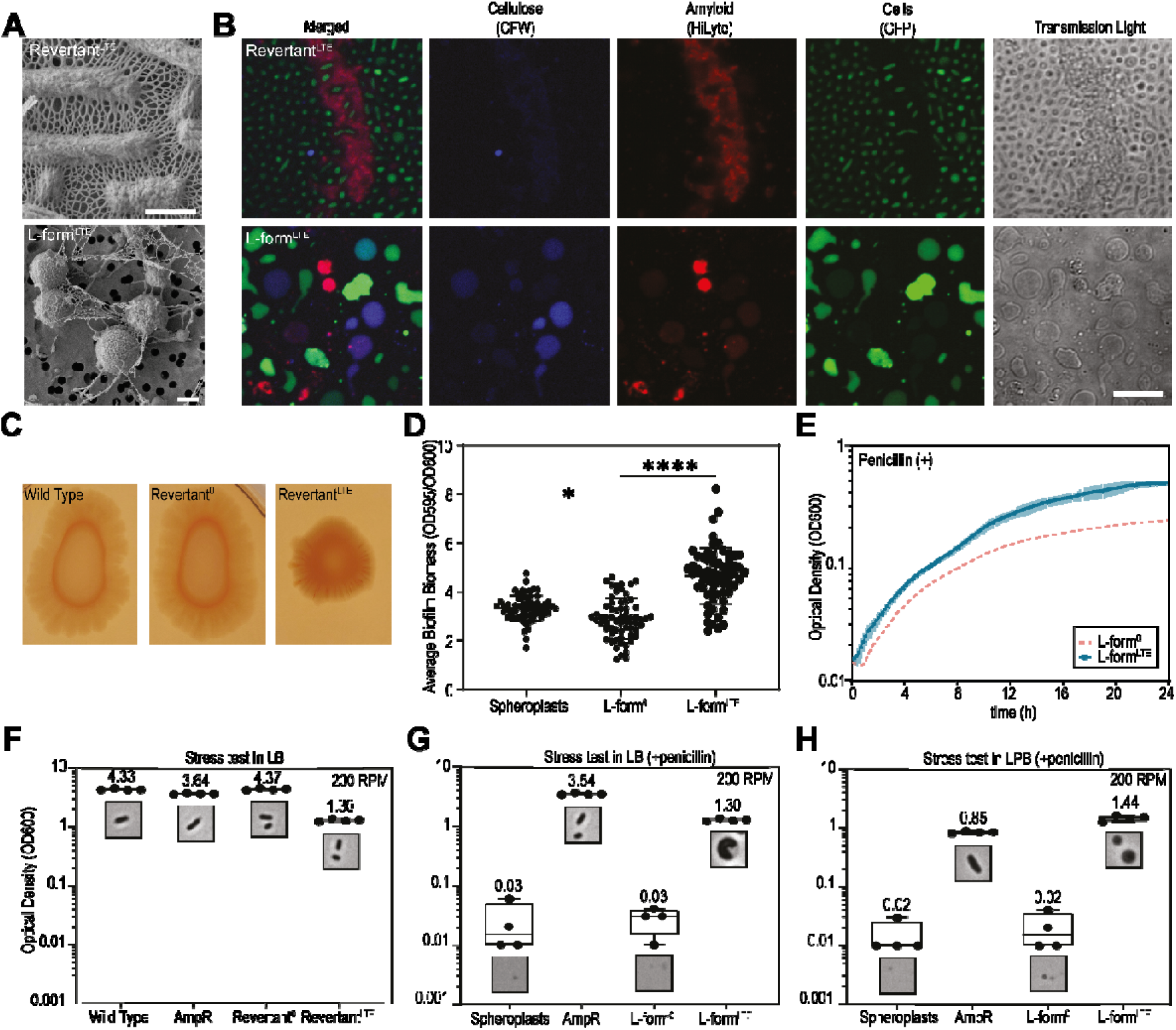
Long-term evolution enables robust growth of L-form cells along with increased extracellular matrix production. (A) SEM images of the long-term evolution strain L-form^LTE^ in the presence (top panels) or absence (bottom panels) of the cell wall. The scalebar is 1 µm. (B) Fluorescent images of cells of the L-form^LTE^ strain and its revertant stained with CFW and HiLyte to visualize cellulose and amyloid structures. The scalebar represents 10 µm. (C) Biofilm assay of the wild type, revertant^0^ and revertant^LTE^ strains. 20μl of culture was spotted on LB plates containing Congo Red dye to visualize amyloid formation (D) Quantification of the average biofilm biomass of wall-deficient cells of spheroplasts, L-form^0^ and L-form^LTE^ (Supplementary Table 5). Cells were grown in LPB medium supplemented with penicillin and stained with crystal violet. The P-values were determined using Kruskal-Wallis following Dunnett’s multiple comparisons test. (E) Growth curve of L-form^LTE^ grown in LPB medium with penicillin. The L-form^LTE^ strain is shown in blue. The red dotted line represents data from the L-form under comparable conditions, which is already shown in Fig. 1D, to compare the new data presented in the current figure. The data points and error bars represent the mean and standard deviation across replicates (n=3). (F-H) Stress test of walled cells (F) and wall-less cells (G-H) of the wild-type strain spheroplasts, ampicillin-resistant (AmpR), L-form^0^, and L-form^LTE^ cells grown overnight in baffled flasks at 200 RPM. Cells were grown in LB (F-G) and LPB medium (H). The AmpR cells remained walled in penicillin-rich conditions. Data is shown in box and whiskers (n=4).

**Table 2.**
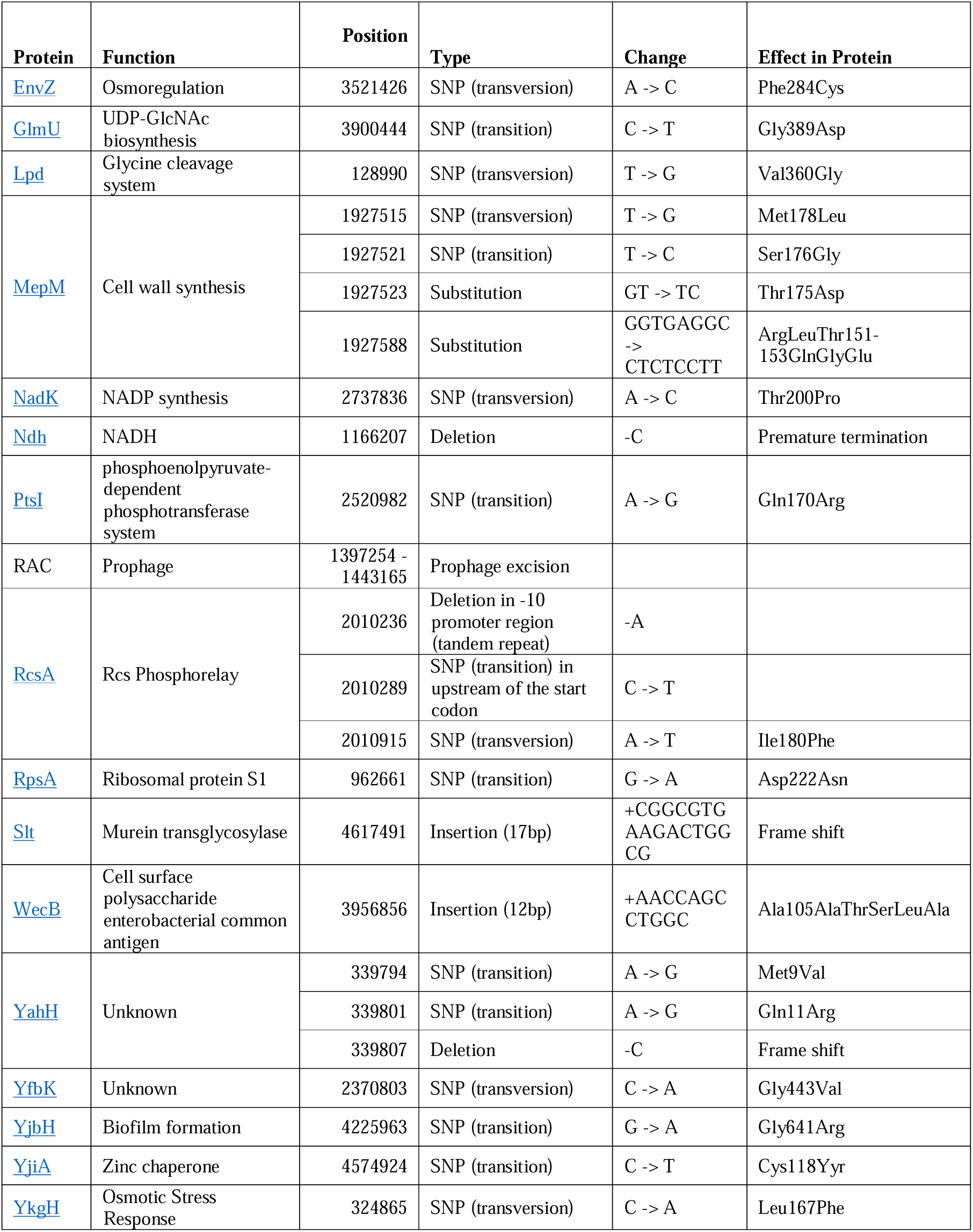
| List of mutations found in the L-form^LTE^ strain. Position reflects the genomic location of the L-form^LTE^ strain. Function is function of affected protein.

The enhanced synthesis of extracellular matrix and advanced growth of the L-form^LTE^ strain prompted us to further investigate its robustness. To this end, we compared the LTE strain to the original L-form^0^ strain and spheroplasts, by subjecting the cultures to growth in low-osmotic conditions and increased shear stress. Unlike the original L-form^0^ strain, the L-form^LTE^ strain could grow robustly in baffled flasks at 200 RPM (Fig. 5F-H, Supplementary Fig. 11), and even did so in media lacking osmotic protection (LB), see Fig. 5F. Interestingly, when comparing the growth of the L-form^LTE^ strain in LPB medium supplemented with penicillin to that of a beta-lactam resistant wild-type strain (growing with its cell wall), we found that the L-form^LTE^ strain outperformed the walled strain, reaching an optical density that is nearly 70% higher (Fig. 5H). Taken together, these results demonstrate that enhanced extracellular matrix production can effectively compensate for the absence of the cell wall in providing cellular robustness.

## Discussion

The peptidoglycan-based cell wall serves as a protective barrier against external stressors. While it remains an interesting target for antibiotics, some bacteria can proliferate without their canonical cell wall as L-forms. Bacterial life in the absence of a cell wall is delicate, with cells awaiting (re-)synthesis of nascent peptidoglycan. Our research on *E. coli* L-forms reveals that one potential mechanism cells use to cope with the loss of their cell wall is by increasing the formation of an extracellular matrix. Long-term adaptation to growth without a cell wall created a matrix barrier of extracellular components, resulting in increased robustness and efficient proliferation, even allowing cells to surpass growth of cells with a cell wall in particular conditions.

Previous research revealed that for efficient proliferation as an L-form, cells have to counteract the formation of reactive oxygen species (ROS), while making membranes that are sufficiently fluid (Mercier, Dominguez-Cuevas et al. 2012, Kawai, Mercier et al. 2015). In this study, we found that the newly generated L-form strain also acquired mutations that result in ROS reduction, by inactivating NADH dehydrogenase encoded by the *ndh* gene. Furthermore, expression of GnsB, involved in mediating membrane fluidity, was significantly upregulated (0.0 to 135 transcript per million) in L-forms (Supplementary Data 1). Overexpression of *gnsB* leads to increased levels of unsaturated fatty acids and suppresses cold-sensitive mutations (Sugai, Shimizu et al. 2001). This gene has not been linked to *E. coli* L-forms before and may offer new insights into L-form proliferation.

Our main finding reveals that increased extracellular matrix production facilitates growth of *E. coli* cells lacking their cell wall (Fig. 5). The increased production of extracellular matrix results in the formation of a dense layer of polysaccharides and proteins around the cells, resembling a biofilm structure. This matrix appears to act as a protective structure, compensating for the absence of the cell wall by shielding the cells against osmotic pressure and shear stress. Elevated levels of extracellular matrix have been linked to enhanced antibiotic resistance (Høiby, Bjarnsholt et al. 2010, Flemming, van Hullebusch et al. 2023, Pai, Patil et al. 2023), and are associated with chronic and recurrent infections (Ascenzioni, Cloeckaert et al. 2021). Notably, *E. coli* L-forms have been implicated in recurrent urinary tract infections (Mickiewicz, Kawai et al. 2019). A better understanding of the survival mechanisms used by L-forms, such as increased extracellular matrix production, will aid in developing new intervention strategies to combat infections involving L-forms and counteract antibiotic resistance.

The ability of the L-form strain to produce more matrix is coupled to the increased expression of *rcsA*. Although RcsA is recognized as an activator of colanic acid synthesis (Laubacher and Ades 2008, Meng, Young et al. 2021), our findings indicate that elevated levels of RcsA also enhance the synthesis of curli and cellulose as shown in Fig. 4. This is notable because the strain retains the *bcsQ* mutation, which disrupts the cellulose synthesis pathway (Serra, Richter et al. 2013). To understand how increased RcsA can lead to elevated expression of curli and cellulose, we looked at the modulators of biofilm development. A central regulator controlling expression of curli proteins and cellulose is CsgD (Vianney, Jubelin et al. 2005, Yan, Chen et al. 2023). The expression of *csgD* itself is controlled by the RNA polymerase sigma factor RpoS. Additionally, several small regulatory RNAs, including *dsrA*, *rprA, mcaS,* and *iraP*, fine-tune the expression of curli and cellulose through numerous feedback loops (Grabowicz and Silhavy 2017, Fröhlich and Gottesman 2018). Here, we propose a model in which RcsA increases synthesis of cellulose and curli via RprA*-*activated RpoS, which in turn upregulates CsgD (Supplementary Fig. 12). This leads to increased curli and cellulose synthesis, collectively contributing to survival of the cells.

Lastly, our work suggests that the enhanced extracellular matrix can compensate for lack of cell wall. Excitingly, the morphology of the L-form strain described here is remarkably similar to cells of the archaeon *Candidatus Altarchaeum hamiconexum* (formerly known as SM1 Euryarchaeon) (Perras, Wanner et al. 2014), which are also embedded in a thick matrix (Probst, Holman et al. 2013). This diderm species is found in subterranean sediments in swamps, and, just like wall-deficient bacteria, considered difficult to cultivate (Probst, Weinmaier et al. 2014). This archaeon has no peptidoglycan or pseudopeptidoglycan and lacks all associated biosynthesis proteins in its genome (Witwinowski, Sartori-Rupp et al. 2022). However, it is unknown what the composition of the matrix of *Candidatus Altarchaeum hamiconexum* is (Perras, Wanner et al. 2014), analogs of genes involved in the synthesis of cellulose, curli and colanic acid can be found in several archaea species. A big obstacle is the fact that archaea are much more difficult to culture and study. However, when blasted similarities of colanic acid initiating enzyme *wcaJ* and catalytic subunit of celllose synthesis *bcsA* are found in several uncultured archaeon DNA samples (data not shown). The similarities in matrix production between the *E. coli* L-form strain and the archaeon indicate a robust mechanism for overcoming the challenges of cell survival in conditions where the cell wall is absent.

## Methods

### Bacterial strains and culture conditions

Bacterial strains used in this study are shown in Supplementary Table 7. The Δ*rcsA*, Δ*rcsB*, Δ*rcsC*, Δ*rcsD,* and Δ*rcsF* strains used in this study were obtained from the KEIO collection (Baba, Ara et al. 2006). The reporter strains SerW-GFP and promoter-less U66-GFP strain were obtained from the *E. coli* promoter collection (Zaslaver, Bren et al. 2006). The RcsA overexpression strain was obtained from the ASKA (-) collection (Kitagawa, Ara et al. 2005). All primers used in this study are found in Supplementary Table 8.

#### Media

Cell-wall deficient cells were grown at 37°C in L-phase broth (LPB) consisting of a 1:1 mixture of YEME and TSBS containing 25 mM MgCl_2_, supplemented with 50 μg ml^−1^ kanamycin (Sigma) and 0.4 mg ml^−1^ benzylpenicillin (Sigma). Walled cells were grown in LPB or LB low salt at 37°C, supplemented with 50 μg ml^−1^ kanamycin, 25 μg ml^−1^ chloramphenicol (Sigma) and/or 0.5mM IPTG (Sigma) if necessary. For growth on solid medium, L-phase media agar (LPMA) plates containing, LPB, 5% horse serum, 0.75% Iberian agar (w/v), supplemented with 50 μg ml^−1^ kanamycin and 0.4 mg ml^−1^ penicillin if needed.

#### Formation of spheroplasts, L-form^0^ and L-form^LTE^

To obtain spheroplasts, an overnight culture of *E. coli* K-12 MG1655 was diluted to OD=0.01 in LPB medium, supplemented 50 μg ml^−1^ kanamycin and 0.4 mg ml^−1^ penicillin and grown overnight at 37°C. The generation of L-form^0^ and L-form^LTE^ are described in our recent manuscript (Crooijmans et al., *manuscript under review*).

#### Growth curves

Overnight cultures of cells grown in LPB and LB medium were measured with Ultrospec™ 2100 pro UV-Visible spectrophotometer. Then cells were diluted to OD_600_=0.05 in flat base 96-well cell culture plate (Sarstedt) and grown in a Tecan i-control, Infinite 200Pro plate reader at 37 while shaking for 2 s in 6 mm orbital amplitude, resulting in an RPM of 162. Every 15 mins, the average absorbance in each well was determined as the average of 5 measurements per well (once in the center and 4 times 1500 µm of center in in a rectangle shape). Data were normalized by removing the optical density of the used media. Points and error bars represent the mean and standard deviation across replicates.

#### Stress survival tests

The OD_600_ of overnight cultures of spheroplasts, L-form^0^ and L-form^LTE^ cells were measured with an Ultrospec™ 2100 pro UV-Visible spectrophotometer, after which cells were diluted to OD_600_=0.05 in LPB or LB medium, supplemented with penicillin (0.4 mg ml^−1^) if necessary, and grown in 250 ml baffled flasks (Corning) at 200RPM. After 24 hours, the OD_600_ was remeasured using the spectrophotometer. The experiment was performed in quadruplicate.

### Microscopy

#### Transmitted light microscopy

Cultures were imaged with a Zeiss Axio Lab A1 A-plan 40x/0.65 Ph2 microscope equipped with a Zeiss Axiocam 105 camera and Zeiss 2.5 Lite software. Cell sizes were measured using FIJI (Schindelin, Arganda-Carreras et al. 2012).

#### Fluorescence microscopy

All confocal microscopy pictures were taken with an inverted Zeiss LSM 900 confocal microscope with Airyscan 2 module with Zeiss Zen 3.1 software (blue edition, Zeiss). Cells were imaged underneath an LPMA agar pad on an 8-well μ-Slide (Ibidi). Fluorescent dyes were added to exponentially growing cells for one hour, unless stated otherwise. The C-Apochromat 63x/1.20 W Korr UV VIS IR objective was to image GFP, HiLyte^TM^ Fluor 647 (Anaspec Inc.), calcofluor white (CFW) (Sigma), and CellROX Orange (Invitrogen). For GFP, cells were excited at 488 nm and observed using an emission filter of 509 nm. Cells treated with 0.5µM HiLyte were excited at 640 nm an fluorescence emission was captured at 668nm. Cells treated with 0.1 mg ml^−1^ CFW were excited at 405 nm and fluorescence emission was captured at 432 nm. For the oxidation stress, CellROX was added to cells at a final concentration of 15 µM treated with or without 1 mM H_2_0_2_ (Sigma) in flasks for three hours at 37°C. These cells were excited at 561 nm and fluorescence emission captured at 570 nm. For the promoter activity of *rcsA*, cells of the exponentially growing reporter strain *rcsA*-GFP and the promoter-less strain U66-GFP were incubated for three hours in LPB supplemented with or without penicillin 0.4 mg ml^−1^. The C-Apochromat 40x/1.20 W Korr objective was used to image DNA stained with SYTO^TM^ 63 using 640 nm excitation and 681 nm emission.

#### Time-lapse microscopy

To visualize the growth of L-forms cells and the formation of spheroplast, cells were imaged using the Lionheart FX Automated Microscope (Agilent) with software Gen5 version 3.12. To this end, cells were diluted in a 96-well Greiner Bio-One SensoPlate (non-treated black plate with clear bottom) spun down to settle the mixture and overlaid with an LPMA patch containing antibiotics, if needed. Bright field images were taken every 10 min with the 40x objective at 37°C.

#### Scanning electron microscopy

Overnight cultures were fixed in a 1.5% glutaraldehyde solution in LPB medium for 15 minutes. This solution was fixated to a Millichrom, 0.4 μm Isopore polycarbonate membrane of 13mm disks. After dehydration in ethanol (70%, 80%, 90%, 100%) and critical point drying (Baltec CPD-030), sample was coated with 10 nm platinum/palladium and imaged with a JEOL JSM6700F scanning electron microscope.

### Flow Cytometry

CellROX Orange was measured using a Bio-Rad S3e Cell Sorter. In total, 100,000 cells were measured following excitation at 561 nm and 586/25 nm emission filter. Data was analyzed using FlowJo, while statistics were performed in Prism. The amount of fluorescence is indicated in relative arbitrary units.

### Genomic sequencing and SNP analysis of the L-form strain

Detailed description of the DNA isolation, Illumina sequencing, reads alignment and annotation, are described elsewhere (Crooijmans et al., *manuscript under review*). The Illumina sequencing is performed by BaseClear, Leiden, the Netherlands. The SNP variants were identified using Geneious with a minimum sequence coverage of 50 and a variant frequency of 100%. ColabFold v1.5.3: AlphaFold2 (MMseqs2) was used to predict the 3D protein structure of the new protein sequences. Figures were made using PyMOL (version 2.5.4).

### RNA sequencing

RNA was isolated using phenol/chloroform of a fresh batch of spheroplasts and mid-exponentially growing L-form cells (Toni, Garcia et al. 2018). As reference, RNA of *E. coli* MG1655 U66-GFP grown in LPB medium (without penicillin) was included. RNA sequencing was performed by BGI (Hongkong, China). Data were analyzed and visualized using BGI’s Dr.Tom (Venn diagram, DEG analysis) and GraphPad Prism 9.0.0 (bar plots) software. The statistical analysis was performed using a Kruskal-Wallis nonparametric ANOVA with Dunn’s multiple comparisons test.

### Quantification of matrix components

#### Congo Red assay

For the detection of amyloids, 20 μl of overnight cultures of the wild type strain and revertant^0^ and revertant^LTE^ were spotted on LB plates supplemented with kanamycin and 30 μg ml^−1^ Congo Red. Plates were grown for 48 hours before imaging.

#### Crystal violet assay

For the quantification of biofilm, cells were grown for 72 hours in LPB medium supplemented with penicillin at 37°C in a Greiner Bio-One 384-well, medium binding treated, black plate with clear bottom covered with Greiner Bio-One BREATHseal. After OD_600_ was measured, cells were removed and wells were gently washed four times before airdrying. Then, wells were filled with 0.1% crystal violet solution (in deionized water) was added to each well for 20 minutes to stain the adhering biofilm. After incubation, the excess dye was removed, and wells were washed 4 times with deionized water before drying. The dye associated with the biofilm was subsequently extracted by adding 50 μl 33% acetic acid to each well and incubation for 30 minutes. Finally, the extracted dye was quantified by measuring plates (OD_595_) using a Tecan Spark 10M. Biofilm biomass was normalized with the OD_600_ values. Outliers detected using ROUT (Motulsky and Brown 2006) were excluded from the dataset.

### Statistics

GraphPad Prism 9.0.0 was used to perform statistical tests and create graphs. Ns: not significant, * = P ≤ 0.05, ** = P ≤ 0.01, *** = P ≤ 0.001, and **** = P ≤ 0.0001.

### Data availability

RNA sequencing are deposited in the Sequence Read Archive (SRA) database of NCBI. The accession number of the RNA seq data is SAMN43528363 (wild type), SAMN43528364 (L-form^0^), and SAMN43528362 (spheroplast) under the BioProject PRJNA1156547. All microscopy data is stored at Leiden University in Omero (Allan, Burel et al. 2012).

## Acknowledgments

This work was funded by the VICI grant (no. VI.C.192.002) from the Dutch Research Council to Dennis Claessen. Lennart Schada von Borzyskowski was so kind to provide us with the KEIO and ASKA collection strains. We thank Gerda Lamers for providing microscopy support and Ben Nannings for the background work on L-form standardization. Lastly, we thank Alexandra Possling for letting us borrow the W3110 and AR3110 strains.

## Author contributions

M.C. and D.C. designed the experiments. M.C. and J.W., collected data. M.C., H.D.W. and D.C. aided in data analysis. D.C and H.D.W. supervised the research. D.C. provided funding. All authors approved the submitted version.

## Additional information

**Supplementary Information:** contains Supplementary Tables and Supplementary Figures.

**Supplementary Data:** contains supplementary RNA sequencing data.

**Supplementary Videos:** contains time-lapse microscopy videos.

Supplementary Video 1: L-form^0^ grown under LPMA pad with penicillin.

Supplementary Video 2: Wild type (U66-GFP) to spheroplast grown under LPMA pad with penicillin.

Supplementary Video 3: Wild type (U66-GFP) grown under LPMA pad.

Supplementary Video 4: L-form^0^ to revertant grown under LPMA pad.

Supplementary Video 5: Spheroplasts to rod-shaped grown under LPMA pad.

