## Supplementary Data 1 for "Enhanced extracellular matrix production provides protection to cell wall-deficient *Escherichia coli*"

Supplementary Data 1. List of genes uniquely expressed in L-forms, spheroplasts, and wild type. Venn diagram of RNA sequencing data.

| Gene ID | Gene Symbol | | L-form Average TPM | Wild-Type-LPB Average TPM | Spheroplasts Average TPM | GO_C Desc | GO_F Desc | GO_P Desc |
|---|---|---|---|---|---|---|---|---|
| **L-form only expressed genes** | | | | | | | | |
| 946054 | gnsB' | mRNA | 134.60 | 0.00 | 0.00 | GO:0005829 | | GO:0006636 |
| 38094963 | ynfQ' | mRNA | 111.07 | 0.00 | 0.00 | | | |
| 948344 | 'yigG' | mRNA | 18.93 | 0.00 | 0.00 | GO:0005886+++GO:0005887+++GO:0016020+++GO:0016021 | | |
| 5061515 | 'ymgI' | mRNA | 16.83 | 0.00 | 0.00 | | | |
| 946958 | 'ydcD' | mRNA | 8.45 | 0.00 | 0.00 | | | |
| 1450240 | 'ylcG' | mRNA | 8.31 | 0.00 | 0.00 | | | |
| 948340 | 'yigF' | mRNA | 7.06 | 0.00 | 0.00 | GO:0005886 | | |
| 945729 | 'iraM' | mRNA | 2.41 | 0.00 | 0.00 | GO:0005737 | GO:0005515+++( | GO:0009267+++GO:0010350+++GO:0042177+++GO:0045893+++GO:0071468 |
| 946024 | 'yncH' | mRNA | 2.21 | 0.00 | 0.00 | | | |
| 1450242 | 'rzoD' | mRNA | 2.09 | 0.00 | 0.00 | GO:0009279+++GO:0016020 | | GO:0019076+++GO:0019835 |
| 947336 | 'ygeI' | mRNA | 1.98 | 0.00 | 0.00 | | | |
| 946061 | 'safA' | mRNA | 1.39 | 0.00 | 0.00 | GO:0005886+++GC | GO:0005515 | GO:0010447 |
| 945992 | 'ydfO' | mRNA | 1.07 | 0.00 | 0.00 | | | |
| 5061516 | 'ymgJ' | mRNA | 1.06 | 0.00 | 0.00 | | | |
| 5061498 | 'ykfM' | mRNA | 0.80 | 0.00 | 0.00 | GO:0016020+++GO:0016021 | | GO:0006974+++GO:0046677 |
| 947312 | 'yddK' | mRNA | 0.64 | 0.00 | 0.00 | | | |
| 947557 | 'yhaC' | mRNA | 0.15 | 0.00 | 0.00 | | | GO:0006974 |
| 947329 | 'yqeK' | mRNA | 0.05 | 0.00 | 0.00 | | | |
| 947640 | 'ypjC' | mRNA | 0.04 | 0.00 | 0.00 | | | |
| **Wild Type only expressed genes** | | | | | | | | |
| 5061511 | 'ypaB' | mRNA | 0.00 | 1.22 | 0.00 | | | |
| 5061519 | 'ygdT' | mRNA | 0.00 | 1.06 | 0.00 | | | |
| **Spheroplast only expressed genes** | | | | | | | | |
| 945911 | 'ydaF' | mRNA | 0.00 | 0.00 | 5.49 | | | |
| 945907 | 'ydaG' | mRNA | 0.00 | 0.00 | 5.00 | | | |
| 947417 | 'ynaK' | mRNA | 0.00 | 0.00 | 2.11 | GO:0005694 | | GO:0007059+++GO:0045881 |
| 945629 | 'yceO' | mRNA | 0.00 | 0.00 | 1.67 | GO:0005887 | | GO:0010447+++GO:0042710 |
| 948959 | 'ydaE' | mRNA | 0.00 | 0.00 | 1.51 | | GO:0008270 | |
| 947399 | 'xisR' | mRNA | 0.00 | 0.00 | 0.98 | | GO:1990837 | GO:0032359 |
| 38094954 | 'ymjE' | mRNA | 0.00 | 0.00 | 0.72 | | | |
| 945921 | 'kilR' | mRNA | 0.00 | 0.00 | 0.72 | | | GO:0046677+++GO:0051301+++GO:0051782 |
| 5061514 | 'yojO' | mRNA | 0.00 | 0.00 | 0.63 | | | |
| 946110 | 'dicB' | mRNA | 0.00 | 0.00 | 0.60 | | GO:0005515 | GO:0051301+++GO:0051302+++GO:0051782 |
| 945920 | 'racC' | mRNA | 0.00 | 0.00 | 0.21 | | | |
| **Spheroplast and Wild Type expressed genes** | | | | | | | | |
| 945944 | insH-5' | mRNA | 0.00 | 9.46 | 39.80 | GO:0005829 | GO:0003677+++( | GO:0006310+++GO:0006313+++GO:0032196+++GO:0045893 |
| 4056027 | 'ydbJ' | mRNA | 0.00 | 81.39 | 28.55 | | | |
| 945927 | 'ydaW' | mRNA | 0.00 | 5.32 | 10.36 | | GO:0005515 | |
| 1450262 | 'rzoR' | mRNA | 0.00 | 3.58 | 7.63 | GO:0009279+++GO:0016020 | | GO:0019076+++GO:0019835 |
| 946062 | 'tfaR' | mRNA | 0.00 | 1.77 | 5.41 | | | |
| 945924 | 'ydaT' | mRNA | 0.00 | 4.00 | 5.35 | | | GO:0043093 |
| 945925 | 'ydaU' | mRNA | 0.00 | 2.93 | 4.63 | | | |
| 945923 | 'ydaS' | mRNA | 0.00 | 4.24 | 4.46 | | GO:0003677+++( | GO:0008219 |
| 945207 | 'stfR' | mRNA | 0.00 | 1.72 | 3.84 | | GO:0005198 | |
| 945959 | 'paaD' | mRNA | 0.00 | 1.37 | 3.73 | | | GO:0010124 |
| 945926 | 'ydaV' | mRNA | 0.00 | 3.01 | 3.51 | | GO:0005524 | GO:0006260+++GO:0006271 |
| 945939 | 'tynA' | mRNA | 0.00 | 1.17 | 2.53 | GO:0042597 | GO:0005507+++( | GO:0006559+++GO:0009308+++GO:0019607+++GO:0055114 |
| 946313 | 'ompN' | mRNA | 0.00 | 0.32 | 2.47 | GO:0009279+++GC | GO:0015288 | GO:0006811+++GO:0034219+++GO:0055085 |
| 945917 | 'recT' | mRNA | 0.00 | 1.87 | 2.36 | GO:0032993 | GO:0003677+++( | GO:0006259+++GO:0006310+++GO:0032508+++GO:0043150 |
| 945946 | 'ynbE' | mRNA | 0.00 | 1.65 | 2.32 | | | |
| 946208 | 'ydaY' | mRNA | 0.00 | 0.12 | 2.27 | | GO:0005515 | |
| 947595 | 'paaB' | mRNA | 0.00 | 0.93 | 2.02 | | GO:0005515 | GO:0010124 |
| 945918 | 'recE' | mRNA | 0.00 | 1.24 | 1.60 | | GO:0004518+++( | GO:0006259+++GO:0090305 |
| 946078 | 'ynaE' | mRNA | 0.00 | 0.38 | 1.59 | | | GO:0009409 |
| 945035 | 'yahH' | mRNA | 0.00 | 1.06 | 1.33 | | | |
| 945956 | 'paaC' | mRNA | 0.00 | 0.49 | 1.23 | GO:0005829 | GO:0005515 | GO:0010124 |
| 945833 | 'paaA' | mRNA | 0.00 | 0.23 | 0.97 | GO:0005829 | GO:0005515+++( | GO:0010124+++GO:0055114 |
| 38094967 | 'yecU' | mRNA | 0.00 | 17.74 | 0.91 | | | |
| 945954 | 'paaZ' | mRNA | 0.00 | 0.30 | 0.89 | | GO:0003824+++( | GO:0008152+++GO:0010124+++GO:0055114 |
| 947543 | 'ynaA' | mRNA | 0.00 | 0.34 | 0.88 | | | |
| 945913 | 'sieB' | mRNA | 0.00 | 0.68 | 0.66 | GO:0005886+++GO:0016020+++GO:0016021 | | |
| 945914 | 'ralR' | mRNA | 0.00 | 0.54 | 0.54 | | GO:0004518+++( | GO:0009307+++GO:0046677+++GO:0090305 |
| 5061499 | 'ylcI' | mRNA | 0.00 | 0.28 | 0.27 | | | |
| 947504 | 'rcbA' | mRNA | 0.00 | 0.45 | 0.23 | | GO:0005515 | GO:0006259+++GO:0046677 |
| **L-form and Wild Type expressed genes** | | | | | | | | |
| 946951 | ymcE' | mRNA | 47.06 | 0.17 | 0.00 | GO:0005886+++GO:0005887+++GO:0016020+++GO:0016021 | | |
| 1450238 | 'ykgO' | mRNA | 1.59 | 1.46 | 0.00 | GO:0005840 | GO:0003735 | GO:0006412 |
| 945737 | 'yddL' | mRNA | 0.65 | 0.09 | 0.00 | GO:0016020 | GO:0015288 | GO:0055085 |
| 2847744 | 'hokC' | mRNA | 0.55 | 0.26 | 0.00 | GO:0005886+++GO:0005887+++GO:0016020+++GO:0016021+++GO:0042597 | | |
| 1450308 | 'ghoT' | mRNA | 0.46 | 2.26 | 0.00 | GO:0005886+++GO:0016020+++GO: | | GO:0008219 |
| 946986 | 'ygeG' | mRNA | 0.26 | 0.03 | 0.00 | | | |
| 948953 | 'ybfK' | mRNA | 0.12 | 0.13 | 0.00 | | | |
| 945953 | 'smrA' | mRNA | 0.03 | 0.15 | 0.00 | | GO:0003677+++( | GO:0006259+++GO:0090305 |
| **L-form and Spheroplast expressed genes** | | | | | | | | |
| 38094933 | 'yabQ' | mRNA | 75.67 | 0.00 | 0.88 | | | |
| 946097 | 'ynfN' | mRNA | 13.77 | 0.00 | 5.15 | | | |
| 7751632 | 'ykgS' | mRNA | 6.95 | 0.00 | 13.11 | | | |
| 946176 | 'ydfB' | mRNA | 6.93 | 0.00 | 3.93 | GO:0005737 | | |
| 948704 | 'yjfI' | mRNA | 4.19 | 0.00 | 0.49 | | | |
| 1450267 | 'ynfO' | mRNA | 3.18 | 0.00 | 0.61 | | | |
| 945178 | 'ybcV' | mRNA | 2.85 | 0.00 | 0.05 | | | |
| 945183 | 'tfaX' | mRNA | 1.44 | 0.00 | 0.36 | | | |
| 947566 | 'ybfB' | mRNA | 1.18 | 0.00 | 0.07 | GO:0005886+++GO:0016020+++GO:0016021 | | |
| 945415 | 'citD' | mRNA | 0.98 | 0.00 | 0.62 | GO:0005737 | GO:0051192 | |
| 946621 | 'yehC' | mRNA | 0.57 | 0.00 | 0.02 | GO:0030288+++GO:0042597 | | GO:0006457+++GO:0043711+++GO:0061077+++GO:0071555 |
| 5061530 | 'ytcA' | mRNA | 0.52 | 0.00 | 0.10 | GO:0005886+++GO:0016020+++GO:0016021 | | |
| 947700 | 'yhdU' | mRNA | 0.40 | 0.00 | 1.19 | GO:0005886+++GO:0005887+++GO:0016020+++GO:0016021 | | |
| 944929 | 'yafW' | mRNA | 0.38 | 0.00 | 0.40 | | | GO:0051495 |
| 948839 | 'ykfF' | mRNA | 0.30 | 0.00 | 0.47 | | | |
| 948576 | 'yjcF' | mRNA | 0.23 | 0.00 | 0.01 | | | |
| 945901 | 'ynaJ' | mRNA | 0.13 | 0.00 | 0.12 | GO:0005886+++GO:0016020+++GO:0016021 | | |
