## Supplementary Data 2 for "Enhanced extracellular matrix production provides protection to cell wall-deficient *Escherichia coli*"

Supplementary Data 2. The 150 most upregulated DEGs of the L-form compared to wild type.

| | Gene ID | Gene Symbol | Type | log2 (L-form / Wild-Type-LPB) | Qvalue (L-form / Wild-Type-LPB) | Lform_1 TPM | Lform_2 TPM | Lform_3 TPM | WT_1 TPM | WT_2 TPM | WT_3 TPM | GO_C Desc | GO_F Desc | GO_P Desc |
|---|---|---|---|---|---|---|---|---|---|---|---|---|---|---|
| 1 | 946558 | 'wza' | mRNA | 10.29067 | 2.73E-137 | 1318.79 | 1079.58 | 2256.74 | 0.28 | 0.35 | 0.66 | GO:00092 | GO:00151 | GO:0000271///polysaccharide biosynthetic process+++GO:0006811///ion transport+++GO:0008643///carbohydrate transport+++GO:0015774///polysaccharide transport+++GO:0046377///colanic acid metabolic process+++GO:0055085///transmembrane transport |
| 2 | 946562 | 'gmd' | mRNA | 10.26405 | 5.53E-174 | 3517.18 | 3328.89 | 4391.92 | 0.6 | 0.96 | 1.64 | GO:00057 | GO:00084 | GO:0009242///colanic acid biosynthetic process+++GO:0019673///GDP-mannose metabolic process+++GO:0042351///'de novo' GDP-L-fucose biosynthetic process |
| 3 | 946564 | 'wzb' | mRNA | 10.07434 | 8.55E-87 | 1380.84 | 1483.42 | 1429 | 0.36 | 0.37 | 0.64 | | GO:00047 | GO:0000271///polysaccharide biosynthetic process+++GO:0006470///protein dephosphorylation+++GO:0009242///colanic acid biosynthetic process+++GO:0035335///peptidyl-tyrosine dephosphorylation |
| 4 | 946578 | 'wcaF' | mRNA | 10.06062 | 1.73E-57 | 562.29 | 464.68 | 656.29 | 0.09 | 0.18 | 0.28 | | GO:00164 | GO:0009103///lipopolysaccharide biosynthetic process+++GO:0044010///single-species biofilm formation+++GO:0045228///slime layer polysaccharide biosynthetic process+++GO:0046377///colanic acid metabolic process |
| 5 | 946563 | 'fcl' | mRNA | 9.751977 | 1.62E-113 | 1497.68 | 1498.65 | 2503.89 | 0.88 | 0.26 | 1.07 | GO:00057 | GO:00038 | GO:0008152///metabolic process+++GO:0009226///nucleotide-sugar biosynthetic process+++GO:0009242///colanic acid biosynthetic process+++GO:0042351///'de novo' GDP-L-fucose biosynthetic process+++GO:0055114///oxidation-reduction process |
| 6 | 946543 | 'wcaE' | mRNA | 9.246853 | 3.79E-120 | 873.82 | 738.16 | 1576.21 | 0.46 | 0.59 | 0.74 | | GO:00167 | GO:0009103///lipopolysaccharide biosynthetic process+++GO:0045228///slime layer polysaccharide biosynthetic process+++GO:0046377///colanic acid metabolic process |
| 7 | 946567 | 'wzc' | mRNA | 9.039269 | 2.90E-285 | 814.33 | 799.8 | 800.75 | 0.45 | 0.63 | 0.57 | GO:00058 | GO:00001 | GO:0000271///polysaccharide biosynthetic process+++GO:0009103///lipopolysaccharide biosynthetic process+++GO:0009242///colanic acid biosynthetic process+++GO:0016310///phosphorylation+++GO:0038083///peptidyl-tyrosine autophosphorylation+++GO:0045226///extracellular polysaccharide biosynthetic process |
| 8 | 946559 | 'gmm' | mRNA | 8.990562 | 5.17E-80 | 1221.2 | 1479.62 | 922.64 | 1.07 | 0.65 | 0.8 | | GO:00002 | GO:0009103///lipopolysaccharide biosynthetic process |
| 9 | 948533 | 'yjbF' | mRNA | 8.945249 | 7.97E-91 | 460.02 | 385.53 | 618.39 | 0.43 | 0.22 | 0.38 | GO:0005886///plasm | | GO:0045226///extracellular polysaccharide biosynthetic process |
| 10 | 946550 | 'wcaD' | mRNA | 8.841565 | 1.94E-174 | 632.96 | 700.87 | 625.3 | 0.61 | 0.42 | 0.51 | GO:0005886///plasm | | GO:0009103///lipopolysaccharide biosynthetic process+++GO:0045228///slime layer polysaccharide biosynthetic process+++GO:0046377///colanic acid metabolic process |
| 11 | 948152 | 'yiaB' | mRNA | 8.401476 | 1.25E-11 | 78.86 | 51.54 | 344.26 | 0 | 0 | 0.41 | GO:0005886///plasm | | GO:0006974///cellular response to DNA damage stimulus |
| 12 | 946579 | 'wcaC' | mRNA | 8.35596 | 9.71E-223 | 708.25 | 712.91 | 701.42 | 0.77 | 0.78 | 0.78 | | GO:00167 | GO:0009103///lipopolysaccharide biosynthetic process+++GO:0045228///slime layer polysaccharide biosynthetic process+++GO:0046377///colanic acid metabolic process |
| 13 | 946573 | 'wcaB' | mRNA | 8.273241 | 3.19E-69 | 465.76 | 389.11 | 753.54 | 0.31 | 0.32 | 1.11 | GO:00057 | GO:00090 | GO:0006535///cysteine biosynthetic process from serine+++GO:0009103///lipopolysaccharide biosynthetic process+++GO:0045228///slime layer polysaccharide biosynthetic process+++GO:0046377///colanic acid metabolic process |
| 14 | 946467 | 'rcsA' | mRNA | 8.231135 | 1.67E-89 | 926.37 | 757.69 | 1878.59 | 1.1 | 0.97 | 1.87 | | GO:00036 | GO:0006351///transcription, DNA-templated+++GO:0006355///regulation of transcription, DNA-templated+++GO:0050896///response to stimulus |
| 15 | 948526 | 'yibG' | mRNA | 8.12245 | 1.92E-46 | 248.75 | 257.08 | 146.26 | 0.12 | 0.36 | 0.37 | | | GO:0045226///extracellular polysaccharide biosynthetic process |
| 16 | 946588 | 'wcaI' | mRNA | 8.050221 | 1.79E-267 | 844.17 | 841.67 | 903.61 | 1.11 | 1.06 | 1.31 | | GO:00167 | GO:0009103///lipopolysaccharide biosynthetic process+++GO:0045228///slime layer polysaccharide biosynthetic process+++GO:0046377///colanic acid metabolic process |
| 17 | 946580 | 'cpsB' | mRNA | 8.035029 | 4.25E-243 | 405.41 | 406.2 | 503.57 | 0.66 | 0.56 | 0.56 | | GO:00001 | GO:0000271///polysaccharide biosynthetic process+++GO:0005976///polysaccharide metabolic process+++GO:0006972///hyperosmotic response+++GO:0009058///biosynthetic process+++GO:0009103///lipopolysaccharide biosynthetic process+++GO:0009242///colanic acid biosynthetic process+++GO:0009298///GDP-mannose biosynthetic process |
| 18 | 946570 | 'wcaA' | mRNA | 7.904639 | 1.00E-97 | 951.92 | 672.64 | 1679.97 | 1.34 | 1.27 | 2.06 | | GO:00167 | GO:0009103///lipopolysaccharide biosynthetic process+++GO:0045228///slime layer polysaccharide biosynthetic process+++GO:0046377///colanic acid metabolic process |
| 19 | 946581 | 'wzxC' | mRNA | 7.85114 | 1.81E-189 | 362.23 | 363.28 | 433.25 | 0.41 | 0.68 | 0.7 | GO:0005886///plasm | | GO:0009103///lipopolysaccharide biosynthetic process+++GO:0009242///colanic acid biosynthetic process |
| 20 | 946569 | 'wcaK' | mRNA | 7.766667 | 1.59E-223 | 690.05 | 571.99 | 633.49 | 1.05 | 1.04 | 1.02 | | GO:00167 | GO:0009103///lipopolysaccharide biosynthetic process+++GO:0045228///slime layer polysaccharide biosynthetic process |
| 21 | 948534 | 'yjbE' | mRNA | 7.690035 | 3.17E-144 | 6284.14 | 7126.26 | 14175.92 | 8.5 | 15 | 14.73 | | | GO:0042710///biofilm formation+++GO:0045226///extracellular polysaccharide biosynthetic process |
| 22 | 945945 | 'yddA' | mRNA | 7.596849 | 7.07E-12 | 9.35 | 5.74 | 9.66 | 0 | 0.02 | 0.02 | GO:00058 | GO:00001 | GO:0055085///transmembrane transport |
| 23 | 946583 | 'wcaJ' | mRNA | 7.514338 | 3.75E-204 | 398.58 | 394.28 | 588.55 | 0.88 | 0.83 | 0.95 | GO:00058 | GO:00167 | GO:0000271///polysaccharide biosynthetic process+++GO:0009103///lipopolysaccharide biosynthetic process+++GO:0009242///colanic acid biosynthetic process |
| 24 | 946561 | 'wcaM' | mRNA | 7.486639 | 1.34E-131 | 422.48 | 346.48 | 630.68 | 0.66 | 1 | 1.07 | | | GO:0009103///lipopolysaccharide biosynthetic process+++GO:0045228///slime layer polysaccharide biosynthetic process |
| 25 | 948344 | 'yigG' | mRNA | 7.40463 | 0.023833 | 1.65 | 0.16 | 54.97 | 0 | 0 | 0 | GO:0005886///plasma membrane+++GO:0005887///integral component of plasma membrane+++GO:0016020///membrane+++GO:0016021///integral component of membrane | | |
| 26 | 946574 | 'cpsG' | mRNA | 7.362218 | 6.80E-156 | 842.84 | 754.36 | 1136.3 | 1.45 | 1.9 | 2.48 | | GO:00002 | GO:0005975///carbohydrate metabolic process+++GO:0009103///lipopolysaccharide biosynthetic process+++GO:0009298///GDP-mannose biosynthetic process+++GO:0071704///organic substance metabolic process |
| 27 | 948078 | 'yiaA' | mRNA | 7.266985 | 5.83E-15 | 52.82 | 33.92 | 238.49 | 0.25 | 0.25 | 0.13 | GO:0005886///plasm | | GO:0006974///cellular response to DNA damage stimulus |
| 28 | 948515 | 'yjaA' | mRNA | 7.145886 | 4.94E-10 | 47.4 | 47.62 | 31.95 | 0.15 | 0 | 0.16 | | | GO:0044011///single-species biofilm formation on inanimate substrate+++GO:0070301///cellular response to hydrogen peroxide+++GO:0071468///cellular response to acidic pH |
| 29 | 947451 | 'ydfK' | mRNA | 6.897004 | 1.25E-06 | 48.83 | 24.7 | 419.91 | 0.22 | 0.92 | 0 | | | GO:0009409///response to cold |
| 30 | 946822 | 'yfdI' | mRNA | 6.895823 | 1.00E-07 | 3.55 | 2.8 | 15.03 | 0 | 0.06 | 0 | GO:0005886///plasma membrane+++GO:0016020///membrane+++GO:0016021///integral component of membrane | | |
| 31 | 946565 | 'wcaL' | mRNA | 6.752932 | 6.50E-53 | 520.97 | 530.54 | 236.89 | 1.27 | 1.42 | 1.69 | | GO:00167 | GO:0009103///lipopolysaccharide biosynthetic process+++GO:0045228///slime layer polysaccharide biosynthetic process+++GO:0046377///colanic acid metabolic process |
| 32 | 945115 | 'ybbC' | mRNA | 6.740353 | 1.79E-04 | 3.5 | 9 | 69.57 | 0 | 0.17 | 0 | | | |
| 33 | 946571 | 'ugd' | mRNA | 6.383371 | 2.89E-70 | 1721.69 | 1428.34 | 3562.03 | 6.91 | 9.78 | 10.73 | | GO:00039 | GO:0000271///polysaccharide biosynthetic process+++GO:0006065///UDP-glucuronate biosynthetic process+++GO:0009103///lipopolysaccharide biosynthetic process+++GO:0009242///colanic acid biosynthetic process+++GO:0055114///oxidation-reduction process |
| 34 | 947165 | 'proX' | mRNA | 6.344946 | 1.17E-175 | 1352.13 | 1082.87 | 1897.24 | 6.82 | 6.81 | 4.99 | GO:00302 | GO:00055 | GO:0006865///amino acid transport+++GO:0006972///hyperosmotic response+++GO:0031460///glycine betaine transport+++GO:0055085///transmembrane transport |
| 35 | 946101 | 'rzpQ' | mRNA | 6.311566 | 1.49E-05 | 5.92 | 3.49 | 21.02 | 0 | 0 | 0.15 | | | |
| 36 | 946951 | 'ymcE' | mRNA | 6.174294 | 3.48E-06 | 34 | 29.04 | 78.14 | 0 | 0 | 0.52 | GO:0005886///plasma membrane+++GO:0005887///integral component of plasma membrane+++GO:0016020///membrane+++GO:0016021///integral component of membrane | | |
| 37 | 947148 | 'proV' | mRNA | 6.01647 | 3.12E-153 | 1752.8 | 1663.84 | 1476.51 | 9.04 | 10.3 | 7.92 | GO:00058 | GO:00001 | GO:0006865///amino acid transport+++GO:0006972///hyperosmotic response+++GO:0015837///amine transport+++GO:0031460///glycine betaine transport |
| 38 | 948340 | 'yigF' | mRNA | 5.956764 | 0.001605 | 1.49 | 0.94 | 18.74 | 0 | 0 | 0 | GO:0005886///plasma membrane | | |
| 39 | 946181 | 'ydiL' | mRNA | 5.873321 | 2.55E-05 | 14.55 | 9.15 | 8.63 | 0.17 | 0 | 0 | | GO:0003677///DNA binding | |
| 40 | 946239 | 'ynjI' | mRNA | 5.79566 | 3.79E-21 | 19.44 | 16.31 | 48.39 | 0.27 | 0.08 | 0.16 | GO:0005886///plasma membrane+++GO:0016020///membrane+++GO:0016021///integral component of membrane | | |
| 41 | 945679 | 'ycfT' | mRNA | 5.751633 | 3.05E-107 | 110.48 | 97.76 | 133.86 | 0.7 | 0.71 | 0.82 | GO:00058 | GO:0016747///transferase activity, transferring acyl groups other than amino-acyl groups | |
| 42 | 945659 | 'ycjM' | mRNA | 5.737795 | 8.65E-15 | 8.21 | 5.08 | 26.46 | 0.11 | 0.07 | 0.07 | | GO:00038 | GO:0005975///carbohydrate metabolic process |
| 43 | 948704 | 'yjfI' | mRNA | 5.569967 | 2.11E-04 | 3.3 | 2.56 | 6.71 | 0 | 0 | 0 | | | |
| 44 | 946274 | 'ydjE' | mRNA | 5.389041 | 1.47E-21 | 10.75 | 8.43 | 11.29 | 0.08 | 0.11 | 0.06 | GO:00058 | GO:00228 | GO:0055085///transmembrane transport |
| 45 | 945972 | 'ynbB' | mRNA | 5.374976 | 2.42E-08 | 5.92 | 5.11 | 5.04 | 0.05 | 0.09 | 0 | GO:00058 | GO:00167 | GO:0009273///peptidoglycan-based cell wall biogenesis |
| 46 | 945511 | 'entC' | mRNA | 5.277314 | 9.91E-53 | 543.83 | 591.65 | 343.71 | 4.09 | 5.2 | 4.61 | | GO:00002 | GO:0009058///biosynthetic process+++GO:0009239///enterobactin biosynthetic process+++GO:0009697///salicylic acid biosynthetic process |
| 47 | 947591 | 'higB' | mRNA | 5.226803 | 1.04E-10 | 55.67 | 29.72 | 265.06 | 0.67 | 1.14 | 0.72 | | GO:00037 | GO:0017148///negative regulation of translation+++GO:0043488///regulation of mRNA stability+++GO:0090305///nucleic acid phosphodiester bond hydrolysis+++GO:0090502///RNA phosphodiester bond hydrolysis, endonucleolytic |
| 48 | 946511 | 'yecT' | mRNA | 5.162027 | 1.41E-07 | 18.6 | 9.85 | 75.97 | 0 | 0.21 | 0.67 | | | |
| 49 | 947426 | 'entE' | mRNA | 5.151465 | 6.11E-161 | 353.78 | 376.74 | 364.79 | 3.61 | 3.92 | 3.57 | GO:00058 | GO:00001 | GO:0009239///enterobactin biosynthetic process+++GO:0019290///siderophore biosynthetic process |
| 50 | 946088 | 'pinQ' | mRNA | 5.125165 | 6.93E-47 | 80.91 | 68.7 | 99.2 | 0.6 | 0.74 | 1.18 | | GO:00001 | GO:0006310///DNA recombination+++GO:0015074///DNA integration |
| 51 | 945981 | 'ydeM' | mRNA | 5.044166 | 1.48E-06 | 2.99 | 1.75 | 5.7 | 0.03 | 0 | 0.07 | | GO:00038 | GO:0006974///cellular response to DNA damage stimulus+++GO:0051604///protein maturation |
| 52 | 945201 | 'fepC' | mRNA | 4.969995 | 2.85E-79 | 108.59 | 111.98 | 116.34 | 1.34 | 1.15 | 1.32 | GO:00058 | GO:00001 | GO:0006811///ion transport+++GO:0015685///ferric-enterobactin transport+++GO:0055072///iron ion homeostasis |
| 53 | 949098 | 'yfdH' | mRNA | 4.964432 | 1.10E-20 | 97.86 | 70.82 | 305.03 | 1.16 | 1.76 | 2.07 | GO:00058 | GO:0005515///protein binding+++GO:0016740///transferase activity+++GO:0016757///transferase activity, transferring glycosyl groups | |
| 54 | 38094947 | 'ymcF' | mRNA | 4.91431 | 6.39E-07 | 317.42 | 479.71 | 155.5 | 0 | 8.45 | 2.05 | | | |
| 55 | 945846 | 'trpE' | mRNA | 4.827728 | 3.30E-70 | 189.87 | 196.54 | 198.06 | 2.41 | 3.32 | 1.7 | GO:00058 | GO:00038 | GO:0000162///tryptophan biosynthetic process+++GO:0008152///metabolic process+++GO:0008652///cellular amino acid biosynthetic process+++GO:0009058///biosynthetic process+++GO:0009073///aromatic amino acid family biosynthetic process |
| 56 | 946178 | 'entB' | mRNA | 4.786766 | 2.92E-73 | 263.88 | 327.61 | 444.52 | 4.27 | 3.6 | 5.15 | GO:00057 | GO:00002 | GO:0008152///metabolic process+++GO:0009239///enterobactin biosynthetic process |
| 57 | 946108 | 'ydfU' | mRNA | 4.761301 | 1.11E-28 | 16.23 | 14.86 | 19.82 | 0.23 | 0.23 | 0.2 | | | |
| 58 | 947145 | 'proW' | mRNA | 4.71814 | 1.28E-19 | 1668.99 | 2109.48 | 568.4 | 23.99 | 19.77 | 17.93 | GO:00058 | GO:00052 | GO:0006865///amino acid transport+++GO:0006972///hyperosmotic response+++GO:0015837///amine transport+++GO:0015871///choline transport+++GO:0031460///glycine betaine transport+++GO:0055085///transmembrane transport+++GO:1902603///carnitine transmembrane transport |
| 59 | 945922 | 'ydeO' | mRNA | 4.676866 | 1.89E-08 | 7.34 | 4.03 | 16.07 | 0.06 | 0.17 | 0.12 | | GO:00036 | GO:0006351///transcription, DNA-templated+++GO:0006355///regulation of transcription, DNA-templated+++GO:0045893///positive regulation of transcription, DNA-templated |
| 60 | 945242 | 'ypjB' | mRNA | 4.668485 | 0.001807 | 1.22 | 0.91 | 2.6 | 0.05 | 0 | 0 | | | |
| 61 | 945243 | 'ybeT' | mRNA | 4.589777 | 0.003236 | 2.16 | 0.95 | 4.39 | 0 | 0 | 0.09 | | | |
| 62 | 949009 | 'yafT' | mRNA | 4.582205 | 2.20E-23 | 37.06 | 22.18 | 50.69 | 0.27 | 0.66 | 0.63 | GO:0005886///plasma membrane+++GO:0016020///membrane | | |
| 63 | 945184 | 'entF' | mRNA | 4.571082 | 8.60E-46 | 241.52 | 257.52 | 149.72 | 3.3 | 3.52 | 3.23 | GO:00057 | GO:00001 | GO:0008152///metabolic process+++GO:0009058///biosynthetic process+++GO:0009239///enterobactin biosynthetic process+++GO:0043041///amino acid activation for nonribosomal peptide biosynthetic process+++GO:0044550///secondary metabolite biosynthetic process |
| 64 | 948352 | 'yihF' | mRNA | 4.555265 | 1.04E-22 | 17 | 13.27 | 33.45 | 0.19 | 0.4 | 0.34 | | | |
| 65 | 944837 | 'yadC' | mRNA | 4.547549 | 1.94E-05 | 2.86 | 1.06 | 17.43 | 0.13 | 0.03 | 0.13 | GO:0009289///pilus | | GO:0006974///cellular response to DNA damage stimulus+++GO:0007155///cell adhesion+++GO:0043709///cell adhesion involved in single-species biofilm formation+++GO:0043711///pilus organization+++GO:0046677///response to antibiotic |
| 66 | 946975 | 'purM' | mRNA | 4.541058 | 9.97E-21 | 327.18 | 404.26 | 135.77 | 5.01 | 5.4 | 3.47 | GO:00057 | GO:00001 | GO:0006164///purine nucleotide biosynthetic process+++GO:0006189///'de novo' IMP biosynthetic process+++GO:0046084///adenine biosynthetic process |
| 67 | 946006 | 'yncE' | mRNA | 4.527969 | 2.39E-172 | 1297.4 | 1220.53 | 1357.25 | 19.12 | 20.1 | 20.41 | GO:00302 | GO:0003677///DNA binding | |
| 68 | 947094 | 'yfiB' | mRNA | 4.510483 | 6.27E-10 | 142.07 | 68.08 | 828.6 | 4.33 | 4.63 | 4.63 | GO:0005886///plasma membrane+++GO:0009279///cell outer membrane+++GO:0016020///membrane+++GO:0016021///integral component of membrane | | |
| 69 | 947513 | 'ykgH' | mRNA | 4.492032 | 0.001387 | 1.94 | 1.33 | 7.07 | 0 | 0 | 0.14 | GO:0005886///plasma membrane+++GO:0016020///membrane+++GO:0016021///integral component of membrane | | |
| 70 | 945801 | 'ychO' | mRNA | 4.451117 | 4.37E-22 | 112.67 | 98.71 | 348.72 | 2.57 | 3.44 | 2.66 | GO:0009279///cell ou | | GO:0009405///pathogenesis |
| 71 | 946773 | 'elfD' | mRNA | 4.408428 | 3.41E-07 | 7.01 | 3.68 | 24.18 | 0.19 | 0.13 | 0.2 | GO:0030288///outer | | GO:0006457///protein folding+++GO:0043711///pilus organization+++GO:0061077///chaperone-mediated protein folding+++GO:0071555///cell wall organization |
| 72 | 948172 | 'xanP' | mRNA | 4.349084 | 2.41E-18 | 104.15 | 117.83 | 39.68 | 1.97 | 1.34 | 1.48 | GO:00058 | GO:00052 | GO:0042906///xanthine transport+++GO:0055085///transmembrane transport |
| 73 | 946650 | 'yehF' | mRNA | 4.33873 | 1.03E-05 | 3.53 | 2.43 | 20.75 | 0.2 | 0.16 | 0.05 | | | |
| 74 | 948235 | 'bglG' | mRNA | 4.336695 | 0.001476 | 1.71 | 0.94 | 3.79 | 0.1 | 0 | 0 | | GO:00037 | GO:0006355///regulation of transcription, DNA-templated+++GO:0045893///positive regulation of transcription, DNA-templated |
| 75 | 945591 | 'cspG' | mRNA | 4.31626 | 2.31E-19 | 2652.63 | 2422.35 | 2021.06 | 20.82 | 55.28 | 35.2 | GO:00057 | GO:00010 | GO:0009409///response to cold+++GO:0010468///regulation of gene expression+++GO:0060567///negative regulation of DNA-templated transcription, termination |
| 76 | 947344 | 'ygeQ' | mRNA | 4.314747 | 3.05E-13 | 24.05 | 13.64 | 59.52 | 0.32 | 0.6 | 0.68 | GO:00058 | GO:0003824///catalytic activity | |
| 77 | 948719 | 'yjfZ' | mRNA | 4.274078 | 8.01E-05 | 3.53 | 1.54 | 8.5 | 0 | 0.05 | 0.17 | GO:0016020///membrane+++GO:0016021///integral component of membrane | | |
| 78 | 1450243 | 'ybdZ' | mRNA | 4.266607 | 8.42E-19 | 147.6 | 128.88 | 279.52 | 2.73 | 2.81 | 2.43 | GO:00058 | GO:00055 | GO:0009239///enterobactin biosynthetic process+++GO:0019290///siderophore biosynthetic process+++GO:0043085///positive regulation of catalytic activity |
| 79 | 948432 | 'metF' | mRNA | 4.230716 | 2.80E-24 | 131.57 | 142.08 | 71.13 | 2.64 | 2.45 | 1.67 | GO:00058 | GO:00044 | GO:0006555///methionine metabolic process+++GO:0006730///one-carbon metabolic process+++GO:0008652///cellular amino acid biosynthetic process+++GO:0009086///methionine biosynthetic process+++GO:0035999///tetrahydrofolate interconversion+++GO:0046654///tetrahydrofolate biosynthetic process+++GO:0051289///protein homotetramerization+++GO:0055114///oxidation-reduction process |
| 80 | 946633 | 'gatA' | mRNA | 4.20865 | 1.06E-56 | 977.28 | 889.94 | 1107.54 | 13.41 | 23.41 | 17.61 | GO:00057 | GO:00163 | GO:0008643///carbohydrate transport+++GO:0009401///phosphoenolpyruvate-dependent sugar phosphotransferase system+++GO:0015796///galactitol transport+++GO:0016310///phosphorylation+++GO:0019402///galactitol metabolic process+++GO:0034219///carbohydrate transmembrane transport |
| 81 | 945215 | 'entH' | mRNA | 4.182018 | 3.22E-30 | 193.5 | 199.53 | 515.83 | 5.52 | 4.67 | 5.5 | GO:00057 | GO:00055 | GO:0009239///enterobactin biosynthetic process |
| 82 | 948148 | 'waaL' | mRNA | 4.118609 | 1.45E-05 | 14.13 | 6.33 | 128.45 | 0.4 | 1.22 | 1.11 | GO:00058 | GO:00167 | GO:0009103///lipopolysaccharide biosynthetic process+++GO:0009244///lipopolysaccharide core region biosynthetic process |
| 83 | 948821 | 'yfdG' | mRNA | 4.100039 | 6.41E-14 | 85.83 | 62.77 | 216.31 | 1.01 | 1.9 | 3.46 | GO:00058 | GO:00038 | GO:0000271///polysaccharide biosynthetic process+++GO:0006783///heme biosynthetic process+++GO:0055114///oxidation-reduction process |
| 84 | 946209 | 'ppsA' | mRNA | 4.06703 | 1.17E-32 | 499.3 | 572.36 | 314.77 | 11.93 | 10.33 | 8.34 | GO:00058 | GO:00001 | GO:0006090///pyruvate metabolic process+++GO:0006094///gluconeogenesis+++GO:0016310///phosphorylation |
| 85 | 946268 | 'entS' | mRNA | 4.064273 | 3.59E-25 | 115.95 | 138.65 | 72.05 | 1.89 | 2.77 | 2.49 | GO:00058 | GO:00429 | GO:0006974///cellular response to DNA damage stimulus+++GO:0015891///siderophore transport+++GO:0042930///enterobactin transport+++GO:0046677///response to antibiotic |
| 86 | 946642 | 'yehA' | mRNA | 4.058034 | 0.004945 | 1.23 | 0.23 | 4.98 | 0 | 0.08 | 0.04 | GO:0009289///pilus | | GO:0007155///cell adhesion+++GO:0043709///cell adhesion involved in single-species biofilm formation |
| 87 | 946610 | 'gatB' | mRNA | 3.998417 | 1.25E-37 | 429.65 | 310.46 | 665.45 | 6.04 | 11.54 | 8.98 | GO:00057 | GO:00089 | GO:0008643///carbohydrate transport+++GO:0009401///phosphoenolpyruvate-dependent sugar phosphotransferase system+++GO:0015796///galactitol transport+++GO:0019402///galactitol metabolic process+++GO:0034219///carbohydrate transmembrane transport |
| 88 | 945284 | 'entA' | mRNA | 3.98642 | 1.07E-54 | 206.03 | 251.24 | 221.15 | 5.26 | 5.87 | 4.11 | GO:00058 | GO:00055 | GO:0009239///enterobactin biosynthetic process+++GO:0019290///siderophore biosynthetic process+++GO:0051289///protein homotetramerization+++GO:0055114///oxidation-reduction process |
| 89 | 946664 | 'psuK' | mRNA | 3.966133 | 0.002642 | 1.65 | 0.95 | 1.18 | 0.04 | 0.04 | 0 | | GO:00001 | GO:0016310///phosphorylation+++GO:0046835///carbohydrate phosphorylation |
| 90 | 946686 | 'mepS' | mRNA | 3.949072 | 6.45E-47 | 713.36 | 633.83 | 1283.4 | 15.97 | 23.22 | 17.74 | GO:00092 | GO:00041 | GO:0000270///peptidoglycan metabolic process+++GO:0006508///proteolysis+++GO:0009254///peptidoglycan turnover+++GO:0045227///capsule polysaccharide biosynthetic process+++GO:0071555///cell wall organization |
| 91 | 944974 | 'rclC' | mRNA | 3.947406 | 1.04E-07 | 7.91 | 5.47 | 13.53 | 0.16 | 0.08 | 0.33 | GO:0005886///plasm | | GO:1901530///response to hypochlorite |
| 92 | 945181 | 'fes' | mRNA | 3.937424 | 2.03E-87 | 165.92 | 147.52 | 167.61 | 3.34 | 4.01 | 3.84 | GO:00057 | GO:00055 | GO:0006811///ion transport+++GO:0006826///iron ion transport+++GO:0033214///iron assimilation by chelation and transport+++GO:0046214///enterobactin catabolic process+++GO:0055072///iron ion homeostasis |
| 93 | 945696 | 'ycjN' | mRNA | 3.916436 | 9.48E-14 | 8.45 | 6.98 | 17.56 | 0.21 | 0.15 | 0.38 | GO:0042597///periplasmic space | | |
| 94 | 946196 | 'ydiM' | mRNA | 3.899964 | 4.06E-07 | 2.9 | 2.22 | 3.23 | 0.1 | 0 | 0.1 | GO:00058 | GO:00052 | GO:0055085///transmembrane transport |
| 95 | 944912 | 'yafP' | mRNA | 3.898705 | 3.21E-09 | 37.34 | 16.83 | 112 | 0.82 | 1.31 | 1.25 | | GO:00167 | GO:0006281///DNA repair+++GO:0006974///cellular response to DNA damage stimulus+++GO:0009432///SOS response |
| 96 | 946091 | 'cspB' | mRNA | 3.87168 | 6.01E-10 | 3136.14 | 4709.59 | 3462.28 | 27.89 | 151.54 | 62.18 | GO:00057 | GO:00010 | GO:0009409///response to cold+++GO:0010468///regulation of gene expression+++GO:0060567///negative regulation of DNA-templated transcription, termination |
| 97 | 945983 | 'ydbD' | mRNA | 3.848437 | 5.94E-28 | 8.56 | 6.75 | 11.75 | 0.19 | 0.25 | 0.22 | | | GO:0051595///response to methylglyoxal |
| 98 | 946598 | 'gatD' | mRNA | 3.84482 | 3.79E-50 | 363.22 | 312.46 | 282.79 | 6.95 | 8.93 | 8.05 | GO:00058 | GO:00082 | GO:0019402///galactitol metabolic process+++GO:0019404///galactitol catabolic process+++GO:0055114///oxidation-reduction process |
| 99 | 945155 | 'ybcI' | mRNA | 3.844717 | 1.16E-28 | 87.55 | 108.07 | 82.54 | 2.63 | 2.2 | 2.01 | GO:0005886///plasma membrane+++GO:0016020///membrane+++GO:0016021///integral component of membrane | | |
| 100 | 945216 | 'ybdO' | mRNA | 3.843313 | 1.92E-04 | 3.76 | 0.91 | 16.54 | 0.18 | 0.19 | 0.1 | | GO:00036 | GO:0006355///regulation of transcription, DNA-templated |
| 101 | 948081 | 'insK' | mRNA | 3.841251 | 9.99E-15 | 12.93 | 12.45 | 13.47 | 0.15 | 0.25 | 0.57 | | GO:00036 | GO:0006310///DNA recombination+++GO:0006974///cellular response to DNA damage stimulus+++GO:0015074///DNA integration+++GO:0032196///transposition |
| 102 | 946493 | 'yeeN' | mRNA | 3.825603 | 1.15E-22 | 184.43 | 126.62 | 353.3 | 3.65 | 6.5 | 5.5 | GO:00057 | GO:00036 | GO:0006355///regulation of transcription, DNA-templated |
| 103 | 946902 | 'iprA' | mRNA | 3.825344 | 0.048149 | 0.76 | 0.15 | 3.14 | 0 | 0.07 | 0 | | | GO:0006351///transcription, DNA-templated+++GO:0006979///response to oxidative stress |
| 104 | 1450297 | 'topAI' | mRNA | 3.814251 | 0.017014 | 3.84 | 1.78 | 87.85 | 0 | 1.07 | 0.38 | | GO:00055 | GO:0010911///regulation of isomerase activity |
| 105 | 946234 | 'katE' | mRNA | 3.811393 | 2.81E-55 | 2231.1 | 1918.69 | 3286.26 | 78.94 | 60.05 | 49.48 | GO:00057 | GO:00040 | GO:0006972///hyperosmotic response+++GO:0006974///cellular response to DNA damage stimulus+++GO:0006979///response to oxidative stress+++GO:0042744///hydrogen peroxide catabolic process+++GO:0055114///oxidation-reduction process+++GO:0098869///cellular oxidant detoxification |
| 106 | 946572 | 'yeaD' | mRNA | 3.806639 | 3.17E-50 | 199.22 | 171.76 | 334.45 | 5.65 | 6.55 | 5.15 | GO:00057 | GO:00038 | GO:0005975///carbohydrate metabolic process+++GO:0006974///cellular response to DNA damage stimulus |
| 107 | 945973 | 'ynbA' | mRNA | 3.796803 | 1.73E-04 | 6.03 | 2.96 | 2.73 | 0.15 | 0.08 | 0.08 | GO:00058 | GO:00084 | GO:0008654///phospholipid biosynthetic process |
| 108 | 949104 | 'fecR' | mRNA | 3.781223 | 7.86E-20 | 160.25 | 173.01 | 79.05 | 3.67 | 3.64 | 3.7 | GO:00425 | GO:00169 | GO:0006811///ion transport+++GO:0050896///response to stimulus+++GO:0055072///iron ion homeostasis+++GO:1903507///negative regulation of nucleic acid-templated transcription |
| 109 | 945153 | 'purK' | mRNA | 3.779187 | 1.01E-63 | 135.7 | 132.41 | 129.26 | 3.35 | 3.25 | 3.75 | GO:00058 | GO:00001 | GO:0006164///purine nucleotide biosynthetic process+++GO:0006189///'de novo' IMP biosynthetic process |
| 110 | 946612 | 'nrdA' | mRNA | 3.758 | 9.02E-76 | 187.16 | 172.16 | 285.68 | 5.54 | 5.94 | 5.38 | GO:00058 | GO:00001 | GO:0006260///DNA replication+++GO:0008152///metabolic process+++GO:0009263///deoxyribonucleotide biosynthetic process+++GO:0015949///nucleobase-containing small molecule interconversion+++GO:0055114///oxidation-reduction process |
| 111 | 945936 | 'ortT' | mRNA | 3.721387 | 0.001317 | 63.26 | 59.58 | 50.48 | 1.28 | 0 | 2.94 | GO:0005886///plasm | | GO:0001896///autolysis |
| 112 | 945239 | 'rlmH' | mRNA | 3.721035 | 1.23E-25 | 357.03 | 420.71 | 282.34 | 8.76 | 8.13 | 11.27 | GO:00057 | GO:00055 | GO:0006364///rRNA processing+++GO:0031167///rRNA methylation+++GO:0032259///methylation+++GO:0070475///rRNA base methylation |
| 113 | 949031 | 'purE' | mRNA | 3.713278 | 2.23E-25 | 138.31 | 139.61 | 97.37 | 3.7 | 3.57 | 2.91 | GO:00058 | GO:00168 | GO:0006164///purine nucleotide biosynthetic process+++GO:0006189///'de novo' IMP biosynthetic process |
| 114 | 945240 | 'mrdA' | mRNA | 3.696342 | 6.42E-88 | 195.7 | 179.71 | 277.76 | 5.69 | 6.29 | 5.83 | GO:00058 | GO:00041 | GO:0006508///proteolysis+++GO:0008360///regulation of cell shape+++GO:0009252///peptidoglycan biosynthetic process+++GO:0042493///response to drug+++GO:0046677///response to antibiotic+++GO:0071555///cell wall organization |
| 115 | 38094965 | 'yoaL' | mRNA | 3.690478 | 1.94E-06 | 82.36 | 45.18 | 224.43 | 1.25 | 0.64 | 4.87 | GO:0005886///plasma membrane+++GO:0016020///membrane+++GO:0016021///integral component of membrane | | |
| 116 | 945194 | 'entD' | mRNA | 3.690224 | 1.87E-18 | 31.12 | 23.88 | 57.92 | 0.96 | 0.75 | 1.18 | GO:00058 | GO:00002 | GO:0009237///siderophore metabolic process+++GO:0009239///enterobactin biosynthetic process |
| 117 | 947128 | 'yfjW' | mRNA | 3.68815 | 0.019629 | 1.37 | 0.84 | 30.95 | 0.09 | 0.38 | 0.35 | GO:0005886///plasma membrane+++GO:0016020///membrane+++GO:0016021///integral component of membrane | | |
| 118 | 947435 | 'ydcC' | mRNA | 3.687089 | 9.87E-14 | 11.16 | 8.59 | 22.82 | 0.26 | 0.36 | 0.52 | | GO:00036 | GO:0006313///transposition, DNA-mediated |
| 119 | 945180 | 'fepE' | mRNA | 3.662456 | 6.58E-07 | 4.25 | 4.27 | 2.34 | 0.1 | 0.04 | 0.18 | GO:00058 | GO:00047 | GO:0006811///ion transport+++GO:0009103///lipopolysaccharide biosynthetic process+++GO:0015685///ferric-enterobactin transport+++GO:0018108///peptidyl-tyrosine phosphorylation+++GO:0055072///iron ion homeostasis |
| 120 | 948832 | 'fimB' | mRNA | 3.642608 | 5.99E-20 | 34.75 | 24.56 | 55.62 | 1.07 | 1.25 | 0.74 | | GO:00036 | GO:0006310///DNA recombination+++GO:0006974///cellular response to DNA damage stimulus+++GO:0015074///DNA integration |
| 121 | 947704 | 'envR' | mRNA | 3.630862 | 5.13E-04 | 5.33 | 4.44 | 1.43 | 0.2 | 0 | 0.14 | | GO:00036 | GO:0042493///response to drug+++GO:0045892///negative regulation of transcription, DNA-templated |
| 122 | 946553 | 'wzzB' | mRNA | 3.623316 | 3.12E-50 | 720.29 | 719.35 | 927.93 | 16.98 | 25.91 | 24.41 | GO:00058 | GO:00047 | GO:0009103///lipopolysaccharide biosynthetic process+++GO:0018108///peptidyl-tyrosine phosphorylation |
| 123 | 945274 | 'ybfP' | mRNA | 3.622944 | 3.70E-05 | 5.77 | 4.07 | 15.94 | 0.2 | 0.1 | 0.33 | GO:0005886///plasma membrane+++GO:0016020///membrane | | |
| 124 | 946438 | 'ampG' | mRNA | 3.620512 | 2.10E-50 | 51.5 | 54.05 | 64.53 | 1.41 | 2.14 | 1.38 | GO:00058 | GO:00152 | GO:0000270///peptidoglycan metabolic process+++GO:0015835///peptidoglycan transport+++GO:0055085///transmembrane transport+++GO:1902600///proton transmembrane transport |
| 125 | 945592 | 'ycdU' | mRNA | 3.612012 | 3.35E-11 | 7.37 | 6.34 | 5.84 | 0.29 | 0.12 | 0.17 | GO:0005886///plasma membrane+++GO:0016020///membrane+++GO:0016021///integral component of membrane | | |
| 126 | 945367 | 'sfmC' | mRNA | 3.596684 | 0.032014 | 0.53 | 0.76 | 1.34 | 0 | 0.06 | 0 | GO:0030288///outer | | GO:0006457///protein folding+++GO:0043711///pilus organization+++GO:0061077///chaperone-mediated protein folding+++GO:0071555///cell wall organization |
| 127 | 945157 | 'ygiA' | mRNA | 3.596171 | 3.70E-07 | 99.08 | 64.82 | 661.81 | 5.73 | 5.89 | 5.18 | | | |
| 128 | 946752 | 'yfbO' | mRNA | 3.590098 | 9.09E-04 | 4.95 | 2.87 | 14.16 | 0.26 | 0.27 | 0 | | | |
| 129 | 948663 | 'yieJ' | mRNA | 3.587588 | 9.93E-11 | 11.42 | 9 | 9.5 | 0.19 | 0.15 | 0.56 | | | |

| | | | | | | | | | | | | | | |
|---|---|---|---|---|---|---|---|---|---|---|---|---|---|---|
| 130 | 945711 | 'csgE' | mRNA | 3.585415 | 3.73E-04 | 8.71 | 4.8 | 8.33 | 0.45 | 0.15 | 0 | GO:00092 | GO:00055 | GO:0022610///biological adhesion+++GO:0044010///single-species biofilm formation+++GO:0071806///protein transmembrane transport |
| 131 | 946490 | 'torY' | mRNA | 3.570535 | 6.06E-12 | 6.91 | 5.92 | 13.1 | 0.14 | 0.36 | 0.23 | GO:00058 | GO:00055 | GO:0009061///anaerobic respiration+++GO:0019645///anaerobic electron transport chain+++GO:0022900///electron transport chain+++GO:0055114///oxidation-reduction process |
| 132 | 947741 | 'yhcA' | mRNA | 3.568781 | 0.002256 | 2.19 | 1.71 | 3.26 | 0 | 0.2 | 0 | GO:0016020///memb | | GO:0071555///cell wall organization |
| 133 | 947538 | 'fepB' | mRNA | 3.566983 | 6.42E-43 | 166.28 | 151.98 | 244.87 | 5.35 | 4.44 | 6.66 | GO:0030288///outer | | GO:0006811///ion transport+++GO:0006974///cellular response to DNA damage stimulus+++GO:0015685///ferric-enterobactin transport+++GO:0042930///enterobactin transport+++GO:0055072///iron ion homeostasis |
| 134 | 945209 | 'fepG' | mRNA | 3.565618 | 4.79E-42 | 57.05 | 65.11 | 56.88 | 1.91 | 1.86 | 1.68 | GO:00058 | GO:00052 | GO:0006811///ion transport+++GO:0015685///ferric-enterobactin transport+++GO:0033214///iron assimilation by chelation and transport+++GO:0055072///iron ion homeostasis |
| 135 | 946148 | 'rsxA' | mRNA | 3.5501 | 3.49E-38 | 194.35 | 179.14 | 290.89 | 4.59 | 7.54 | 7.04 | GO:0005886///plasma | | GO:0022900///electron transport chain+++GO:0055114///oxidation-reduction process |
| 136 | 945193 | 'fepA' | mRNA | 3.546303 | 1.76E-41 | 99.14 | 85.84 | 172.76 | 3.65 | 3.67 | 3.42 | GO:00092 | GO:00055 | GO:0006811///ion transport+++GO:0015685///ferric-enterobactin transport+++GO:0015891///siderophore transport+++GO:0033214///iron assimilation by chelation and transport+++GO:0042914///colicin transport+++GO:0042930///enterobactin transport+++GO:0044718///siderophore transmembrane transport+++GO:0055072///iron ion homeostasis+++GO:0055085///transmembrane transport |
| 137 | 946090 | 'cspF' | mRNA | 3.543558 | 3.11E-08 | 144.81 | 134.23 | 106.02 | 1.78 | 6.14 | 2.66 | GO:00057 | GO:00010 | GO:0010468///regulation of gene expression+++GO:0060567///negative regulation of DNA-templated transcription, termination |
| 138 | 945521 | 'yajG' | mRNA | 3.529555 | 6.08E-20 | 139.27 | 93.38 | 276.25 | 3.73 | 5.78 | 4.93 | GO:0005886///plasma membrane+++GO:0016020///membrane | | |
| 139 | 946847 | 'lpxP' | mRNA | 3.511436 | 6.96E-44 | 633.36 | 598.42 | 614.32 | 16.95 | 25.12 | 15.66 | GO:00058 | GO:00089 | GO:0009103///lipopolysaccharide biosynthetic process+++GO:0009245///lipid A biosynthetic process+++GO:0009409///response to cold+++GO:0036104///Kdo2-lipid A biosynthetic process |
| 140 | 948527 | 'yjbH' | mRNA | 3.507974 | 4.48E-37 | 278.87 | 277.52 | 197.98 | 8.39 | 7.6 | 8.2 | GO:0005886///plasma | | GO:0045226///extracellular polysaccharide biosynthetic process |
| 141 | 945644 | 'ydjM' | mRNA | 3.505068 | 1.77E-18 | 152.11 | 113.31 | 89.18 | 2.68 | 3.77 | 4.62 | GO:0005886///plasma | | GO:0006974///cellular response to DNA damage stimulus |
| 142 | 946011 | 'paaF' | mRNA | 3.502765 | 2.91E-26 | 27.97 | 29.39 | 46.13 | 1.39 | 0.96 | 0.77 | | GO:00038 | GO:0006629///lipid metabolic process+++GO:0006631///fatty acid metabolic process+++GO:0006635///fatty acid beta-oxidation+++GO:0010124///phenylacetate catabolic process |
| 143 | 945851 | 'add' | mRNA | 3.483457 | 7.13E-41 | 342.78 | 287.64 | 254.69 | 10.49 | 10.24 | 7.83 | GO:00058 | GO:00040 | GO:0006154///adenosine catabolic process+++GO:0006974///cellular response to DNA damage stimulus+++GO:0009117///nucleotide metabolic process+++GO:0009168///purine ribonucleoside monophosphate biosynthetic process+++GO:0015950///purine nucleotide interconversion+++GO:0032261///purine nucleotide salvage+++GO:0043103///hypoxanthine salvage+++GO:0046101///hypoxanthine biosynthetic process+++GO:0046103///inosine biosynthetic process |
| 144 | 948140 | 'waaH' | mRNA | 3.480522 | 2.60E-22 | 22.16 | 18.84 | 39.57 | 0.7 | 0.86 | 0.9 | GO:00160 | GO:00150 | GO:0016036///cellular response to phosphate starvation |
| 145 | 945961 | 'yddB' | mRNA | 3.465073 | 1.03E-18 | 5.86 | 4.66 | 9.89 | 0.23 | 0.2 | 0.21 | GO:0009279///cell outer membrane | | |
| 146 | 945237 | 'rsfS' | mRNA | 3.458871 | 1.50E-19 | 420.31 | 504.89 | 347.65 | 12.4 | 16.26 | 10.62 | GO:00057 | GO:00055 | GO:0006417///regulation of translation+++GO:0017148///negative regulation of translation+++GO:0042256///mature ribosome assembly+++GO:0090071///negative regulation of ribosome biogenesis |
| 147 | 948063 | 'dppB' | mRNA | 3.458626 | 4.19E-11 | 40.16 | 50.44 | 16.55 | 1.49 | 1.24 | 0.92 | GO:00058 | GO:00719 | GO:0015031///protein transport+++GO:0015833///peptide transport+++GO:0015886///heme transport+++GO:0035442///dipeptide transmembrane transport+++GO:0055085///transmembrane transport |
| 148 | 948310 | 'metR' | mRNA | 3.446666 | 3.63E-15 | 92.57 | 88.83 | 40.69 | 2.39 | 2.81 | 2.3 | GO:00057 | GO:00009 | GO:0006355///regulation of transcription, DNA-templated+++GO:0008652///cellular amino acid biosynthetic process+++GO:0009086///methionine biosynthetic process |
| 149 | 947149 | 'nrdF' | mRNA | 3.443391 | 4.33E-63 | 69.51 | 67.12 | 77.26 | 2.33 | 2.32 | 2.33 | GO:00057 | GO:00047 | GO:0006260///DNA replication+++GO:0009263///deoxyribonucleotide biosynthetic process+++GO:0055114///oxidation-reduction process |
| 150 | 945219 | 'ybeF' | mRNA | 3.443108 | 6.67E-06 | 7.95 | 3.94 | 33.13 | 0.34 | 0.39 | 0.59 | | GO:00036 | GO:0006351///transcription, DNA-templated+++GO:0006355///regulation of transcription, DNA-templated |
