## Supplementary Data 3 for "Enhanced extracellular matrix production provides protection to cell wall-deficient *Escherichia coli*"

Supplementary Data 3. The 150 most downregulated DEGs of the L-form compared to wild type.

| | Gene ID | Gene Symbol | Type | log2 (L-form / Wild- | Qvalue (L-form / Wild- | Lform_1 TPM | Lform_2 TPM | Lform_3 TPM | WT_1 TPM | WT_2 TPM | WT_3 TPM | GO_C Desc | GO_F Desc | GO_P Desc |
|---|---|---|---|---|---|---|---|---|---|---|---|---|---|---|
| 1 | 946315 | 'ldhA' | mRNA | -13.53 | 2.27E-60 | 0 | 0.12 | 0 | 264.94 | 263.07 | 256.92 | GO:0005 | GO:0008 | GO:0009408///response to heat+++GO:0019664///mixed acid fermentation+++GO:0055114///oxidation-reduction process |
| 2 | 946587 | 'ydbK' | mRNA | -13.133 | ######## | 0.01 | 0.11 | 0.13 | 280.18 | 221.07 | 182.61 | | GO:0003 | GO:0006979///response to oxidative stress+++GO:0022900///electron transport chain+++GO:0055114///oxidation-reduction process |
| 3 | 945948 | 'uspF' | mRNA | -12.717 | 1.05E-28 | 0.13 | 0 | 0 | 172.32 | 215.39 | 212.92 | | GO:0005 | GO:0006979///response to oxidative stress+++GO:0007155///cell adhesion |
| 4 | 4056027 | 'ydbJ' | mRNA | -10.979 | 4.79E-19 | 0 | 0 | 0 | 85.19 | 83.86 | 75.12 | | | |
| 5 | 945898 | 'ynaI' | mRNA | -10.963 | 2.92E-21 | 0 | 0.04 | 0 | 14.93 | 22.1 | 18.29 | GO:0005 | GO:0008 | GO:0006811///ion transport+++GO:0055085///transmembrane transport+++GO:0071470///cellular response to osmotic stress |
| 6 | 945944 | 'insH-5' | mRNA | -10.848 | 1.92E-18 | 0 | 0 | 0 | 8.84 | 9.36 | 10.17 | GO:0005 | GO:0003 | GO:0006310///DNA recombination+++GO:0006313///transposition, DNA-mediated+++GO:0032196///transposition+++GO:0045893///positive regulation of transcription, DNA-templated |
| 7 | 945933 | 'feaB' | mRNA | -10.434 | 1.28E-27 | 0 | 0.05 | 0 | 14.46 | 14.11 | 11.72 | | GO:0008 | GO:0006559///L-phenylalanine catabolic process+++GO:0009435///NAD biosynthetic process+++GO:0019607///phenylethylamine catabolic process+++GO:0046196///4-nitrophenol catabolic process+++GO:0051289///protein homotetramerization+++GO:0055114///oxidation-reduction process |
| 8 | 945207 | 'stfR' | mRNA | -10.365 | 6.21E-17 | 0 | 0 | 0 | 1.77 | 1.64 | 1.76 | | GO:0005198///structural molecule activity | |
| 9 | 945899 | 'racR' | mRNA | -10.166 | 3.16E-18 | 0 | 0.11 | 0 | 22.71 | 33.01 | 32.82 | | GO:0001 | GO:2000143///negative regulation of DNA-templated transcription, initiation |
| 10 | 946525 | 'hslJ' | mRNA | -9.6823 | 1.14E-23 | 0 | 0.26 | 0 | 40.32 | 46.3 | 37.55 | GO:0016020///mem | | GO:0009408///response to heat |
| 11 | 945918 | 'recE' | mRNA | -9.5048 | 6.74E-14 | 0 | 0 | 0 | 0.99 | 1.36 | 1.37 | | GO:0004 | GO:0006259///DNA metabolic process+++GO:0090305///nucleic acid phosphodiester bond hydrolysis |
| 12 | 948967 | 'ttcA' | mRNA | -9.3616 | 1.04E-15 | 0 | 0.04 | 0 | 5.71 | 7.71 | 7.07 | GO:0005 | GO:0000 | GO:0006400///tRNA modification+++GO:0008033///tRNA processing+++GO:0034227///tRNA thio-modification |
| 13 | 945939 | 'tynA' | mRNA | -9.2056 | 3.66E-13 | 0 | 0 | 0 | 1.05 | 1.15 | 1.3 | GO:0042 | GO:0005 | GO:0006559///L-phenylalanine catabolic process+++GO:0009308///amine metabolic process+++GO:0019607///phenylethylamine catabolic process+++GO:0055114///oxidation-reduction process |
| 14 | 945927 | 'ydaW' | mRNA | -8.9728 | 1.38E-12 | 0 | 0 | 0 | 5.01 | 5.7 | 5.25 | | GO:0005515///protein binding | |
| 15 | 945925 | 'ydaU' | mRNA | -8.9232 | 1.99E-12 | 0 | 0 | 0 | 3.2 | 2.66 | 2.93 | | | |
| 16 | 945926 | 'ydaV' | mRNA | -8.704 | 1.08E-11 | 0 | 0 | 0 | 3.7 | 2.76 | 2.58 | | GO:0005 | GO:0006260///DNA replication+++GO:0006271///DNA strand elongation involved in DNA replication |
| 17 | 945917 | 'recT' | mRNA | -8.1694 | 2.93E-10 | 0 | 0 | 0 | 1.77 | 2.07 | 1.77 | GO:0032 | GO:0003 | GO:0006259///DNA metabolic process+++GO:0006310///DNA recombination+++GO:0032508///DNA duplex unwinding+++GO:0043150///DNA synthesis involved in double-strand break repair via homologous recombination |
| 18 | 945949 | 'ydbH' | mRNA | -8.1109 | 6.20E-45 | 0.04 | 0.07 | 0 | 5.95 | 5.29 | 4.52 | GO:0005886///plasma membrane | | |
| 19 | 946057 | 'gadC' | mRNA | -8.0921 | 0 | 225.8 | 257.08 | 237.95 | 23084.9 | 24033.3 | 23620.7 | GO:0005 | GO:0015 | GO:0006865///amino acid transport+++GO:0051454///intracellular pH elevation+++GO:0055085///transmembrane transport |
| 20 | 945581 | 'hyaC' | mRNA | -8.0677 | ######## | 3.79 | 4.01 | 5.65 | 395.83 | 410.23 | 406.66 | GO:0005 | GO:0005 | GO:0006113///fermentation+++GO:0009061///anaerobic respiration+++GO:0022904///respiratory electron transport chain+++GO:0055114///oxidation-reduction process |
| 21 | 945575 | 'hyaD' | mRNA | -8.0286 | ######## | 7.45 | 9.59 | 10.31 | 808.83 | 802.39 | 880.19 | | GO:0004 | GO:0006464///cellular protein modification process+++GO:0006508///proteolysis+++GO:0016485///protein processing+++GO:0043085///positive regulation of catalytic activity |
| 22 | 945924 | 'ydaT' | mRNA | -7.9345 | 3.04E-09 | 0 | 0 | 0 | 5.59 | 3.05 | 3.35 | | | GO:0043093///FtsZ-dependent cytokinesis |
| 23 | 947997 | 'yhiM' | mRNA | -7.9252 | ######## | 17.77 | 13.76 | 13.59 | 1345.22 | 1456.04 | 1214.11 | GO:0005886///plas | | GO:0071229///cellular response to acid chemical |
| 24 | 945951 | 'mppA' | mRNA | -7.7041 | 3.47E-27 | 0.07 | 0.05 | 0 | 3.88 | 4.62 | 4.2 | GO:0030 | GO:1900 | GO:0015031///protein transport+++GO:0015833///peptide transport+++GO:0015834///peptidoglycan-associated peptide transport+++GO:0042939///tripeptide transport+++GO:0055085///transmembrane transport |
| 25 | 948027 | 'gadA' | mRNA | -7.6491 | ######## | 265.89 | 275.42 | 162.69 | 17321.1 | 16397.2 | 17896.9 | GO:0005 | GO:0003 | GO:0006536///glutamate metabolic process+++GO:0006538///glutamate catabolic process+++GO:0019752///carboxylic acid metabolic process+++GO:0051454///intracellular pH elevation |
| 26 | 945573 | 'hyaE' | mRNA | -7.5852 | ######## | 10.19 | 11.67 | 5.68 | 653.44 | 698.02 | 712.5 | | GO:0005048///signal sequence binding | |
| 27 | 946976 | 'intR' | mRNA | -7.4556 | 7.40E-10 | 0.03 | 0 | 0 | 1.19 | 1.24 | 1.5 | | GO:0003 | GO:0006310///DNA recombination+++GO:0015074///DNA integration+++GO:0044826///viral genome integration into host DNA+++GO:0046718///viral entry into host cell+++GO:0075713///establishment of integrated proviral latency |
| 28 | 946062 | 'tfaR' | mRNA | -7.4079 | 9.40E-08 | 0 | 0 | 0 | 1.21 | 1.79 | 2.32 | | | |
| 29 | 945941 | 'feaR' | mRNA | -7.3419 | 1.26E-09 | 0 | 0.05 | 0 | 1.82 | 1.98 | 1.49 | | GO:0000 | GO:0006355///regulation of transcription, DNA-templated+++GO:0045893///positive regulation of transcription, DNA-templated |
| 30 | 945466 | 'bssR' | mRNA | -7.3205 | ######## | 10.59 | 15.1 | 18.43 | 836.3 | 706.32 | 775.31 | | | GO:0010468///regulation of gene expression+++GO:1900190///regulation of single-species biofilm formation |
| 31 | 945580 | 'hyaB' | mRNA | -7.305 | 0 | 6.89 | 7.57 | 8.05 | 483.08 | 439.91 | 367.12 | GO:0005 | GO:0005 | GO:0006113///fermentation+++GO:0009061///anaerobic respiration+++GO:0022900///electron transport chain+++GO:0055114///oxidation-reduction process |
| 32 | 945954 | 'paaZ' | mRNA | -7.0943 | 2.75E-07 | 0 | 0 | 0 | 0.38 | 0.25 | 0.28 | | GO:0003 | GO:0008152///metabolic process+++GO:0010124///phenylacetate catabolic process+++GO:0055114///oxidation-reduction process |
| 33 | 945932 | 'trkG' | mRNA | -7.0691 | 3.48E-14 | 0.05 | 0.03 | 0 | 1.27 | 2.06 | 2.28 | GO:0005 | GO:0005 | GO:0006811///ion transport+++GO:0006812///cation transport+++GO:0006813///potassium ion transport+++GO:0055085///transmembrane transport+++GO:0071805///potassium ion transmembrane transport |
| 34 | 945923 | 'ydaS' | mRNA | -7.0669 | 5.03E-07 | 0 | 0 | 0 | 3.26 | 5.91 | 3.55 | | GO:0003 | GO:0008219///cell death |
| 35 | 945572 | 'hyaF' | mRNA | -7.0398 | ######## | 6.61 | 7.47 | 12.01 | 355.54 | 389.96 | 394.27 | | | |
| 36 | 947547 | 'appB' | mRNA | -6.9932 | 1.25E-50 | 9.67 | 12.94 | 3.5 | 417.06 | 458.81 | 393.69 | GO:0005 | GO:0009 | GO:0006119///oxidative phosphorylation+++GO:0019646///aerobic electron transport chain+++GO:0055114///oxidation-reduction process |
| 37 | 3.8E+07 | 'yecU' | mRNA | -6.8633 | 1.89E-05 | 0 | 0 | 0 | 4.75 | 17.24 | 31.22 | | | |
| 38 | 945959 | 'paaD' | mRNA | -6.749 | 4.82E-06 | 0 | 0 | 0 | 2.12 | 1.23 | 0.75 | | | GO:0010124///phenylacetate catabolic process |
| 39 | 945101 | 'dps' | mRNA | -6.7117 | 2.35E-87 | 2654.68 | 1822.13 | 5154.95 | 89990.9 | 108541 | 128367 | GO:0005 | GO:0003 | GO:0006879///cellular iron ion homeostasis+++GO:0030261///chromosome condensation+++GO:0042594///response to starvation+++GO:0055114///oxidation-reduction process |
| 40 | 945579 | 'hyaA' | mRNA | -6.6812 | ######## | 11.16 | 9.93 | 23.54 | 545.34 | 552.56 | 440.2 | GO:0005 | GO:0005 | GO:0006113///fermentation+++GO:0009061///anaerobic respiration+++GO:0022900///electron transport chain+++GO:0055114///oxidation-reduction process |
| 41 | 946058 | 'gadB' | mRNA | -6.6673 | 6.44E-74 | 405.58 | 483.83 | 231.82 | 13998.8 | 13870.7 | 14018.5 | GO:0005 | GO:0003 | GO:0006536///glutamate metabolic process+++GO:0006538///glutamate catabolic process+++GO:0019752///carboxylic acid metabolic process+++GO:0051454///intracellular pH elevation |
| 42 | 948638 | 'adiA' | mRNA | -6.6609 | ######## | 3.61 | 2.99 | 6.4 | 117.58 | 187.45 | 143.36 | GO:0005 | GO:0003 | GO:0006520///cellular amino acid metabolic process+++GO:0006527///arginine catabolic process+++GO:0051454///intracellular pH elevation |
| 43 | 945597 | 'cbpM' | mRNA | -6.6507 | 2.25E-22 | 134.62 | 174.24 | 22.91 | 4885.53 | 3861.24 | 3499.45 | | GO:0004 | GO:0043086///negative regulation of catalytic activity |
| 44 | 945585 | 'appC' | mRNA | -6.5022 | ######## | 5.2 | 6.11 | 6.8 | 195.21 | 212.12 | 179.4 | GO:0005 | GO:0009 | GO:0006119///oxidative phosphorylation+++GO:0019646///aerobic electron transport chain+++GO:0055114///oxidation-reduction process |
| 45 | 945103 | 'asr' | mRNA | -6.4938 | 3.80E-06 | 6.25 | 0.94 | 83.29 | 462.29 | 695.39 | 821.66 | GO:0042597///peri | | GO:0001101///response to acid chemical+++GO:0010043///response to zinc ion |
| 46 | 948710 | 'aidB' | mRNA | -6.4231 | ######## | 24.01 | 26.84 | 39.22 | 1111.52 | 937.13 | 686.4 | GO:0005 | GO:0003 | GO:0006974///cellular response to DNA damage stimulus+++GO:0045892///negative regulation of transcription, DNA-templated+++GO:0055114///oxidation-reduction process |
| 47 | 946313 | 'ompN' | mRNA | -6.1982 | 4.13E-05 | 0 | 0 | 0 | 0.31 | 0.42 | 0.22 | GO:0009 | GO:0015 | GO:0006811///ion transport+++GO:0034219///carbohydrate transmembrane transport+++GO:0055085///transmembrane transport |
| 48 | 947543 | 'ynaA' | mRNA | -6.1654 | 5.30E-05 | 0 | 0 | 0 | 0.23 | 0.43 | 0.37 | | | |
| 49 | 946206 | 'appA' | mRNA | -6.133 | ######## | 9.8 | 9.7 | 11.71 | 242.55 | 262.46 | 272.38 | GO:0030 | GO:0003 | GO:0008152///metabolic process+++GO:0016036///cellular response to phosphate starvation+++GO:0016311///dephosphorylation+++GO:0033518///myo-inositol hexakisphosphate dephosphorylation+++GO:0071454///cellular response to anoxia |
| 50 | 947095 | 'yegP' | mRNA | -6.0946 | ######## | 158.46 | 125.71 | 194.24 | 3436.87 | 3268.5 | 3807.99 | | | GO:0006302///double-strand break repair |
| 51 | 945956 | 'paaC' | mRNA | -6.0903 | 6.50E-05 | 0 | 0 | 0 | 0.46 | 0.65 | 0.37 | GO:0005 | GO:0005 | GO:0010124///phenylacetate catabolic process |
| 52 | 945401 | 'ybhL' | mRNA | -6.0784 | ######## | 40.78 | 37.06 | 32.78 | 980.38 | 963.05 | 762.35 | GO:0005886///plasma membrane+++GO:0016020///membrane+++GO:0016021///integral component of membrane | | |
| 53 | 947910 | 'yrbL' | mRNA | -6.0607 | 2.82E-35 | 111.75 | 117.88 | 38.88 | 2671.25 | 2211.19 | 1758.59 | | | |
| 54 | 948930 | 'yccJ' | mRNA | -5.9459 | ######## | 501.23 | 437.85 | 614.26 | 9219.68 | 8648 | 10894.6 | GO:0005829///cytosol | | |
| 55 | 948628 | 'adiC' | mRNA | -5.7999 | ######## | 5.83 | 6.03 | 7.66 | 119.31 | 170.49 | 95.88 | GO:0005 | GO:0015 | GO:0003333///amino acid transmembrane transport+++GO:0006865///amino acid transport+++GO:0051454///intracellular pH elevation+++GO:0055085///transmembrane transport+++GO:1903826///arginine transmembrane transport |
| 56 | 944953 | 'sodB' | mRNA | -5.7881 | ######## | 551.14 | 393.01 | 805.76 | 10206.9 | 11068.5 | 11209.7 | GO:0005 | GO:0004 | GO:0000303///response to superoxide+++GO:0006801///superoxide metabolic process+++GO:0019430///removal of superoxide radicals+++GO:0055114///oxidation-reduction process |
| 57 | 947572 | 'cbpA' | mRNA | -5.7801 | ######## | 131.65 | 122.5 | 151.75 | 3019.49 | 2641.16 | 2236.82 | GO:0005 | GO:0003 | GO:0006457///protein folding+++GO:0042026///protein refolding+++GO:0051085///chaperone cofactor-dependent protein refolding |
| 58 | 945913 | 'sieB' | mRNA | -5.7291 | 4.62E-04 | 0 | 0 | 0 | 0.52 | 0.53 | 1 | GO:0005886///plasma membrane+++GO:0016020///membrane+++GO:0016021///integral component of membrane | | |
| 59 | 946481 | 'hchA' | mRNA | -5.6992 | ######## | 253.99 | 196.69 | 404.22 | 4732.31 | 4891.49 | 5556.83 | GO:0005 | GO:0005 | GO:0006281///DNA repair+++GO:0006974///cellular response to DNA damage stimulus+++GO:0010447///response to acidic pH+++GO:0019243///methylglyoxal catabolic process to D-lactate via S-lactoyl-glutathione+++GO:0019249///lactate biosynthetic process+++GO:0030091///protein repair+++GO:0036525///protein deglycation+++GO:0042245///RNA repair+++GO:0051595///response to methylglyoxal+++GO:0106044///guanine deglycation |
| 60 | 945598 | 'ymdF' | mRNA | -5.663 | ######## | 2037.87 | 1575.45 | 3987.54 | 31112.7 | 30774.5 | 35408.9 | | | |
| 61 | 946051 | 'ygiW' | mRNA | -5.6194 | ######## | 425.36 | 351.57 | 844.5 | 8388.97 | 7953.85 | 8783.53 | GO:0030288///oute | | GO:0044011///single-species biofilm formation on inanimate substrate+++GO:0070301///cellular response to hydrogen peroxide+++GO:0071276///cellular response to cadmium ion |
| 62 | 948415 | 'uspD' | mRNA | -5.5656 | ######## | 115.8 | 100.42 | 196.89 | 1937.58 | 2130.93 | 2230.38 | GO:0005737///cyto | | GO:0000303///response to superoxide+++GO:0009411///response to UV+++GO:0042594///response to starvation |
| 63 | 945943 | 'ydbL' | mRNA | -5.5366 | 7.10E-05 | 0.22 | 0 | 0 | 2.47 | 3.16 | 1.34 | | | |
| 64 | 947263 | 'wrbA' | mRNA | -5.5294 | 6.12E-28 | 266.6 | 192.81 | 934.44 | 6119.44 | 6221.02 | 8105.18 | GO:0005 | GO:0000 | GO:0006979///response to oxidative stress+++GO:0051289///protein homotetramerization+++GO:0055114///oxidation-reduction process |
| 65 | 948736 | 'ytfK' | mRNA | -5.4553 | 2.30E-25 | 126.88 | 173.38 | 52.07 | 1670.95 | 1789.76 | 1907.81 | | | GO:0006979///response to oxidative stress |
| 66 | 948416 | 'yiiS' | mRNA | -5.4084 | 3.55E-69 | 259.64 | 242.81 | 208.39 | 2978.59 | 3283.12 | 3779.3 | GO:0005829///cytosol | | |
| 67 | 945833 | 'paaA' | mRNA | -5.3694 | 0.00267 | 0 | 0 | 0 | 0.31 | 0.09 | 0.28 | GO:0005 | GO:0005 | GO:0010124///phenylacetate catabolic process+++GO:0055114///oxidation-reduction process |
| 68 | 948025 | 'hdeA' | mRNA | -5.3069 | ######## | 2694.04 | 1947.38 | 2754.79 | 33435.7 | 33774.1 | 28126.1 | GO:0030 | GO:0042 | GO:0071468///cellular response to acidic pH+++GO:1990451///cellular stress response to acidic pH |
| 69 | 948022 | 'slp' | mRNA | -5.2913 | 1.25E-54 | 297.43 | 224.93 | 663.83 | 4605.66 | 5485.96 | 5083.97 | GO:0009 | GO:0042802///identical protein binding | |
| 70 | 944975 | 'yahK' | mRNA | -5.243 | ######## | 61.7 | 61.81 | 51.22 | 833.57 | 767.32 | 793.68 | | GO:0008 | GO:0055114///oxidation-reduction process |
| 71 | 947452 | 'ybfA' | mRNA | -5.2083 | 2.72E-47 | 518.7 | 580.68 | 453.72 | 5307.45 | 6121.09 | 6554.45 | GO:0005887///inte | | GO:0009314///response to radiation |
| 72 | 948507 | 'zraP' | mRNA | -5.1925 | 7.08E-69 | 4.49 | 4.26 | 4.99 | 51.05 | 66.1 | 58.23 | GO:0030 | GO:0005 | GO:0036460///cellular response to cell envelope stress |
| 73 | 945035 | 'yahH' | mRNA | -5.0968 | 0.01264 | 0 | 0 | 0 | 0.75 | 0.26 | 2.18 | | | |
| 74 | 949125 | 'yohK' | mRNA | -5.0612 | ######## | 22.91 | 30.06 | 28.94 | 338.93 | 341.57 | 291.62 | GO:0005886///plasma membrane+++GO:0016020///membrane+++GO:0016021///integral component of membrane | | |
| 75 | 948973 | 'pinR' | mRNA | -5.0308 | 2.71E-05 | 0.11 | 0 | 0.78 | 1.05 | 1.19 | 1.09 | | GO:0000 | GO:0006310///DNA recombination+++GO:0015074///DNA integration |
| 76 | 945492 | 'bhsA' | mRNA | -5.011 | ######## | 241.06 | 171.28 | 442.63 | 2536.34 | 2400.47 | 2918.27 | GO:0009279///cell | | GO:0042710///biofilm formation+++GO:0046688///response to copper ion |
| 77 | 946931 | 'yffR' | mRNA | -5.0038 | ######## | 20.47 | 18.5 | 33.97 | 227.37 | 239.73 | 265.43 | | | |
| 78 | 949127 | 'yohJ' | mRNA | -4.9631 | 4.12E-87 | 45.03 | 50.02 | 43.15 | 524.58 | 569.74 | 418.96 | GO:0005886///plasma membrane+++GO:0016020///membrane+++GO:0016021///integral component of membrane | | |
| 79 | 947144 | 'kbp' | mRNA | -4.8902 | ######## | 437.39 | 399.63 | 595.69 | 5180.3 | 4104.19 | 4885.56 | GO:0005 | GO:0030 | GO:0035864///response to potassium ion |
| 80 | 945363 | 'ybaT' | mRNA | -4.8611 | ######## | 47.59 | 46.88 | 44.9 | 552.2 | 479.23 | 435.64 | GO:0005 | GO:0022 | GO:0006865///amino acid transport+++GO:0009268///response to pH+++GO:0055085///transmembrane transport+++GO:1990169///stress response to copper ion |
| 81 | 948201 | 'uhpT' | mRNA | -4.8517 | ######## | 3.23 | 3.64 | 3.47 | 35.53 | 37.5 | 36.85 | GO:0005 | GO:0015 | GO:0006817///phosphate ion transport+++GO:0008643///carbohydrate transport+++GO:0015712///hexose phosphate transport+++GO:0015760///glucose-6-phosphate transport+++GO:0055085///transmembrane transport |
| 82 | 947830 | 'yhcO' | mRNA | -4.8096 | 4.06E-69 | 55.22 | 57.36 | 48.44 | 503.15 | 488.05 | 548.18 | | | |
| 83 | 948668 | 'frdD' | mRNA | -4.8066 | 7.81E-26 | 10.65 | 16.11 | 4.24 | 144.53 | 105.58 | 91.48 | GO:0005 | GO:0000 | GO:0006106///fumarate metabolic process+++GO:0006113///fermentation+++GO:0009061///anaerobic respiration |
| 84 | 947595 | 'paaB' | mRNA | -4.7909 | 0.01094 | 0 | 0 | 0 | 0.8 | 0.82 | 1.17 | | GO:0005 | GO:0010124///phenylacetate catabolic process |
| 85 | 945669 | 'cspD' | mRNA | -4.7677 | 1.94E-30 | 119.05 | 49.78 | 155.82 | 557.98 | 794.32 | 1112.69 | GO:0005 | GO:0001 | GO:0006355///regulation of transcription, DNA-templated+++GO:0008156///negative regulation of DNA replication+++GO:0009405///pathogenesis+++GO:0010468///regulation of gene expression+++GO:0042594///response to starvation+++GO:0060567///negative regulation of DNA-templated transcription, termination |
| 86 | 945491 | 'ybjP' | mRNA | -4.7642 | 1.85E-96 | 126.26 | 88.41 | 183.15 | 1375.29 | 1127.49 | 1091.2 | GO:0005886///plasma membrane+++GO:0016020///membrane | | |
| 87 | 945567 | 'rmf' | mRNA | -4.743 | 4.29E-21 | 5860.3 | 6761.84 | 3471.4 | 45017.3 | 45903.5 | 43353.6 | GO:0005 | GO:0003 | GO:0006417///regulation of translation+++GO:0032055///negative regulation of translation in response to stress |
| 88 | 948024 | 'hdeD' | mRNA | -4.733 | ######## | 312.02 | 327.62 | 354.15 | 3642.76 | 3243.45 | 2350.47 | GO:0005886///plas | | GO:0009268///response to pH |
| 89 | 945229 | 'uspG' | mRNA | -4.7197 | 2.89E-79 | 541.94 | 538.2 | 466.63 | 4647.39 | 4583.28 | 4818.38 | GO:0005 | GO:0004 | GO:0018117///protein adenylylation+++GO:0046777///protein autophosphorylation+++GO:0046940///nucleoside monophosphate phosphorylation+++GO:2000145///regulation of cell motility |
| 90 | 946174 | 'ldtE' | mRNA | -4.6927 | ######## | 195.38 | 180.22 | 184.65 | 1892.55 | 1702.79 | 1589.73 | GO:0042 | GO:0016 | GO:0006508///proteolysis+++GO:0008360///regulation of cell shape+++GO:0009252///peptidoglycan biosynthetic process+++GO:0018104///peptidoglycan-protein cross-linking+++GO:0071555///cell wall organization |
| 91 | 948008 | 'uspB' | mRNA | -4.689 | 1.97E-36 | 49.56 | 36.56 | 143.86 | 554.73 | 559.11 | 582.79 | GO:0005886///plas | | GO:0045471///response to ethanol |
| 92 | 948556 | 'qorA' | mRNA | -4.6628 | 1.70E-76 | 140.65 | 123.88 | 252.8 | 1500.24 | 1443.93 | 1552.12 | GO:0005 | GO:0003 | GO:0034599///cellular response to oxidative stress+++GO:0055114///oxidation-reduction process |
| 93 | 947523 | 'mcbA' | mRNA | -4.6414 | 7.60E-17 | 18.25 | 14.9 | 114.11 | 259.27 | 296.08 | 305.41 | GO:0031236///extr | | GO:0009242///colanic acid biosynthetic process |
| 94 | 944891 | 'rof' | mRNA | -4.6221 | 1.33E-28 | 190.8 | 178.01 | 105.15 | 1585.07 | 1320.87 | 1057.21 | | GO:0005 | GO:0031554///regulation of DNA-templated transcription, termination+++GO:0031564///transcription antitermination |
| 95 | 948680 | 'frdC' | mRNA | -4.6211 | 3.00E-69 | 11.1 | 11.83 | 12.69 | 124.12 | 96.46 | 85.39 | GO:0005 | GO:0000 | GO:0006113///fermentation+++GO:0009061///anaerobic respiration+++GO:0044780///bacterial-type flagellum assembly+++GO:0071973///bacterial-type flagellum-dependent cell motility |
| 96 | 1450233 | 'yaeP' | mRNA | -4.5918 | 1.15E-13 | 642.66 | 834.92 | 227.31 | 6057.26 | 4297.54 | 3630.13 | | | |
| 97 | 948023 | 'gadE' | mRNA | -4.5669 | 3.88E-04 | 10.07 | 8.88 | 140.87 | 133.09 | 487.01 | 484.28 | | GO:0003 | GO:0006355///regulation of transcription, DNA-templated |
| 98 | 946397 | 'tap' | mRNA | -4.5587 | 1.22E-74 | 1.8 | 1.3 | 1.71 | 14.79 | 12.57 | 13.66 | GO:0005 | GO:0004 | GO:0006935///chemotaxis+++GO:0007165///signal transduction |
| 99 | 947077 | 'clpB' | mRNA | -4.5244 | 1.49E-60 | 275.86 | 284.83 | 563.97 | 3304.28 | 3023.17 | 2763.56 | GO:0005 | GO:0000 | GO:0006986///response to unfolded protein+++GO:0009408///response to heat+++GO:0019538///protein metabolic process+++GO:0042026///protein refolding |
| 100 | 947956 | 'yhhA' | mRNA | -4.5046 | 5.21E-11 | 57.75 | 33.42 | 327.94 | 678.6 | 869.97 | 1219.4 | | GO:0005515///protein binding | |
| 101 | 948666 | 'frdB' | mRNA | -4.4953 | 1.15E-72 | 8.24 | 7.63 | 8.65 | 82.1 | 55.45 | 60.47 | GO:0005 | GO:0000 | GO:0006099///tricarboxylic acid cycle+++GO:0006113///fermentation+++GO:0009061///anaerobic respiration+++GO:0022900///electron transport chain+++GO:0044780///bacterial-type flagellum assembly+++GO:0055114///oxidation-reduction process+++GO:0071973///bacterial-type flagellum-dependent cell motility |
| 102 | 948553 | 'yjbJ' | mRNA | -4.4442 | 7.05E-66 | 1668.59 | 1260.5 | 3331.45 | 10051 | 11207.7 | 15154.6 | GO:0005829///cytosol | | |
| 103 | 947599 | 'uxaC' | mRNA | -4.3824 | 1.42E-35 | 5.39 | 4.19 | 3.41 | 25.04 | 32.82 | 42.33 | | GO:0008 | GO:0006064///glucuronate catabolic process+++GO:0019698///D-galacturonate catabolic process+++GO:0042840///D-glucuronate catabolic process |
| 104 | 947919 | 'feoB' | mRNA | -4.3511 | ######## | 12.26 | 14.12 | 12.28 | 100 | 103.24 | 85.93 | GO:0005 | GO:0000 | GO:0006811///ion transport+++GO:0006826///iron ion transport+++GO:0006974///cellular response to DNA damage stimulus+++GO:0034755///iron ion transmembrane transport+++GO:0055072///iron ion homeostasis+++GO:0072511///divalent inorganic cation transport+++GO:0098711///iron ion import across plasma membrane |
| 105 | 948713 | 'yjfY' | mRNA | -4.322 | 1.03E-24 | 7.45 | 3.93 | 22.08 | 50.77 | 51.18 | 50.63 | GO:0042597///periplasmic space | | |
| 106 | 945762 | 'ychH' | mRNA | -4.3217 | 5.71E-54 | 262.55 | 182.23 | 224.81 | 1665.88 | 1496.72 | 1178.99 | | | GO:0044011///single-species biofilm formation on inanimate substrate+++GO:0070301///cellular response to hydrogen peroxide+++GO:0071276///cellular response to cadmium ion |
| 107 | 946106 | 'copA' | mRNA | -4.3156 | 1.72E-57 | 99.25 | 113.21 | 76.16 | 742.21 | 670.95 | 691.2 | GO:0005 | GO:0000 | GO:0006811///ion transport+++GO:0006812///cation transport+++GO:0006825///copper ion transport+++GO:0010273///detoxification of copper ion+++GO:0030001///metal ion transport+++GO:0034220///ion transmembrane transport+++GO:0060003///copper ion export+++GO:0071280///cellular response to copper ion+++GO:0071292///cellular response to silver ion+++GO:0098655///cation transmembrane transport+++GO:1902601///silver ion transmembrane transport |
| 108 | 945479 | 'grxA' | mRNA | -4.3013 | 1.30E-17 | 37.23 | 24.62 | 118.26 | 169.64 | 302.95 | 433.32 | GO:0005 | GO:0009 | GO:0009263///deoxyribonucleotide biosynthetic process+++GO:0022900///electron transport chain+++GO:0045454///cell redox homeostasis+++GO:0055114///oxidation-reduction process |
| 109 | 948071 | 'yiaG' | mRNA | -4.2707 | 1.45E-55 | 304.46 | 190.27 | 449.84 | 1505.55 | 1841 | 2134.76 | | GO:0003677///DNA binding+++GO:0043565///sequence-specific DNA binding | |
| 110 | 946719 | 'atoA' | mRNA | -4.2626 | 5.72E-09 | 0.29 | 0.28 | 0 | 2.07 | 1.55 | 1.84 | | GO:0005 | GO:0006629///lipid metabolic process+++GO:0006631///fatty acid metabolic process+++GO:0046459///short-chain fatty acid metabolic process |
| 111 | 944985 | 'yahO' | mRNA | -4.2495 | 1.42E-30 | 197.58 | 139.84 | 510.71 | 1155.22 | 1475.6 | 1849.6 | GO:0042597///peri | | GO:0009314///response to radiation |
| 112 | 946399 | 'tar' | mRNA | -4.2412 | 4.17E-53 | 1.5 | 1.69 | 1.79 | 14.07 | 11.42 | 8.79 | GO:0005 | GO:0004 | GO:0006935///chemotaxis+++GO:0007165///signal transduction+++GO:0007172///signal complex assembly+++GO:0009593///detection of chemical stimulus+++GO:0050920///regulation of chemotaxis+++GO:0051260///protein homooligomerization+++GO:0071230///cellular response to amino acid stimulus+++GO:1901875///positive regulation of post-translational protein modification |
| 113 | 946469 | 'yodD' | mRNA | -4.234 | 3.52E-18 | 239.46 | 310.52 | 127.97 | 1607.77 | 1414.51 | 1305.93 | | | GO:0044011///single-species biofilm formation on inanimate substrate+++GO:0070301///cellular response to hydrogen peroxide+++GO:0071468///cellular response to acidic pH |
| 114 | 948560 | 'yjbR' | mRNA | -4.2309 | 3.62E-57 | 182.59 | 175.76 | 152.51 | 1213.4 | 1026.9 | 1039.01 | | | |
| 115 | 947533 | 'glgS' | mRNA | -4.1911 | 6.56E-11 | 158.39 | 77.92 | 926.09 | 993.17 | 1355.58 | 2182.79 | | | GO:0010676///positive regulation of cellular carbohydrate metabolic process+++GO:1900191///negative regulation of single-species biofilm formation+++GO:1900232///negative regulation of single-species biofilm formation on inanimate substrate+++GO:1902201///negative regulation of bacterial-type flagellum-dependent cell motility |
| 116 | 5061528 | 'yjbT' | mRNA | -4.1788 | 3.25E-28 | 6.31 | 4.42 | 5.35 | 33.69 | 31.6 | 36.16 | | | |
| 117 | 945340 | 'ariR' | mRNA | -4.1774 | 1.04E-04 | 0.7 | 0.65 | 0 | 3.19 | 2.62 | 6.29 | | GO:0003 | GO:0042542///response to hydrogen peroxide+++GO:0042710///biofilm formation+++GO:0071468///cellular response to acidic pH |
| 118 | 944878 | 'sufD' | mRNA | -4.1649 | 1.18E-73 | 184.78 | 171.73 | 321.57 | 1482.82 | 1377.52 | 1370.23 | GO:1990 | GO:0005 | GO:0006979///response to oxidative stress+++GO:0016226///iron-sulfur cluster assembly |
| 119 | 948895 | 'osmY' | mRNA | -4.1648 | 1.14E-49 | 611.95 | 674.96 | 520.26 | 4669.79 | 3260.59 | 3585.08 | GO:0030 | GO:0005 | GO:0006970///response to osmotic stress+++GO:0006972///hyperosmotic response+++GO:0061077///chaperone-mediated protein folding |
| 120 | 946128 | 'sufC' | mRNA | -4.1524 | ######## | 185.71 | 209.35 | 214.7 | 1425.68 | 1348.62 | 1065.83 | GO:0005 | GO:0000 | GO:0009314///response to radiation+++GO:0016226///iron-sulfur cluster assembly |
| 121 | 946314 | 'yoaG' | mRNA | -4.145 | 7.43E-11 | 54.09 | 88.46 | 0 | 343.18 | 298.08 | 350.72 | | | |
| 122 | 947610 | 'yqjK' | mRNA | -4.1355 | 1.38E-16 | 1003.89 | 933.76 | 377.29 | 5017.25 | 4675.92 | 4498.59 | GO:0005829///cytosol | | |
| 123 | 946187 | 'glsA' | mRNA | -4.1291 | 9.18E-23 | 45.81 | 42.7 | 140.86 | 512.7 | 440.86 | 388.28 | | GO:0004 | GO:0006537///glutamate biosynthetic process+++GO:0006541///glutamine metabolic process+++GO:0006543///glutamine catabolic process+++GO:0010447///response to acidic pH+++GO:0045926///negative regulation of growth+++GO:0051289///protein homotetramerization |
| 124 | 946724 | 'acnA' | mRNA | -4.1282 | 1.86E-65 | 73.81 | 73.28 | 131.59 | 653.49 | 535.32 | 530.13 | GO:0005 | GO:0003 | GO:0006097///glyoxylate cycle+++GO:0006099///tricarboxylic acid cycle+++GO:0006101///citrate metabolic process+++GO:0006979///response to oxidative stress+++GO:0009061///anaerobic respiration |
| 125 | 945070 | 'ybaY' | mRNA | -4.1184 | 1.16E-26 | 364.58 | 393.54 | 196.75 | 2252.59 | 1908.92 | 1836.92 | GO:0005 | GO:0005515///protein binding | |
| 126 | 948667 | 'frdA' | mRNA | -4.1056 | 9.43E-74 | 11.79 | 12.16 | 19.32 | 105.29 | 80.01 | 75.41 | GO:0005 | GO:0000 | GO:0006113///fermentation+++GO:0006974///cellular response to DNA damage stimulus+++GO:0009061///anaerobic respiration+++GO:0022900///electron transport chain+++GO:0044780///bacterial-type flagellum assembly+++GO:0055114///oxidation-reduction process+++GO:0071973///bacterial-type flagellum-dependent cell motility |
| 127 | 946185 | 'sufS' | mRNA | -4.0849 | ######## | 163.79 | 165.47 | 251.21 | 1194.61 | 1183.2 | 1076.74 | GO:0005 | GO:0003 | GO:0001887///selenium compound metabolic process+++GO:0006534///cysteine metabolic process+++GO:0006790///sulfur compound metabolic process+++GO:0016226///iron-sulfur cluster assembly+++GO:0031162///sulfur incorporation into metallo-sulfur cluster |
| 128 | 2847737 | 'ecnB' | mRNA | -4.0588 | 1.04E-34 | 487.73 | 398.35 | 822.26 | 1718.8 | 2228.7 | 3244.7 | GO:0005886///plas | | GO:0009636///response to toxic substance |
| 129 | 947839 | 'bfr' | mRNA | -4.0391 | 3.82E-29 | 411.47 | 275.78 | 851.13 | 2327.95 | 2509.62 | 3239.53 | GO:0005 | GO:0004 | GO:0006826///iron ion transport+++GO:0006879///cellular iron ion homeostasis+++GO:0006880///intracellular sequestering of iron ion+++GO:0055114///oxidation-reduction process |
| 130 | 948902 | 'deoC' | mRNA | -4.0376 | 2.85E-53 | 94.07 | 98.18 | 72.86 | 595.03 | 504.5 | 474.11 | GO:0005 | GO:0003 | GO:0006974///cellular response to DNA damage stimulus+++GO:0009264///deoxyribonucleotide catabolic process+++GO:0015949///nucleobase-containing small molecule interconversion+++GO:0016052///carbohydrate catabolic process+++GO:0046386///deoxyribose phosphate catabolic process |
| 131 | 946530 | 'ivy' | mRNA | -4.0325 | ######## | 625.77 | 583.42 | 792.74 | 3697.35 | 3651.71 | 3712.27 | GO:0030 | GO:0060 | GO:0043086///negative regulation of catalytic activity+++GO:0050790///regulation of catalytic activity+++GO:0061077///chaperone-mediated protein folding |
| 132 | 947918 | 'feoC' | mRNA | -4.0016 | 2.10E-13 | 5.33 | 10.78 | 0 | 45.07 | 35.69 | 38.3 | | GO:0003677///DNA binding+++GO:0005506///iron ion binding+++GO:0046872///metal ion binding+++GO:0051536///iron-sulfur cluster binding | |
| 133 | 946317 | 'yeaR' | mRNA | -3.996 | 1.27E-66 | 191.09 | 246.14 | 245.73 | 1359.62 | 1122.6 | 1162.2 | | | |
| 134 | 945512 | 'ycaC' | mRNA | -3.966 | 6.33E-68 | 87.14 | 74.76 | 143.75 | 501.3 | 522.5 | 576.51 | | GO:0003824///catalytic activity+++GO:0016787///hydrolase activity+++GO:0016829///lyase activity+++GO:0042802///identical protein binding | |
| 135 | 947138 | 'yqaE' | mRNA | -3.9613 | 9.69E-33 | 157.46 | 96.78 | 312.15 | 551.31 | 663.59 | 871.03 | GO:0005886///plas | | GO:0046677///response to antibiotic |
| 136 | 946398 | 'pgl' | mRNA | -3.937 | ######## | 144.6 | 147.75 | 220.33 | 974.95 | 984.18 | 786.64 | GO:0005 | GO:0016 | GO:0005975///carbohydrate metabolic process+++GO:0006006///glucose metabolic process+++GO:0006098///pentose-phosphate shunt |
| 137 | 947617 | 'yhaH' | mRNA | -3.9275 | ######## | 87.54 | 93.25 | 118.48 | 545.27 | 529.78 | 433.61 | GO:0005886///plasma membrane+++GO:0016020///membrane+++GO:0016021///integral component of membrane | | |
| 138 | 945396 | 'moaB' | mRNA | -3.9257 | ######## | 80.67 | 86.47 | 102.83 | 484.99 | 443.57 | 478.9 | GO:0005 | GO:0000 | GO:0006777///Mo-molybdopterin cofactor biosynthetic process+++GO:0034214///protein hexamerization |
| 139 | 1450246 | 'cydX' | mRNA | -3.9088 | 4.43E-21 | 0 | 0 | 0 | 0 | 0 | 0 | GO:0005 | GO:0016 | GO:0006119///oxidative phosphorylation+++GO:0055114///oxidation-reduction process |
| 140 | 947609 | 'ycgR' | mRNA | -3.8812 | 8.12E-21 | 1.16 | 0.76 | 2.47 | 6.43 | 6.95 | 5.14 | GO:0009 | GO:0000 | GO:0051270///regulation of cellular component movement+++GO:0071945///regulation of bacterial-type flagellum-dependent cell motility by regulation of motor speed+++GO:0071973///bacterial-type flagellum-dependent cell motility |
| 141 | 948031 | 'mdtE' | mRNA | -3.8731 | 6.30E-61 | 25.97 | 24.83 | 20.75 | 143.33 | 135.41 | 105.59 | GO:0005 | GO:0005 | GO:0009636///response to toxic substance+++GO:0015721///bile acid and bile salt transport+++GO:0046677///response to antibiotic+++GO:0055085///transmembrane transport+++GO:1990961///drug transmembrane export |
| 142 | 945578 | 'hspQ' | mRNA | -3.8715 | ######## | 485.69 | 509.97 | 666.96 | 2941.35 | 2574.38 | 2316.48 | GO:0005 | GO:0003 | GO:0009408///response to heat |
| 143 | 948019 | 'yhiD' | mRNA | -3.8454 | 3.25E-90 | 45.11 | 38.06 | 68.67 | 268.61 | 245.51 | 220.89 | GO:0005886///plasma membrane+++GO:0016020///membrane+++GO:0016021///integral component of membrane | | |
| 144 | 947365 | 'yphA' | mRNA | -3.8453 | 3.29E-25 | 115.28 | 115.71 | 64.74 | 558.56 | 527.25 | 433.46 | GO:0005886///plasma membrane+++GO:0016020///membrane+++GO:0016021///integral component of membrane | | |
| 145 | 946102 | 'ydfD' | mRNA | -3.8225 | 0.00818 | 1.97 | 0 | 0 | 4.27 | 6.18 | 3.86 | | | GO:0044659///cytolysis by virus of host cell |
| 146 | 945753 | 'sufB' | mRNA | -3.8196 | 6.13E-34 | 141.13 | 135.68 | 309.12 | 1033.73 | 1047.11 | 790.03 | GO:1990 | GO:0005 | GO:0016226///iron-sulfur cluster assembly |
| 147 | 946401 | 'cheA' | mRNA | -3.8158 | 3.52E-79 | 3.63 | 2.92 | 3.94 | 18.82 | 16.19 | 17.71 | GO:0005 | GO:0000 | GO:0000160///phosphorelay signal transduction system+++GO:0006935///chemotaxis+++GO:0007165///signal transduction+++GO:0009454///aerotaxis+++GO:0016310///phosphorylation+++GO:0018106///peptidyl-histidine phosphorylation+++GO:0023014///signal transduction by protein phosphorylation+++GO:0031400///negative regulation of protein modification process+++GO:0043052///thermotaxis+++GO:0046777///protein autophosphorylation+++GO:0050920///regulation of chemotaxis+++GO:0051649///establishment of localization in cell+++GO:1901875///positive regulation of post-translational protein modification+++GO:1902021///regulation of bacterial-type flagellum-dependent cell motility |
| 148 | 946173 | 'sufE' | mRNA | -3.8149 | ######## | 158.56 | 153.8 | 253.77 | 862.07 | 910.5 | 822.33 | GO:0005 | GO:0005 | GO:0006790///sulfur compound metabolic process+++GO:0006979///response to oxidative stress+++GO:0016226///iron-sulfur cluster assembly+++GO:0031162///sulfur incorporation into metallo-sulfur cluster+++GO:0043085///positive regulation of catalytic activity+++GO:0051347///positive regulation of transferase activity |
| 149 | 7751624 | 'sgrT' | mRNA | -3.8125 | 0.00462 | 0 | 24.46 | 0 | 35.62 | 80.64 | 51.08 | | GO:0004 | GO:0008643///carbohydrate transport+++GO:0009401///phosphoenolpyruvate-dependent sugar phosphotransferase system+++GO:0043086///negative regulation of catalytic activity+++GO:0046325///negative regulation of glucose import |
| 150 | 947495 | 'dkgA' | mRNA | -3.7726 | 6.84E-13 | 152.58 | 100.87 | 523.17 | 1034.61 | 1112.37 | 1311.01 | GO:0005 | GO:0004 | GO:0019853///L-ascorbic acid biosynthetic process+++GO:0051596///methylglyoxal catabolic process+++GO:0055114///oxidation-reduction process |

| | | | | | | | | | | | | |
|---|---|---|---|---|---|---|---|---|---|---|---|---|
| 948362 | 'hemN' | mRNA | -0.3008 | 0.03377 | 128.29 | 112.26 | 143.16 | 56.92 | 57.18 | 54.03 | GO:0005: GO:0003 | GO:0006779///porphyrin-containing compound biosynthetic process+++GO:0006782///protoporphyrinogen IX biosynthetic process+++GO:0019353///protoporphyrinogen IX biosynthetic process from glutamate+++GO:0055114///oxidation-reduction process |
