## Supplementary Data 4 for "Enhanced extracellular matrix production provides protection to cell wall-deficient *Escherichia coli*"

Supplementary Data 4. Overview of the expression levels of extracellular matrix genes.

**Colanic Acid**

| | **Wild Type** | | | **Spheroplast** | | | **L-form** | | | **Fold Change** | |
|---|---|---|---|---|---|---|---|---|---|---|---|
| | Mean | SD | N | Mean | SD | N | Mean | SD | N | Wild Type vs L-form | Spheroplast vs L-form |
| cpsB | 0.593 | 0.058 | 3 | 33.080 | 3.057 | 3 | 438.393 | 56.446 | 3 | 738.87 | 13.25 |
| fcl | 0.737 | 0.424 | 3 | 66.390 | 5.249 | 3 | 1833.407 | 580.656 | 3 | 2488.79 | 27.62 |
| gmd | 1.067 | 0.528 | 3 | 156.880 | 16.157 | 3 | 3745.997 | 567.253 | 3 | 3511.87 | 23.88 |
| gmm | 0.840 | 0.213 | 3 | 54.337 | 6.905 | 3 | 1207.820 | 278.731 | 3 | 1437.88 | 22.23 |
| rfbX | 13.930 | 6.424 | 3 | 28.193 | 22.024 | 3 | 96.230 | 115.072 | 3 | 6.91 | 3.41 |
| wcaA | 1.557 | 0.437 | 3 | 95.177 | 12.587 | 3 | 1101.510 | 520.059 | 3 | 707.61 | 11.57 |
| wcaB | 0.580 | 0.459 | 3 | 51.480 | 3.870 | 3 | 536.137 | 192.138 | 3 | 924.37 | 10.41 |
| wcaC | 0.777 | 0.006 | 3 | 82.727 | 10.086 | 3 | 707.527 | 5.779 | 3 | 910.98 | 8.55 |
| wcaD | 0.513 | 0.095 | 3 | 39.993 | 11.578 | 3 | 653.043 | 41.596 | 3 | 1272.16 | 16.33 |
| wcaE | 0.597 | 0.140 | 3 | 48.827 | 19.466 | 3 | 1062.730 | 449.830 | 3 | 1781.11 | 21.77 |
| wcaF | 0.183 | 0.095 | 3 | 36.753 | 9.001 | 3 | 561.087 | 95.811 | 3 | 3060.47 | 15.27 |
| wcaI | 1.160 | 0.132 | 3 | 49.670 | 6.493 | 3 | 863.150 | 35.062 | 3 | 744.09 | 17.38 |
| wcaJ | 0.887 | 0.060 | 3 | 38.773 | 1.500 | 3 | 460.470 | 110.941 | 3 | 519.33 | 11.88 |
| wcaK | 1.037 | 0.015 | 3 | 51.013 | 5.847 | 3 | 631.843 | 59.047 | 3 | 609.50 | 12.39 |
| wcaL | 1.460 | 0.213 | 3 | 38.810 | 3.118 | 3 | 429.467 | 166.845 | 3 | 294.16 | 11.07 |
| wcaM | 0.910 | 0.219 | 3 | 17.500 | 4.540 | 3 | 466.547 | 147.135 | 3 | 512.69 | 26.66 |
| wza | 0.430 | 0.202 | 3 | 104.227 | 20.364 | 3 | 1551.703 | 622.184 | 3 | 3608.61 | 14.89 |
| wzb | 0.457 | 0.159 | 3 | 121.893 | 17.868 | 3 | 1431.087 | 51.322 | 3 | 3133.77 | 11.74 |
| wzc | 0.550 | 0.092 | 3 | 76.400 | 10.141 | 3 | 804.960 | 8.129 | 3 | 1463.56 | 10.54 |

**Curli Protein**

| | **Wild Type** | | | **Spheroplast** | | | **L-form** | | | **Fold Change** | |
|---|---|---|---|---|---|---|---|---|---|---|---|
| | Mean | SD | N | Mean | SD | N | Mean | SD | N | Wild Type vs L-form | Spheroplast vs L-form |
| csgA | 2.023 | 0.259 | 3 | 3.397 | 0.376 | 3 | 5.627 | 1.093 | 3 | 2.78 | 1.66 |
| csgB | 0.240 | 0.208 | 3 | 0.447 | 0.156 | 3 | 1.217 | 0.480 | 3 | 5.07 | 2.72 |
| csgC | 0.613 | 0.191 | 3 | 2.950 | 0.495 | 3 | 3.590 | 2.998 | 3 | 5.85 | 1.22 |
| csgE | 0.200 | 0.229 | 3 | 0.407 | 0.383 | 3 | 7.280 | 2.156 | 3 | 36.40 | 17.90 |
| csgF | 0.313 | 0.199 | 3 | 0.373 | 0.198 | 3 | 2.447 | 1.572 | 3 | 7.81 | 6.55 |
| csgG | 2.637 | 0.526 | 3 | 13.043 | 2.494 | 3 | 6.717 | 2.538 | 3 | 2.55 | 0.51 |
| csgD | 0.3333 | 0.1097 | 3 | 1.337 | 0.5173 | 3 | 10.38 | 6.793 | 3 | 31.14 | 7.76 |

**Cellulose**

| | **Wild Type** | | | **Spheroplast** | | | **L-form** | | | **Fold Change** | |
|---|---|---|---|---|---|---|---|---|---|---|---|
| | Mean | SD | N | Mean | SD | N | Mean | SD | N | L-form vs Wild Type | L-form vs Spheroplast |
| bcsA | 1.397 | 0.156 | 3 | 5.340 | 0.805 | 3 | 13.390 | 3.001 | 3 | 9.59 | 2.51 |
| bcsB | 3.027 | 0.060 | 3 | 7.123 | 1.499 | 3 | 15.180 | 4.097 | 3 | 5.02 | 2.13 |
| bcsC | 20.143 | 2.886 | 3 | 27.163 | 4.604 | 3 | 19.697 | 9.764 | 3 | 0.98 | 0.73 |
| bcsE | 32.003 | 2.150 | 3 | 66.467 | 3.067 | 3 | 129.917 | 80.359 | 3 | 4.06 | 1.95 |
| bcsF | 34.557 | 10.720 | 3 | 80.147 | 16.847 | 3 | 128.027 | 64.683 | 3 | 3.70 | 1.60 |
| bcsG | 8.433 | 0.564 | 3 | 19.127 | 3.652 | 3 | 68.690 | 30.231 | 3 | 8.15 | 3.59 |
| bcsZ | 2.920 | 0.468 | 3 | 5.177 | 0.971 | 3 | 10.907 | 1.684 | 3 | 3.74 | 2.11 |
