## Supplementary Figures and Supplementary Tables for "Enhanced extracellular matrix production provides protection to cell wall-deficient *Escherichia coli*"

*Supplementary Information*

| **Supplementary Figures** | | **Page** |
| --- | --- | --- |
| Supplementary Fig. 1 | Visualization of DNA distribution in L-form^0^. | 3 |
| Supplementary Fig. 2 | Sanger sequence confirmation of L-form^0^ SNPs. | 4 |
| Supplementary Fig. 3 | ROS detection in Wild Type, Revertant^0^, spheroplast and L-form^0^. | 5 |
| Supplementary Fig. 4 | Knockouts of Rcs are unable to survive penicillin treatment. | 6 |
| Supplementary Fig. 5 | Variation in expression of extracellular matrix proteins in L-form0, spheroplast and wild type bacteria. | 8 |
| Supplementary Fig. 6 | Expression levels of csgD and nhaR, regulators of cellulose, curli and PGA. | 9 |
| Supplementary Fig. 7 | Cellulose biosynthesis restored after bcsQ repair. | 10 |
| Supplementary Fig. 8 | Analysis of rcsA in L-form^LTE^. | 11 |
| Supplementary Fig. 9 | ROS detection in L-form^LTE^. | 12 |
| Supplementary Fig. 10 | Transmission light microscopy images of Wild Type, AmpR, Revertant^0^ and Revertant^LTE^ strains after stress test. | 13 |
| Supplementary Fig. 11 | Proposed model of RcsA-activated cellulose and curli protein synthesis. |  |
| **Supplementary Tables** |  | 14 |
| Supplementary Table 1 | Cell Size descriptive statistics. | 15 |
| Supplementary Table 2 | ROS detection of L-form0 using flow cytometry. | 16 |
| Supplementary Table 3 | RNA sequencing data of *ndh, rcsA, yjbH, ykgH, csgD,* and *nhaR*. | 17 |
| Supplementary Table 4 | List of SNPs found in the L-form^LTE^ strain. | 18 |
| Supplementary Table 5 | Crystal Violet biofilm biomass assay. | 19 |
| Supplementary Table 6 | ROS detection of L-form^LTE^ using flow cytometry. | 20 |
| Supplementary Table 7 | Bacterial strains used in the paper. | 21 |
| Supplementary Table 8 | Primers used in this study. | 22 |
| References |  | 23 |

### Supplementary Figures

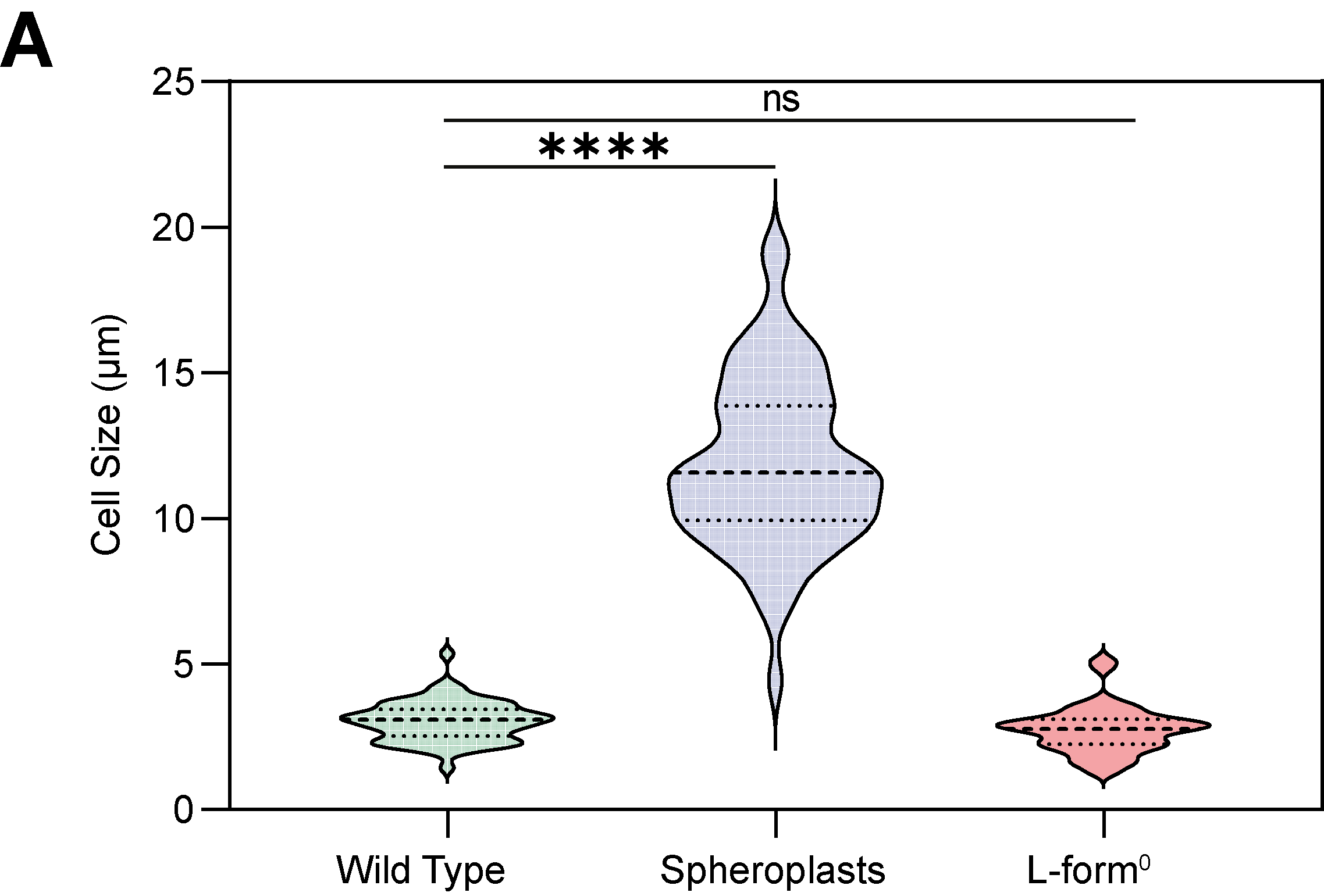

Supplementary Fig. 1 | **Average cell size of wild type, spheroplasts and L-form^0^ cells** (Supplementary Table 1). Data were analyzed using Kruskal–Wallis with Dunn's multiple comparison test.

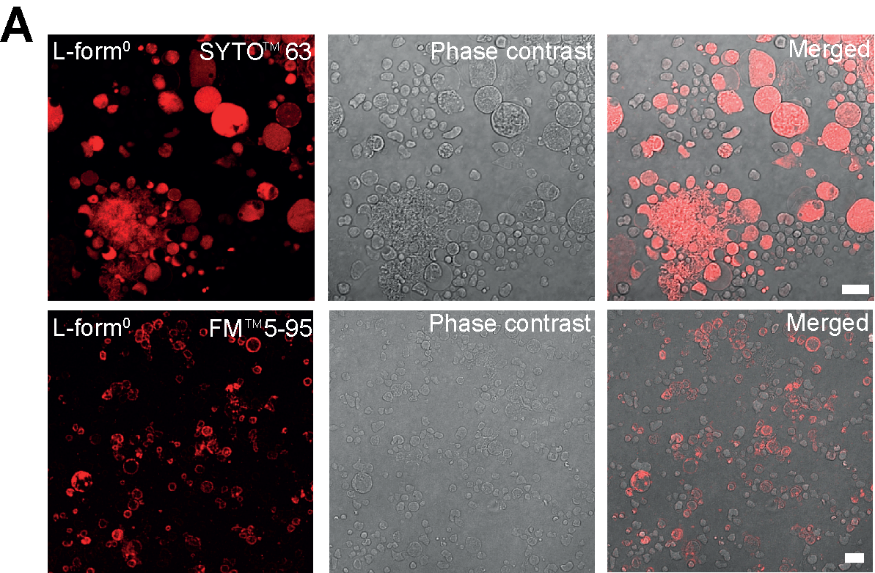

Supplementary Fig. 2 | Visualization of DNA distribution in L-form^0^ (A) Visualization of DNA (SYTO^TM^63) in L-form^0^ cells, cells imaged using the Airyscan microscope. Scalebar is 10µm.

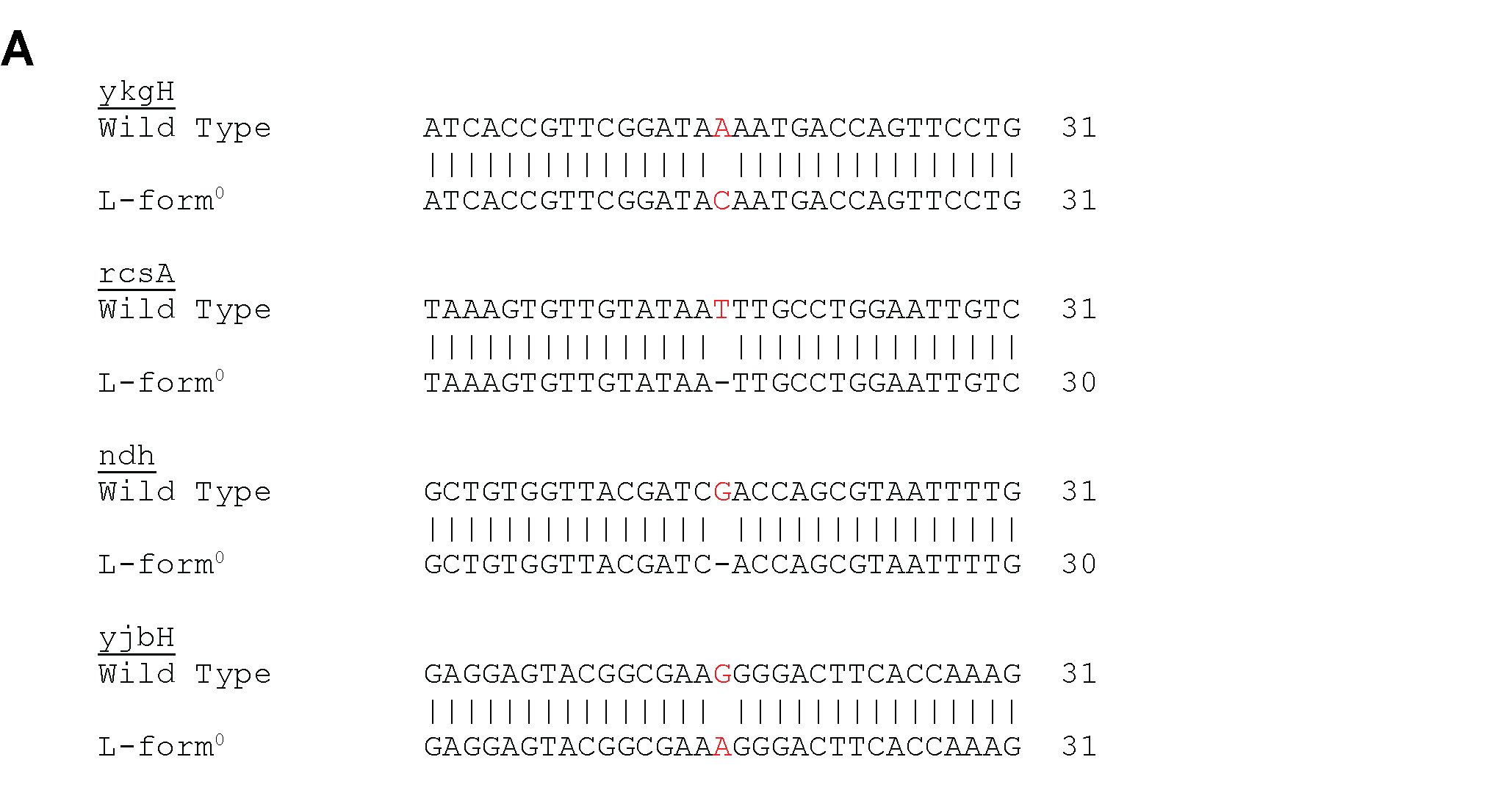

Supplementary Fig. 3 | Sanger sequence confirmation of L-form^0^ SNPs. (A) The regions of the SNPs were amplified in the wild type and L-form^0^ strains using PCR and Sanger sequenced. The mismatches are indicated in red. Primers can be found in Supplementary Table 8.

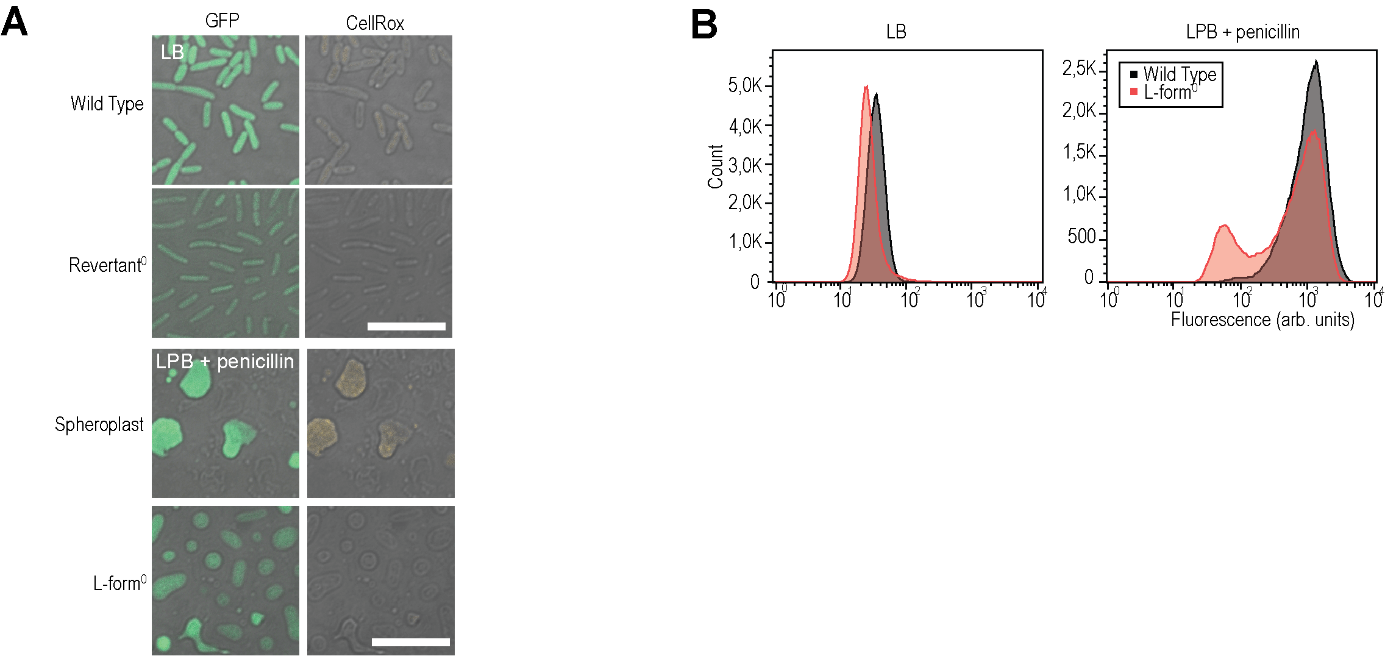

Supplementary Fig. 4 | ROS detection in wild type, revertant^0^, spheroplast and L-form^0^. (A) Fluorescence microscopy images depict exponentially growing cells stained with CellRox Orange to detect ROS in the L-form^0^ strain in wall-deficient and revertant cells. The GFP-labeled Wild Type (parental strain serW-GFP) spheroplasts and rod-shaped cells were used as a control. Scalebar is 10µm. (B) Quantitative analysis of ROS of the L-form^0^ (red) and wild type (black) in LB and LPB medium supplemented with penicillin using flow cytometry (n=100,000). Fluorescence is shown in arbitrary units. A summary of the data can be found in Supplementary Table 2.

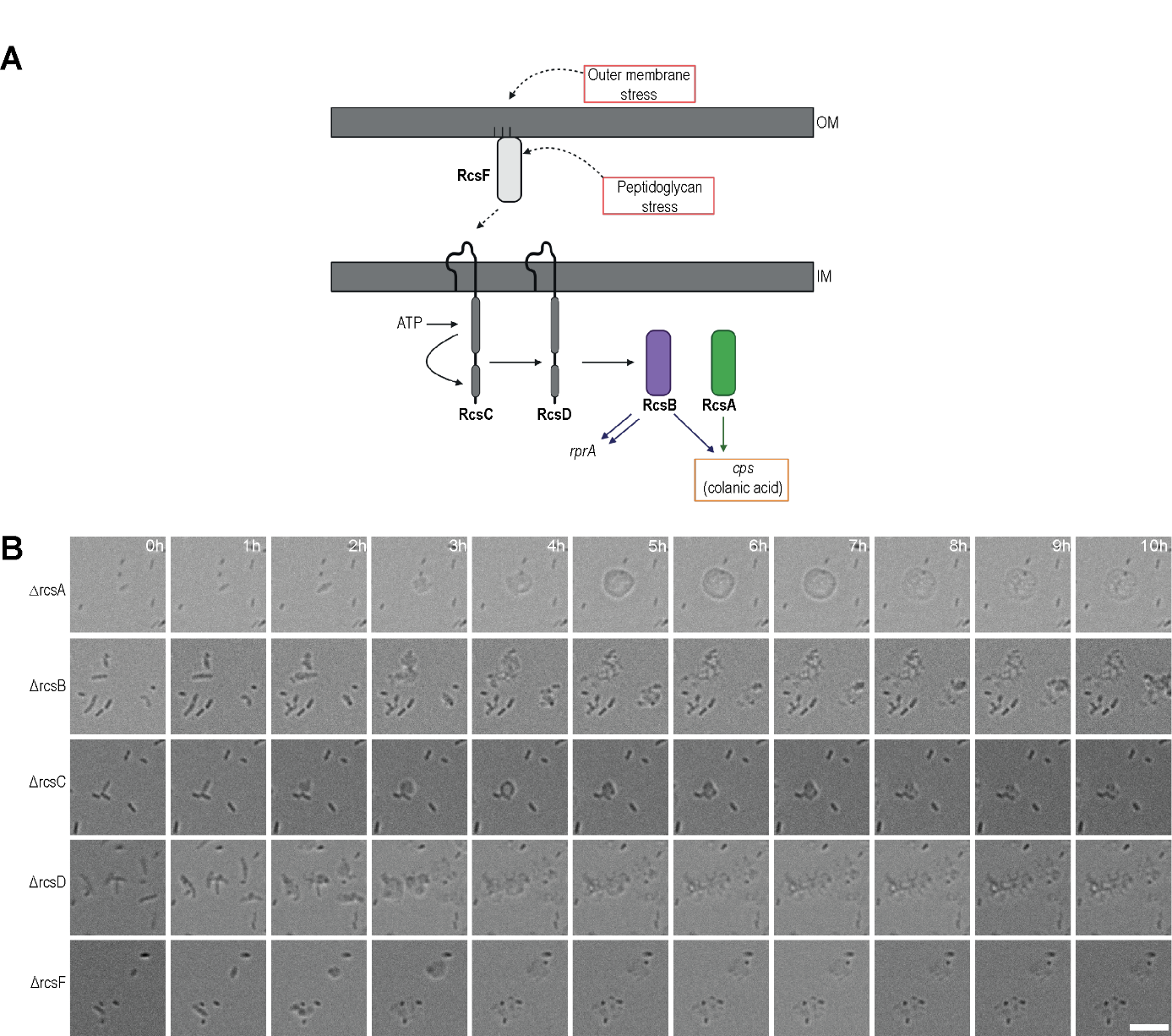

Supplementary Fig. 5 | Knockouts of Rcs are unable to survive penicillin treatment. (A) Schematic overview of the Rcs phosphorelay system within cellular envelope of *E. coli.* Outer membrane lipoprotein, RcsF, responds to stresses in the outer membrane and peptidoglycan, initiating the activation of the Rcs phosphorelay system. The histidine kinase, RcsC, autophosphorelates and subsequently transfers a phosphate group to the phosphotransferase, RcsD. RcsD then transfers the phosphate group to the transcription regulator, RcsB. RcsB has the capability to form either a homodimer (RcsB-RcsB), or a heterodimer (RcsB-RcsA) in the presence of the transcription regulator RcsA (Meng, Young and Chen 2021). Illustration was created using Biorender. (B) Time-lapse imaging of *E. coli* lacking RcsA, RcsB, RcsC, RcsD, and RcsD grown in LPB medium with agar pad supplemented with 0.4 mg ml^-1^ penicillin using the Lionheart microscope. Scalebar is 10μm.

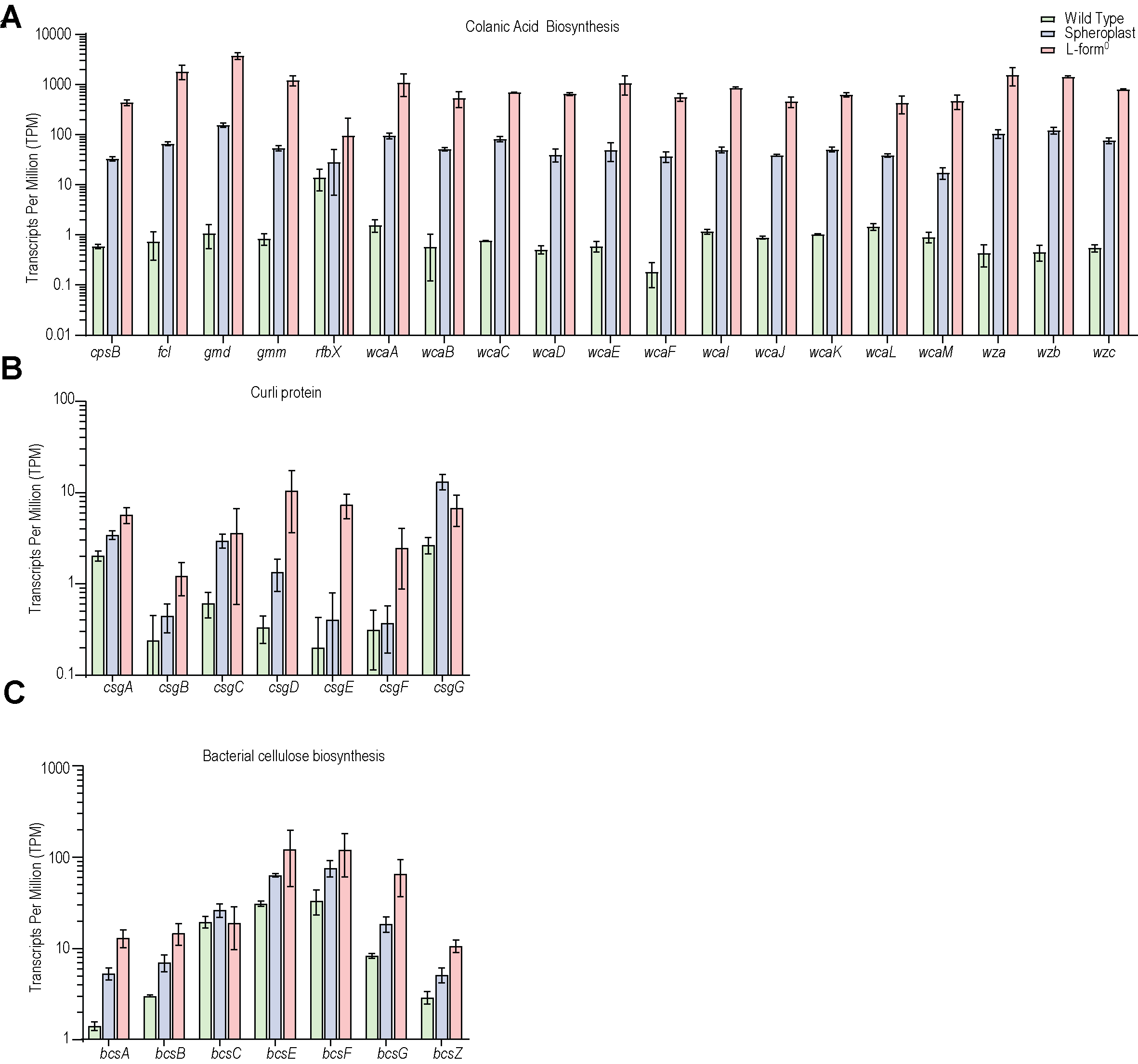

Supplementary Fig. 6 | Variation in expression of extracellular matrix proteins in L-form^0^, spheroplast and wild type bacteria. (A-C) RNA expression levels of genes involved in the biosynthesis of colanic acid (A), curli protein (B), and bacterial cellulose (C) in wild type (green), spheroplast (blue) and L-form^0^ (red). More information can be found in Supplementary Data 4.

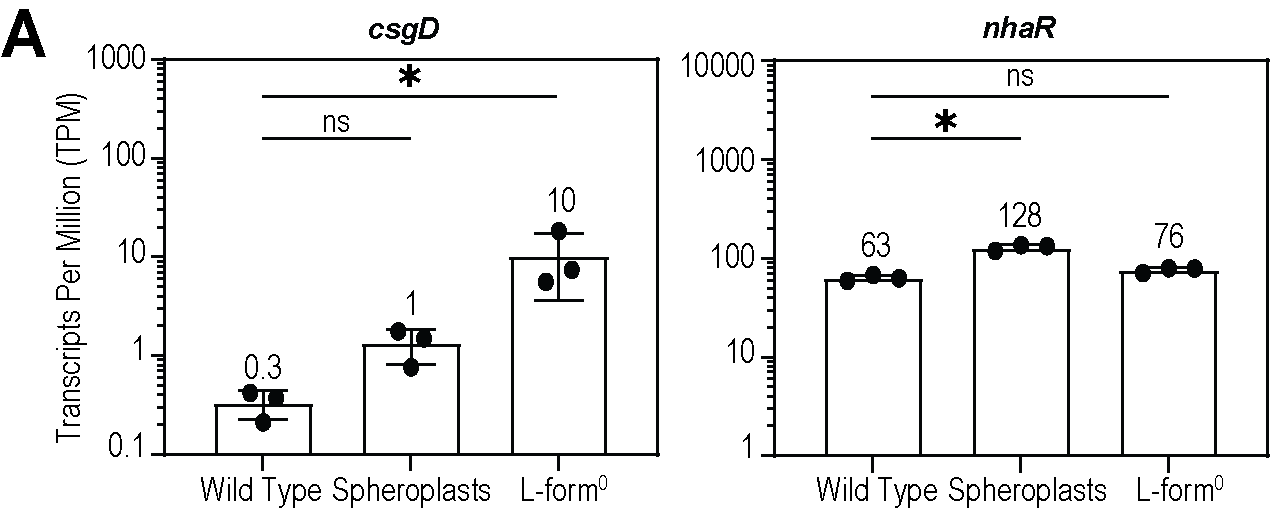

Supplementary Fig. 7 | Expression levels of *csgD* and *nhaR*, regulators of cellulose, curli and PGA. (A) Comparison of RNA levels of csgD and nhaR in wild type, spheroplasts and L-form^0^ cells. The P values were determined using Kruskal-Wallis following Dunnett’s multiple comparisons test.

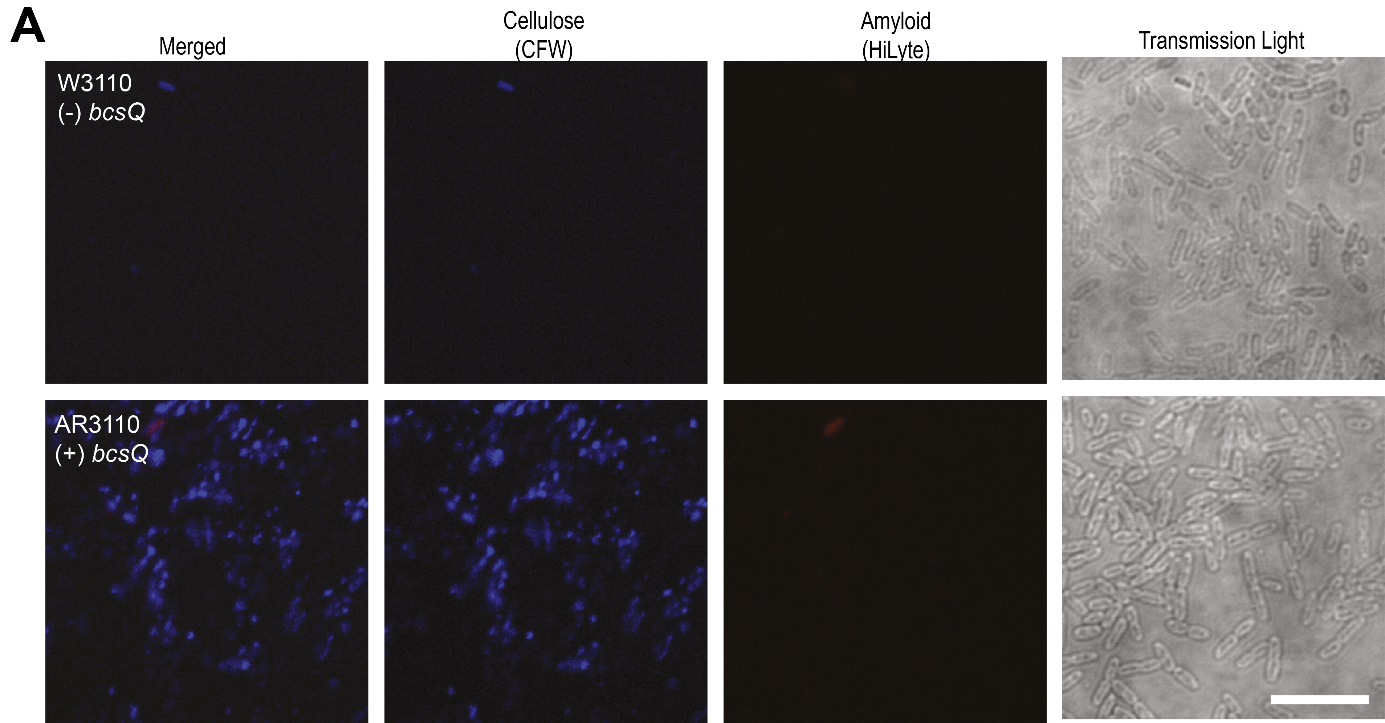

Supplementary Fig. 8 | Cellulose biosynthesis restored after bcsQ repair. (A) Fluorescence microscopy images on bcsQ-repaired AR3110 and its parental strain *E. coli* K-12 W3110 labelled with cytoplasmatic GFP and stained with CFW to visualize cellulose, and HiLyte to label amyloid structures (Serra, Richter and Hengge 2013).

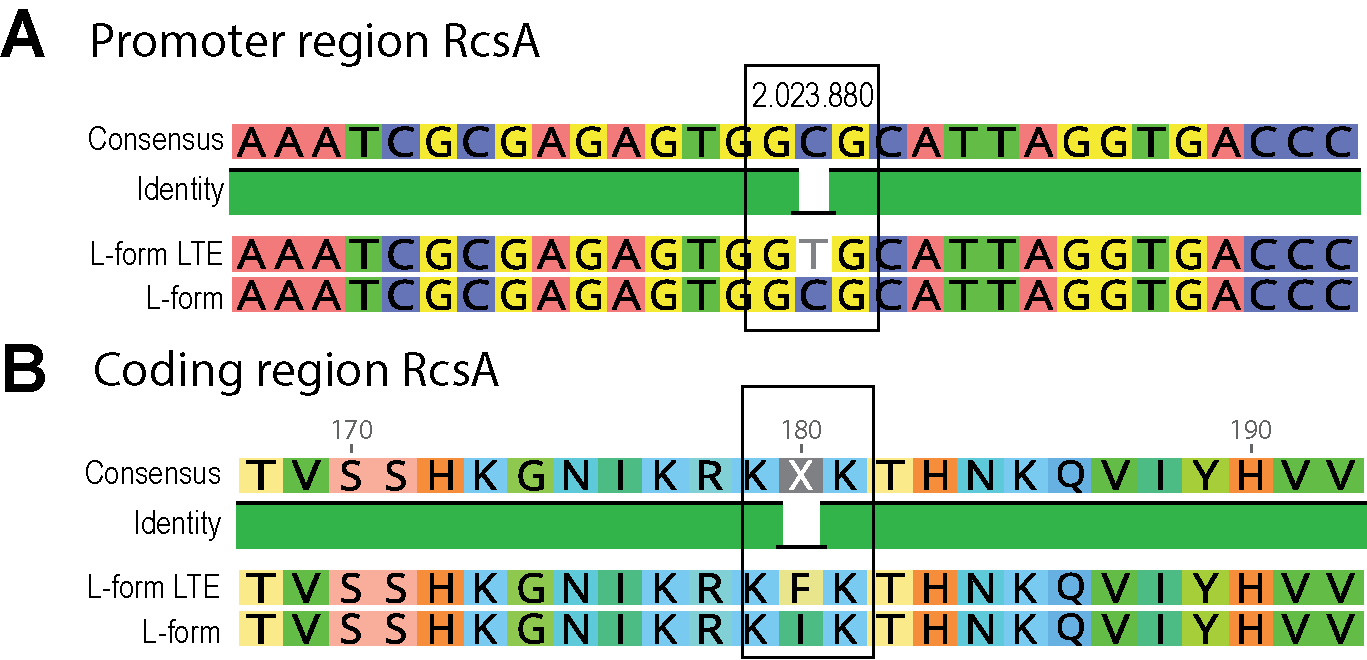

Supplementary Fig. 9 | Analysis of rcsA in L-form^LTE^. (A) The promoter region of RcsA in L-form^LTE^ compared to the original L-form^0^. The C is changed to a T. It remains unknown if the change results in expression changes. (B) The variation in the RcsA coding region at position 180. This mutations results into amino acid 180 isoleucine to change into a phenylalanine.

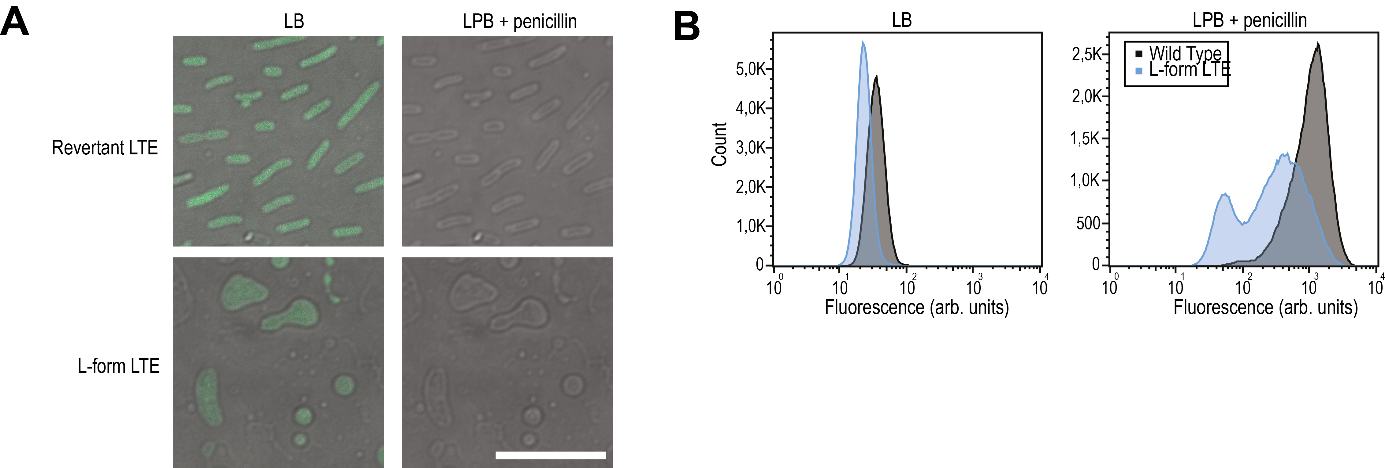

Supplementary Fig. 10 | ROS detection in L-form^LTE^. (A) Fluorescence microscopy images depict exponentially growing cells stained with CellRox Orange to detect ROS in the L-form^LTE^ strain in wall-deficient and revertant cells. Scalebar is 10µm. (B) Quantitative analysis of ROS of the L-form^LTE^ (blue) and wild type (black) in LB and LPB supplemented with penicillin using flow cytometry (n=100,000). Wild type data is the same as Supplementary Fig. 1 for comparison. Fluorescence is shown in arbitrary units. A summary of the data can be found in Supplementary Table 6.

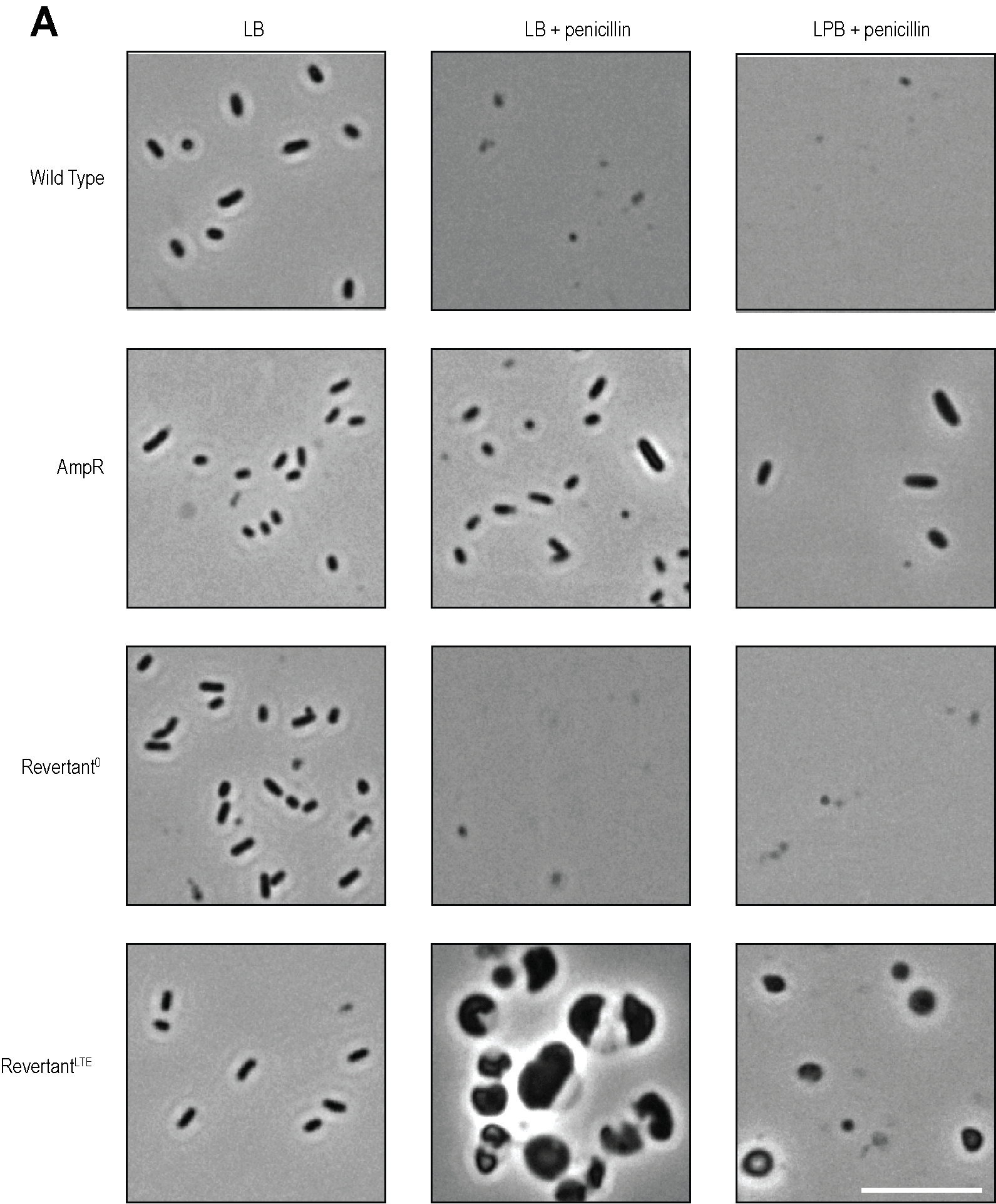

Supplementary Fig. 11 **| Transmission light microscopy images of Wild Type, AmpR, Revertant^0^ and Revertant^LTE^ strains after stress test.** Samples taken from the baffled flask experiment of Fig. 5F. Scalebar is 10µm.

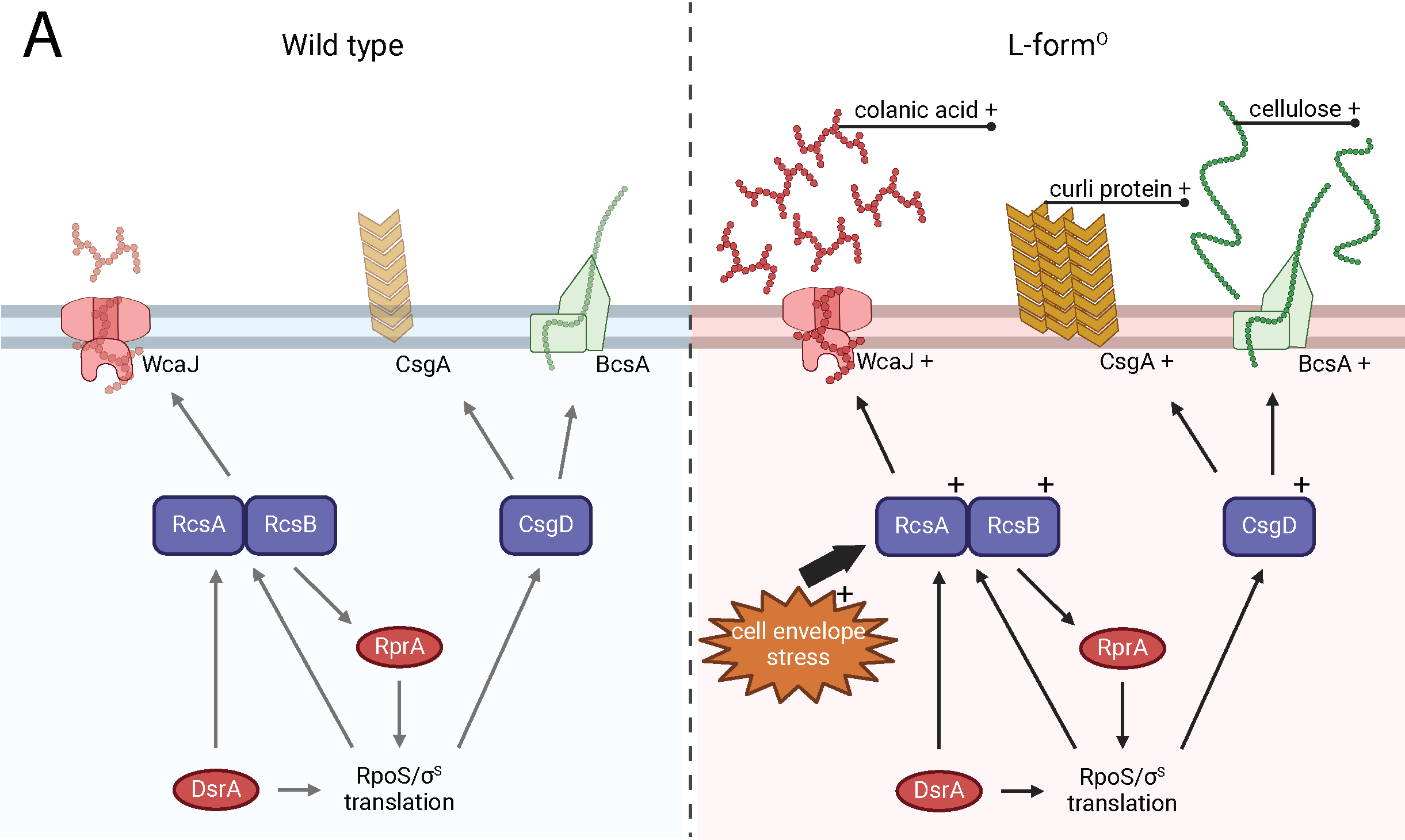

Supplementary Fig. 12 **| Proposed model of RcsA-activated cellulose and curli protein synthesis.** In wild type conditions, the small regulatory RNAs RprA and DsrA regulate RpoS translation. RpoS can activate colanic acid biosynthesis through RcsA/RcsB activation, and cellulose/curli protein synthesis through CsgD. We found that increased levels of RcsA can also activate the synthesis of cellulose and curli protein. In our model, we propose that L-forms due to high exposure to cell envelope stress, have upregulated synthesis of these three extracellular matrix components through activation of RprA. Illustration was created using Biorender.

### Supplementary Tables

| Supplementary Table 1 \| Cell Size descriptive statistics. A Kruskal–Wallis test with Dunn’s post hoc test used to compare the average cell size of L-form^0^ with the wild type and spheroplasts in Prism. ^1^One outlier removed using ROUT (Motulsky and Brown 2006). | | | |
| --- | --- | --- | --- |
| **Strain** | **Mean** | **SD** | **N** |
| Wild Type | 3.036 | 0.68 | 60 |
| Spheroplasts | 11.85^****^ | 2.94 | 60 |
| L-form^0^ | 2.740^ns^ | 0.80 | 59^1^ |

Supplementary Table 2 | ROS detection of L-form^0^ using flow cytometry. The cells were stained with the CellROX dye. The data was not normally distributed according to D'Agostino & Pearson test (P value <0.0001). A Kruskal–Wallis test with Dunn’s *post hoc* test used to compare the L-form^0^ with the wild type grown in the same treatment. The wild type is the strain contains SerW-GFP.

| **Sample** | **Treatment** | **Geometric Mean** | **GSD** | **5^th^ perc.** | **95^th^ perc.** | | **Count** |
| --- | --- | --- | --- | --- | --- | --- | --- |
| Wild Type | LB | 35.4 | 1.3 | 22.05 | | 57.2 | 99,355 |
|  | LPB + penicillin | 968 | 1.9 | 285.1 | | 2359 | 100,000 |
| L-form^0^ | LB | 27.6**** | 1.5 | 16.9 | | 54.0 | 100,042 |
|  | LPB + penicillin | 467**** | 3.4 | 45.0 | | 1960 | 100,000 |

#### Supplementary Table 3 | RNA sequencing data of *ndh, rcsA, yjbH, ykgH, csgD,* and *nhaR*.

|  | **Wild Type** | | | **Spheroplasts** | | | **L-form^0^** | | | **Fold Change** | |
| --- | --- | --- | --- | --- | --- | --- | --- | --- | --- | --- | --- |
|  | Mean | SD | N | Mean | SD | N | Mean | SD | N | Wild Type vs L-form^0^ | Spheroplast vs L-form^0^ |
| [*ndh*](https://www.uniprot.org/uniprotkb/P00393/) | 60.87 | 4.42 | 3 | 208.43 | 18.81 | 3 | 201.36 | 139.02 | 3 | 3.31 | 0.97 |
| [*rcsA*](https://www.uniprot.org/uniprotkb/P0DMC9/) | 1.31 | 0.49 | 3 | 223.58 | 15.82 | 3 | 1187.55 | 604.37 | 3 | 906.5 | 5.31 |
| [*yjbH*](https://www.uniprot.org/uniprotkb/P32689/) | 8.06 | 0.41 | 3 | 290.93 | 10.19 | 3 | 251.46 | 46.32 | 3 | 31.19 | 0.86 |
| [*ykgH*](https://www.uniprot.org/uniprotkb/P77180/) | 0.05 | 0.08 | 3 | 1.30 | 0.22 | 3 | 3.45 | 3.15 | 3 | 73.86 | 2.65 |
| [*csgD*](https://www.uniprot.org/uniprotkb/P52106/) | 0.33 | 0.11 | 3 | 1.34 | 0.52 | 3 | 10.38 | 6.79 | 3 | 31.45 | 7.75 |
| [*nhaR*](https://www.uniprot.org/uniprotkb/P0A9G2/) | 62.98 | 4.21 | 3 | 128.4 | 8.84 | 3 | 76.09 | 4.55 | 3 | 1.21 | 0.59 |

Supplementary Table 4 | List of SNPs found in the RcsA region of the L-form^LTE^ strain. Position reflects the location on the genome. Type and Change specifies the changes. Protein reveals the name of the protein. Effect in protein shows the direct change in the protein. Freq % reveals the frequency of the SNP in all total reads.

| **Protein** | **Position** | **Type** | **Change** | **Effect in Protein** | **Freq (%)** |
| --- | --- | --- | --- | --- | --- |
| rcsA | 2024506 | SNP (transversion) | T-A | Ile180Phe | 100 |
| rcsA | 2023880 | SNP (transition) | T-C |  | 100 |

Supplementary Table 5 | Crystal Violet biofilm biomass assay. A Kruskal–Wallis test with Dunn’s *post hoc* test used to compare the L-form^LTE^ against L-form^0^ and spheroplasts in Prism. ^1^ two outliers removed using ROUT. ^2^ two samples where above the detectable threshold.

| **Strain** | **Mean** | **SD** | **N** |
| --- | --- | --- | --- |
| Spheroplasts | 3.334* | 0.52 | 62^1^ |
| L-form^0^ | 2.907 | 0.83 | 64 |
| L-form^LTE^ | 4.658**** | 1.13 | 62^2^ |

Supplementary Table 6 | ROS detection of L-form^LTE^ using flow cytometry. Data not normally distributed according to D'Agostino & Pearson test (P value <0.0001). A Kruskal–Wallis test with Dunn’s *post hoc* test used to compare the L-form^LTE^ with the wild type (data from supplementary Table 2) in Prism.

| **Sample** | **Treatment** | **Geometric Mean** | **GSD** | **5^th^ perc.** | **95^th^ perc.** | **Count** |
| --- | --- | --- | --- | --- | --- | --- |
| L-form^LTE^ | LB | 23.6**** | 1.3 | 15.5 | 36.2 | 100,000 |
|  | LPB + penicillin | 263**** | 3.1 | 39.3 | 1343 | 100,000 |

#### Supplementary Table 7 | Bacterial strains used in the paper.

| **Strain** | **Genotype** | **Source** |
| --- | --- | --- |
| U66-GFP | K-12 MG1655. F-. λ-. ilvG-. rfb-50. rph-1. promoterless::gfpmut2 | (Zaslaver, Bren et al. 2006). CGSC 6300 |
| serW-GFP | K-12 MG1655. F-. λ-. ilvG.- rfb-50. rph-1. serW::gfpmut2 | (Zaslaver, Bren et al. 2006). CGSC 6300 |
| L-form^0^ | serW-GFP mutant | XXX |
| L-form^LTE^ | Long-term evolution mutant | XXX |
| ΔrcsA | K-12 BW25113. F-. Δ(araD-araB)567. ΔlacZ4787(::rrnB-3). λ-. ΔrcsA726::kan. rph-1. Δ(rhaD-rhaB)568. hsdR514 | KEIO (Baba, Ara et al. 2006) |
| ΔrcsB | K-12 BW25113. F-. Δ(araD-araB)567. ΔlacZ4787(::rrnB-3). λ-. ΔrcsB770::kan. rph-1. Δ(rhaD-rhaB)568. hsdR514 | KEIO (Baba, Ara et al. 2006) |
| ΔrcsC | K-12 BW25113. F-. Δ(araD-araB)567. ΔlacZ4787(::rrnB-3). λ-. ΔrcsC771::kan. rph-1. Δ(rhaD-rhaB)568. hsdR514 | KEIO (Baba, Ara et al. 2006) |
| ΔrcsD | K-12 BW25113. F-. Δ(araD-araB)567. ΔlacZ4787(::rrnB-3). λ-. ΔrcsD769::kan. rph-1. Δ(rhaD-rhaB)568. hsdR514 | KEIO (Baba, Ara et al. 2006) |
| ΔrcsF | K-12 BW25113. F-. Δ(araD-araB)567. ΔrcsF721::kan. ΔlacZ4787(::rrnB-3). λ-. rph-1. Δ(rhaD-rhaB)568. hsdR514 | KEIO (Baba, Ara et al. 2006) |
| W3110 | K-12 F^-^ . lambda^-^. IN(rrnD-rrnE)1. rph-1 | (Serra, Richter and Hengge 2013) |
| AR3110 | W3110 with restored bcsQ | (Serra, Richter and Hengge 2013) |
| rcsA | K-12 W3110. pCA24N::*rcsA* | ASKA (-) (Kitagawa, Ara et al. 2005) |
| AmpR | K-12 W1485. F–. mcrA Δ(mrr-hsdRMS-mcrBC) φ80lacZΔM15. ΔlacX74. recA1. araD139. Δ(ara-leu)7697. galU. galK. λ–rpsL(StrR) endA1. nupG. pET-16b | Novagen™ |

| Supplementary Table 8 \| Primers used in this study. | |  |
| --- | --- | --- |
| **Primer** | **Sequence (5’-3')** | **Purpose** |
| Ndh_SNP_FW | ATCTCACCCGGTAAAGTCGC | SNP confirm |
| Ndh_SNP_REV | GTCGCGCAGTTCTGCAATAG | SNP confirm |
| RcsA_SNP_FW | TCCGTAACGTTTATCATGTTATCCT | SNP confirm |
| RcsA_SNP_REV | TCGTCGAGAGATTCCGGTTT | SNP confirm |
| yjbH_SNP_FW | CATCTGACCGCCTACTGGAC | SNP confirm |
| YjbH_SNP_REV | CTGGCACCTTTCCCATGACT | SNP confirm |
| ykgH_SNP_FW | CTGACTTAACCCCGTTCCGT | SNP confirm |
| ykgH_SNP_REV | AAGGCTGGAATGGTCAGGAG | SNP confirm |
| rcsA_LTE_FW | TCCAGGATACTCCTGCAGCG | L-form^LTE^ rcsA |
| rcsA_LTE_REV | GAAGGCACAATGTACCTGGTTTCG | L-form^LTE^ rcsA |
